# Human genetic diversity alters therapeutic gene editing off-target outcomes

**DOI:** 10.1101/2021.05.20.445054

**Authors:** Samuele Cancellieri, Jing Zeng, Linda Yingqi Lin, Manuel Tognon, My Anh Nguyen, Jiecong Lin, Nicola Bombieri, Stacy A. Maitland, Marioara-Felicia Ciuculescu, Varun Katta, Shengdar Q. Tsai, Myriam Armant, Scot A. Wolfe, Rosalba Giugno, Daniel E. Bauer, Luca Pinello

**Affiliations:** Department of Computer Science, University of Verona, Verona, Italy; Division of Hematology/Oncology, Boston Children’s Hospital, Department of Pediatric Oncology, Dana-Farber Cancer Institute, Harvard Stem Cell Institute, Broad Institute, Department of Pediatrics, Harvard Medical School, Boston, Massachusetts 02115, USA; Department of Molecular, Cell and Cancer Biology, Li Weibo Institute for Rare Diseases Research, University of Massachusetts Medical School, Worcester, Massachusetts 01605, USA; TransLab, Boston Children’s Hospital, Boston, Massachusetts 02115, USA; Department of Hematology, St. Jude Children’s Research Hospital, Memphis, Tennessee 38105, USA; Molecular Pathology Unit, Center for Cancer Research, Massachusetts General Hospital, Department of Pathology, Harvard Medical School, Boston, Massachusetts 02129, USA

## Abstract

CRISPR gene editing holds great promise to modify somatic genomes to ameliorate disease. In silico prediction of homologous sites coupled with biochemical evaluation of possible genomic off-targets may predict genotoxicity risk of individual gene editing reagents. However, standard computational and biochemical methods focus on reference genomes and do not consider the impact of genetic diversity on off-target potential. Here we developed a web application called CRISPRme that explicitly and efficiently integrates human genetic variant datasets with orthogonal genomic annotations to nominate and prioritize off-target sites at scale. The method considers both single-nucleotide variants (SNVs) and indels, accounts for bona fide haplotypes, accepts spacer:protospacer mismatches and bulges, and is suitable for personal genome analyses. We tested the tool with a guide RNA (gRNA) targeting the *BCL11A* erythroid enhancer that has shown therapeutic promise in clinical trials for sickle cell disease (SCD) and β-thalassemia^1^. We find that the top candidate off-target site is produced by a non-reference allele common in African-ancestry populations (rs114518452, minor allele frequency (MAF)=4.5%) that introduces a protospacer adjacent motif (PAM) for SpCas9. We validate that SpCas9 generates indels (∼9.6% frequency) and chr2 pericentric inversions in a strictly allele-specific manner in edited CD34+ hematopoietic stem/progenitor cells (HSPCs), although a high-fidelity Cas9 variant mitigates this off-target. The CRISPRme tool highlights alternative allele-specific off-target editing as a prevalent risk of gRNAs considered for therapeutic gene editing. Our report illustrates how population and private genetic variants should be considered as modifiers of genome editing outcomes. We suggest that variant-aware off-target assessment should be considered in therapeutic genome editing efforts going forward and provide a powerful approach for comprehensive off-target nomination.

## INTRODUCTION

CRISPR genome editing extends unprecedented opportunities to develop novel therapeutics by introducing targeted genetic or epigenetic modifications to genomic regions of interest. Briefly, CRISPR offers a simple and programmable platform that couples binding to a genomic target sequence of choice with diverse effector proteins through RNA:DNA (spacer:protospacer) complementary sequence interactions mediated by a guide RNA (gRNA) spacer sequence matching a genomic protospacer sequence restricted by protospacer adjacent motif (PAM) sequences. Editing effectors may consist of nucleases to introduce targeted double strand breaks leading to short indels and templated repairs (e.g. Cas9), deaminases for precise substitutions (base editors), or chromatin regulators for transcriptional interference or activation (CRISPRi/a) among others to achieve a range of desired biological outcomes^2^.

CRISPR-based systems may create unintended off-target modifications posing potential genotoxicity for therapeutic use. Several experimental assays and computational methods are available to uncover or forecast these off-targets^3^. Off-target sites are partially predictable based on homology to the spacer and PAM sequence. Beyond the number of mismatches or bulges, a variety of sequence features, like position of mismatch or bulge with respect to PAM or specific base changes, contribute to off-target potential^3–6^. Computational models can complement experimental approaches to off-target nomination in several respects: to triage gRNAs prior to experiments by predicting the number and cleavage potential of off-target sites and to prioritize target sites for experimental scrutiny. Genetic variants may alter protospacer and PAM sequences and therefore may influence both on-target and off-target potential. Gene editing strategies designed to specifically recognize patient mutations may increase the likelihood of editing mutant alleles, whereas variants that reduce homology to the anticipated target may decrease the efficiency of the desired genetic modification. Although a variety of in vitro and cell-based experimental methods can be used to empirically nominate off-target sites, these methods either use homology to the reference genome as a criterion to define the search space and/or use a limited set of human donor genomes to evaluate off-target potential^4, 7^. Therefore, computational methods may be especially useful to predict the impact of off-target sequences not found in reference genomes.

Prior studies considering gRNAs targeting therapeutically relevant genes using population-based variant databases like the 1000 Genomes Project (1000G) and the Exome Aggregation Consortium have highlighted how genetic variants can significantly alter the off-target landscape by creating novel and personal off-target sites not present in a single reference genome^8, 9^. Although these prior studies provide code to reproduce analyses, implementation choices make these tools not suitable to analyze large variant datasets and to consider higher numbers of mismatches. In addition, these methods ignore bulges between RNA:DNA hybrids, cannot efficiently model alternative haplotypes and indels, and require extensive computational skills to utilize.

Several user-friendly websites have been developed to aid the design of gRNAs and to assess their potential off-targets^10–13^. Even though variant-aware prediction is an important problem for genome editing interventions, these scalable graphical user interface (GUI) based tools do not account for genetic variants. In addition, these tools artificially limit the number of mismatches for the search and/or do not support DNA/RNA bulges. Therefore, designing gRNAs for therapeutic intervention using current widely available tools could miss important off-target sites that may lead to unwanted genotoxicity. A complete and exhaustive off-target search with an arbitrary number of mismatches, bulges, and genetic variants that is haplotype-aware is a computationally challenging problem that requires specialized and efficient data structures.

We have recently developed a command line tool that partially solves these challenges called CRISPRitz^14^. This tool uses optimized data structures to efficiently account for single variants, mismatches and bulges but with significant limitations^14^. Here we substantially extend this work by developing CRISPRme, a tool to aid gRNA design with added support for haplotype-aware off-target enumeration, short indel variants and a flexible number of mismatches and bulges. CRISPRme is a unified, user-friendly web-based application that provides several reports to prioritize putative off-targets based on their risk in a population or individuals.

CRISPRme is flexible to accept user-defined genomic annotations, which could include empirically identified off-target sites or cell type specific chromatin features. It can integrate population genetic variants from sets of phased individual variants (like those from 1000G^15^), unphased individual variants (like those from the Human Genome Diversity Project, HGDP^16^) and population-level variants (like those from the Genome Aggregation Database, gnomAD^17^). Furthermore, it can accept personal genomes from individual subjects to identify and prioritize private off-targets due to variants specific to a single individual.

Here we demonstrate the utility of CRISPRme by analyzing the off-target potential of a gRNA currently being tested in clinical trials for SCD and β-thalassemia^1, 18, 19^. We identify possible off-targets introduced by genetic variants included within and extending beyond 1000G. We predict that the most likely off-target site, overlooked by prior analyses, is introduced by a variant common in African-ancestry individuals (rs114518452, minor allele frequency (MAF)=4.5%) and provide experimental evidence of its off-target potential in gene edited human CD34+ hematopoietic stem and progenitor cells. Furthermore, we demonstrate that alternative allele-specific off-target potential is widespread across various gRNAs and editors.

## RESULTS

CRISPRme is a web-based tool to predict off-target potential of CRISPR gene editing that accounts for genetic variation. It is available online at http://crisprme.di.univr.it. CRISPRme can also be deployed locally as a web app or used as a command line utility, both of which respect genomic privacy offline. CRISPRme takes as input a Cas protein, gRNA spacer sequence(s) and PAM, genome build, sets of variants (VCF files for populations or individuals), user-defined thresholds of mismatches and bulges, and optional user-defined genomic annotations to produce comprehensive and personalized reports (Fig. 1a**, Supplementary Notes 1-3**).

**Figure 1.**
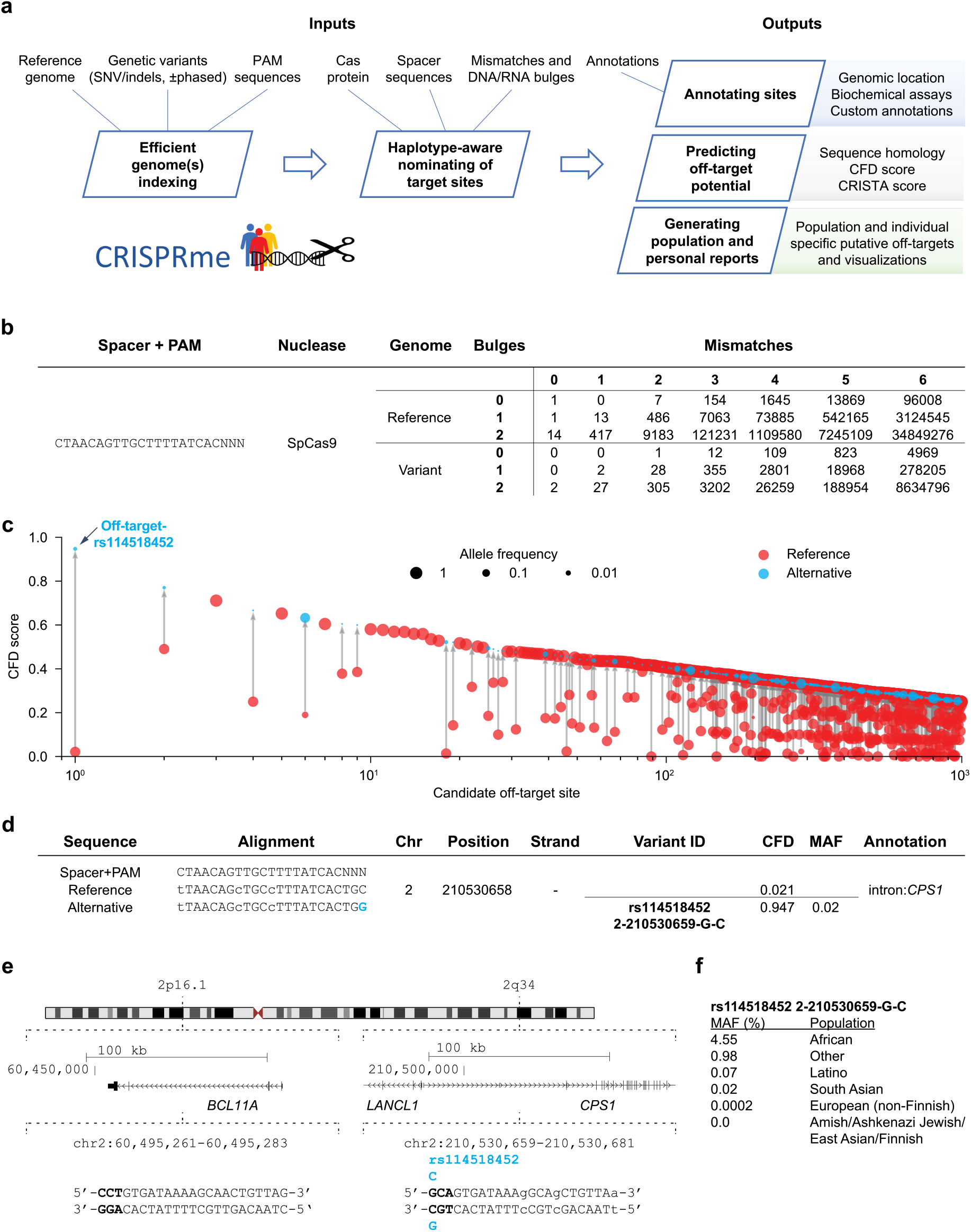
CRISPRme provides web-based analysis of CRISPR-Cas gene editing off-target potential reflecting population genetic diversity. **a)** CRISPRme software takes as input a reference genome, genetic variants, PAM sequence, Cas protein type, spacer sequence, homology threshold and genomic annotations and provides comprehensive, target-focused and individual-focused analyses of off-target potential. It is available as an online webtool and can be deployed locally or used offline as command-line software. **b)** Analysis of the *BCL11A-*1617 spacer targeting the +58 erythroid enhancer with SpCas9, NNN PAM, 1000G variants, up to 6 mismatches and up to 2 bulges. **c)** Top 1000 predicted off-target sites ranked by CFD score, indicating the CFD score of the reference and alternative allele if applicable, with allele frequency indicated by circle size. **d)** The off-target site with the highest CFD score is created by the minor allele of rs114518452. Coordinates are for hg38 and 0-start for the potential off-target and 1-start for the variant-ID. MAF is based on 1000G. **e)** The top predicted off-target site from CRISPRme is an allele-specific off-target with 3 mismatches to the *BCL11A-*1617 spacer sequence, where the rs114518452-C minor allele produces a de novo NGG PAM sequence. PAM sequence shown in bold and mismatches to *BCL11A-*1617 shown as lowercase. Coordinates are for hg38 and 1-start. **f)** rs114518452 allele frequencies based on gnomAD v3.1. Coordinates are for hg38 and 1-start. Spacer shown as DNA sequence for ease of visual alignment.

We have designed CRISPRme to be flexible with support for new gene editors with variable and extremely relaxed PAM requirements^20^. Thanks to a PAM encoding based on Aho-Corasick automata and an index based on a ternary search tree, CRISPRme can perform genome-wide exhaustive searches efficiently even with an NNN PAM, extensive mismatches (tested with up to 7) and RNA:DNA bulges (tested with up to 2) (**Supplementary Note 4**).

Notably, a comprehensive search performed with up to 6 mismatches, 2 DNA/RNA bulges and a fully non-restrictive PAM (NNN) takes only 24 hours on a small computational cluster node (Intel Xeon CPU E5-2609 v4 clocked at 2.2 GHz and 128 GB RAM). All the 1000G variants, including both SNVs and indels, can be included in the search together with all the available metadata for each individual (sex, super-population and age), and the search operation takes into account observed haplotypes (**Supplementary Note 5**). Importantly, off-target sites that represent alternative alignments to a given genomic region are merged to avoid inflating the number of reported sites. Although several tools exist to enumerate off-targets, to our knowledge only two command line tools^8, 21^ incorporate genetic variants in the search. However, they have several limitations in terms of scalability to large searches, number of mismatches, bulges, haplotypes, and variant file formats supported and do not provide an easy-to-use graphical user interface (**Supplementary Note 6**).

CRISPRme generates several reports (**Supplementary Note 2**). First, it summarizes for each gRNA all the potential off-targets found in the reference or variant genomes based on their mismatches and bulges (Fig. 1b) and generates a file with detailed information on each of these candidate off-targets (**Supplementary File 1,** Supplementary Table 1). Second, it compares gRNAs to customizable annotations. By default, it classifies possible off-target sites based on GENCODE^22^ (genomic features) and ENCODE^23^ (candidate cis-regulatory elements, cCREs) annotations. It can also incorporate user-defined annotations in BED format, such as empiric off-target scores or cell type specific chromatin features (Supplementary Figure 1, **Supplementary Note 5**). Third, using 1000G and/or HGDP^16^ variants, CRISPRme reports the cumulative distribution of homologous sites based on the reference genome or super-population. These global reports could be used to compare a set of gRNAs based on how genetic variation impacts their predicted on-and off-target cleavage potential using cutting frequency determination (CFD) or CRISPR Target Assessment (CRISTA)^24^ scores (Supplementary Figure 2). CRISPRme includes multiple scoring metrics and can be easily extended with new ones, including scores tailored for different editors. Finally, CRISPRme can generate personal genome focused reports called *personal risk cards* (**Supplementary Note 3**). These reports highlight private off-target sites due to unique genetic variants.

We tested CRISPRme with a gRNA (#1617) targeting a GATA1 binding motif at the +58 erythroid enhancer of *BCL11A*^18, 19^. A recent clinical report described two patients, one with SCD and one with β-thalassemia, each treated with autologous gene modified hematopoietic stem and progenitor cells (HSPCs) edited with Cas9 and this gRNA, who showed sustained increases in fetal hemoglobin, transfusion-independence and absence of vaso-occlusive episodes (in the SCD patient) following therapy. This study as well as prior pre-clinical studies with the same gRNA (#1617) did not reveal evidence of off-target editing in treated cells when considering off-target sites nominated by bioinformatic analysis of the human reference genome and empiric analysis of in vitro genomic cleavage potential (Supplementary Table 2, **Supplementary Note 7**)^1, 19, 25^. CRISPRme analysis found that the predicted off-target site with both the greatest CFD score and the greatest increase in CFD score from the reference to alternative allele was at an intronic sequence of *CPS1* (Fig. 1c,d), a genomic target subject to common genetic variation (modified by a SNP with MAF ≥ 1%). CFD scores range from 0 to 1, where the on-target site has a score of 1. The alternative allele rs114518452-C generates a TGG PAM sequence (that is, the optimal PAM for SpCas9) for a potential off-target site with 3 mismatches and a CFD score (CFD_alt_ 0.95) approaching that of the on-target site (Fig. 1e). In contrast, the reference allele rs114518452-G disrupts the PAM to TGC, which markedly reduces predicted cleavage potential (CFD_ref_ 0.02). rs114518452-C has an overall MAF of 1.33% in gnomAD v3.1, with MAF of 4.55% in African/African-American, 0.9% in Other, 0.07% in Latino/Admixed American, 0.02% in European (non-Finnish) and 0.00% in East Asian populations (Fig. 1f, Supplementary Table 3).

To consider the off-target potential that could be introduced by personal genetic variation that would not be predicted by 1000G variants, we analyzed HGDP variants identified from whole genome sequences of 929 individuals from 54 diverse human populations. We observed 249 candidate off-targets for gRNA #1617 with CFD ≥0.2 for which the CFD score in HGDP exceeded that found for either the reference genome or 1000G variants by at least 0.1 (Fig. 2a, Supplementary Figure 3). These additional variant off-targets not found from 1000G were observed in each super-population, with the greatest frequency in the African super-population (Fig. 2b). 229 (92.0%) of these variant off-targets were unique to a super-population and 172 (69.1%) of these were private to just one individual (Fig. 2c). Furthermore, single individual focused searches, for example an analysis of HGDP01211, an individual of the Oroqen population within the East Asian super-population, showed that most variant off-targets (with higher CFD score than reference) were due to variants also found in 1000G (n=32369, 90.4%), a subset were due to variants shared with other individuals from HGDP but absent from 1000G (n=3177, 8.9%), and a small fraction were private to the individual (n=234, 0.7%) (Fig. 2d). Among these private off-targets was one generated by a variant (rs1191022522, 3-99137613-A-G, gnomAD v3.1 MAF 0.0053%) where the alternative allele produces a canonical NGG PAM that increases the CFD score from 0.14 to 0.54 (Fig. 2d,e).

**Figure 2.**
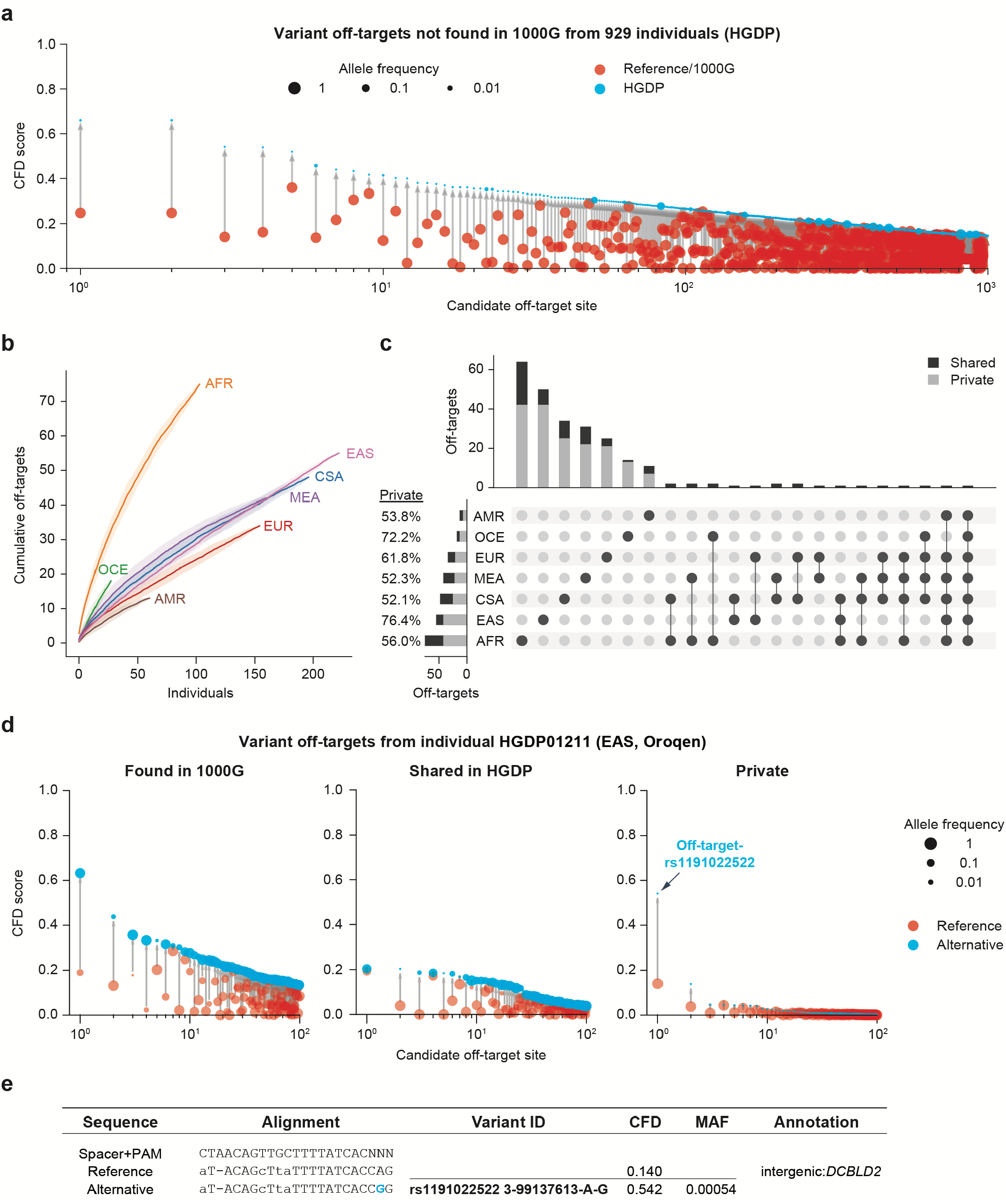
CRISPRme provides analysis of off-target potential of CRISPR-Cas gene editing reflecting population and private genetic diversity. **a)** CRISPRme analysis was conducted with variants from HGDP comprising whole genome sequencing of 929 individuals from 54 diverse human populations. HGDP variant off-targets with greater CFD scores than the reference genome or 1000G were plotted and sorted by CFD score, with HGDP variant off-targets shown in blue and reference or 1000G variant off-targets shown in red. **b)** HGDP variant off-targets with CFD≥0.2 and increase in CFD of ≥0.1. Individual samples from each of the seven super-populations were shuffled 100 times to calculate the mean and 95% confidence interval. **c)** Intersection analysis of HGDP variant off-targets with CFD≥0.2 and increase in CFD of ≥0.1. Shared variants were found in 2 or more HGDP samples while private variants were limited to a single sample. **d)** CRISPRme analysis of a single individual (HGDP01211) showing the top 100 variant off-targets from each of the following three categories: shared with 1000G variant off-targets (left panel), higher CFD score compared to reference genome and 1000G but shared with other HGDP individuals (center panel), and higher CFD score compared to reference genome and 1000G with variant not found in other HGDP individuals (right panel). For the center and right panels, reference refers to CFD score from reference genome or 1000G variants. **e)** The top predicted private off-target site from HGDP01211 is an allele-specific off-target where the rs1191022522-G minor allele produces a canonical NGG PAM sequence in place of a noncanonical NAG PAM sequence. Spacer shown as DNA sequence for ease of visual alignment.

To experimentally test the top predicted off-target from CRISPRme, we identified a CD34+ HSPC donor of African ancestry heterozygous for rs114518452-C, the variant predicted to introduce the greatest increase in off-target cleavage potential (Fig. 1c-f). We performed RNP electroporation using a gene editing protocol that preserves engrafting HSC function. Amplicon sequencing analysis showed 92.0 ± 0.5% indels at the on-target site and 4.8 ± 0.5% indels at the off-target site. For reads spanning the variant position, indels were strictly found at the alternative PAM-creation allele without indels observed at the reference allele (Fig. 3a-c), suggesting 9.6 ± 1.0% off-target editing of the alternative allele. In an additional 6 HSPC donors homozygous for the reference allele rs114518452-G/G, 0.00 ± 0.00% indels were observed at the off-target site, suggesting strict restriction of off-target editing to the alternative allele (Fig. 3d).

**Figure 3.**
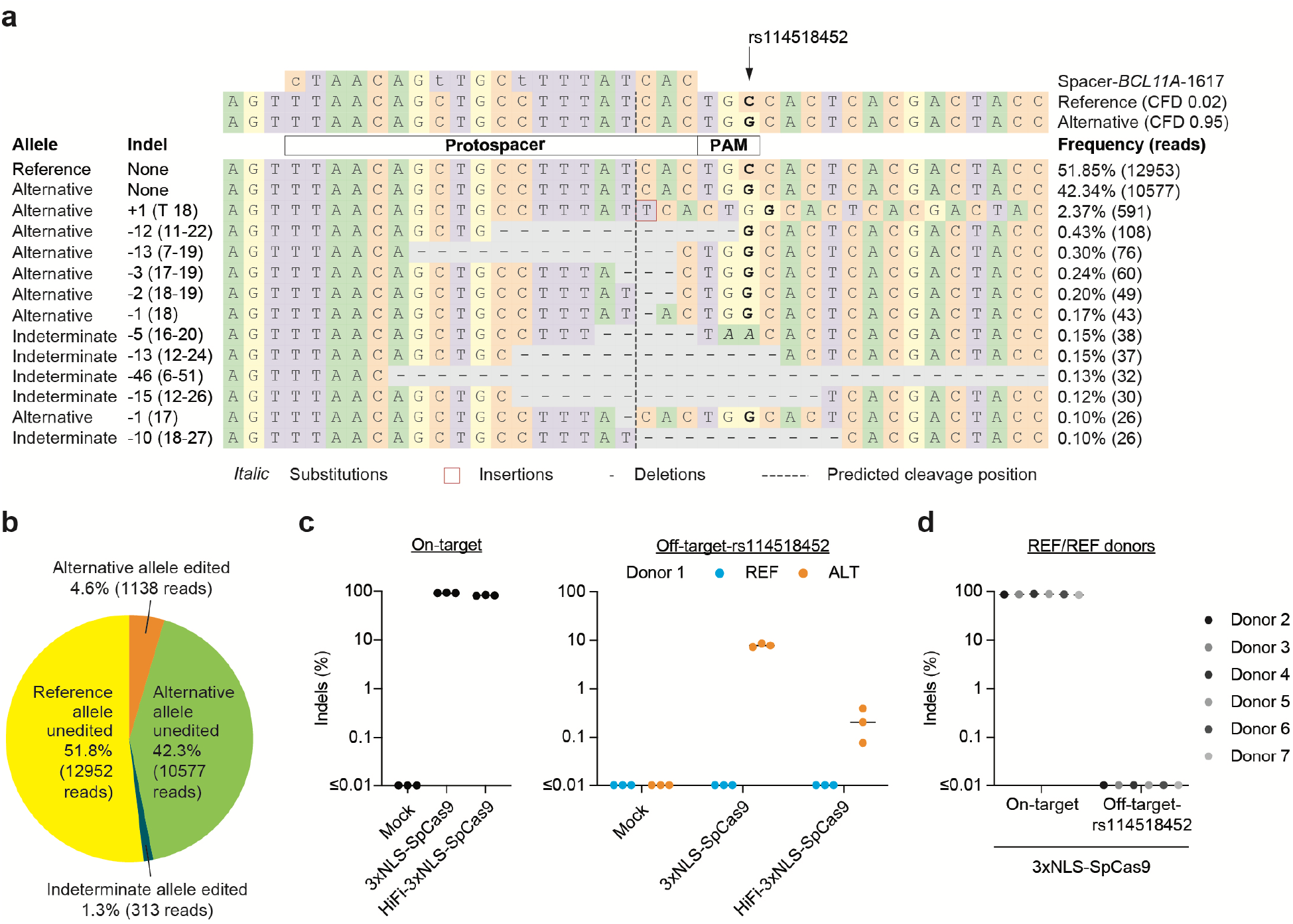
Allele-specific off-target editing by a *BCL11A* enhancer targeting gRNA in clinical trials associated with a common variant in African-ancestry populations. **a)** Human CD34+ HSPCs from a donor heterozygous for rs114518452-G/C (Donor 1, REF/ALT) were subject to 3xNLS-SpCas9:sg1617 RNP electroporation followed by amplicon sequencing of the off-target site around chr2:210,530,659-210,530,681 (off-target-rs114518452 in 1-start hg38 coordinates). CFD scores for the reference and alternative alleles are indicated and representative aligned reads are shown. Spacer shown as DNA sequence for ease of visual alignment, with mismatches indicated by lowercase and the rs114518452 position shown in bold. **b)** Reads classified based on allele (indeterminate if the rs114518452 position is deleted) and presence or absence of indels (edits). **c)** Human CD34+ HSPCs from a donor heterozygous for rs114518452-G/C (Donor 1) were subject to 3xNLS-SpCas9:sg1617 RNP electroporation, HiFi-3xNLS-SpCas9:sg1617 RNP electroporation, or no electroporation (mock) followed by amplicon sequencing of the on-target and off-target-rs114518452 sites. Each dot represents an independent biological replicate (*n* = 3). Indel frequency was quantified for reads aligning to either the reference or alternative allele. **d)** Human CD34+ HSPCs from 6 donors homozygous for rs114518452-G/G (Donors 2-7, REF/REF) were subject to 3xNLS-SpCas9:sg1617 RNP electroporation with 1 biological replicate per donor followed by amplicon sequencing of the on-target and off-target-rs114518452 sites.

The on-target *BCL11A* intronic enhancer site is on chr2p while the off-target-rs114518452 site is on chr2q within an intron of a non-canonical transcript of *CPS1*. Inversion PCR demonstrated inversion junctions consistent with the presence of ∼150 Mb pericentric inversions between *BCL11A* and the off-target site only in edited HSPCs carrying the alternative allele (Fig. 4a,b). Deep sequencing of the inversion junction showed that inversions were restricted to the alternative allele in the heterozygous cells (Fig. 4c,d). Droplet digital PCR revealed these inversions to be present at 0.16 ± 0.04% allele frequency (Fig. 4e).

**Figure 4.**
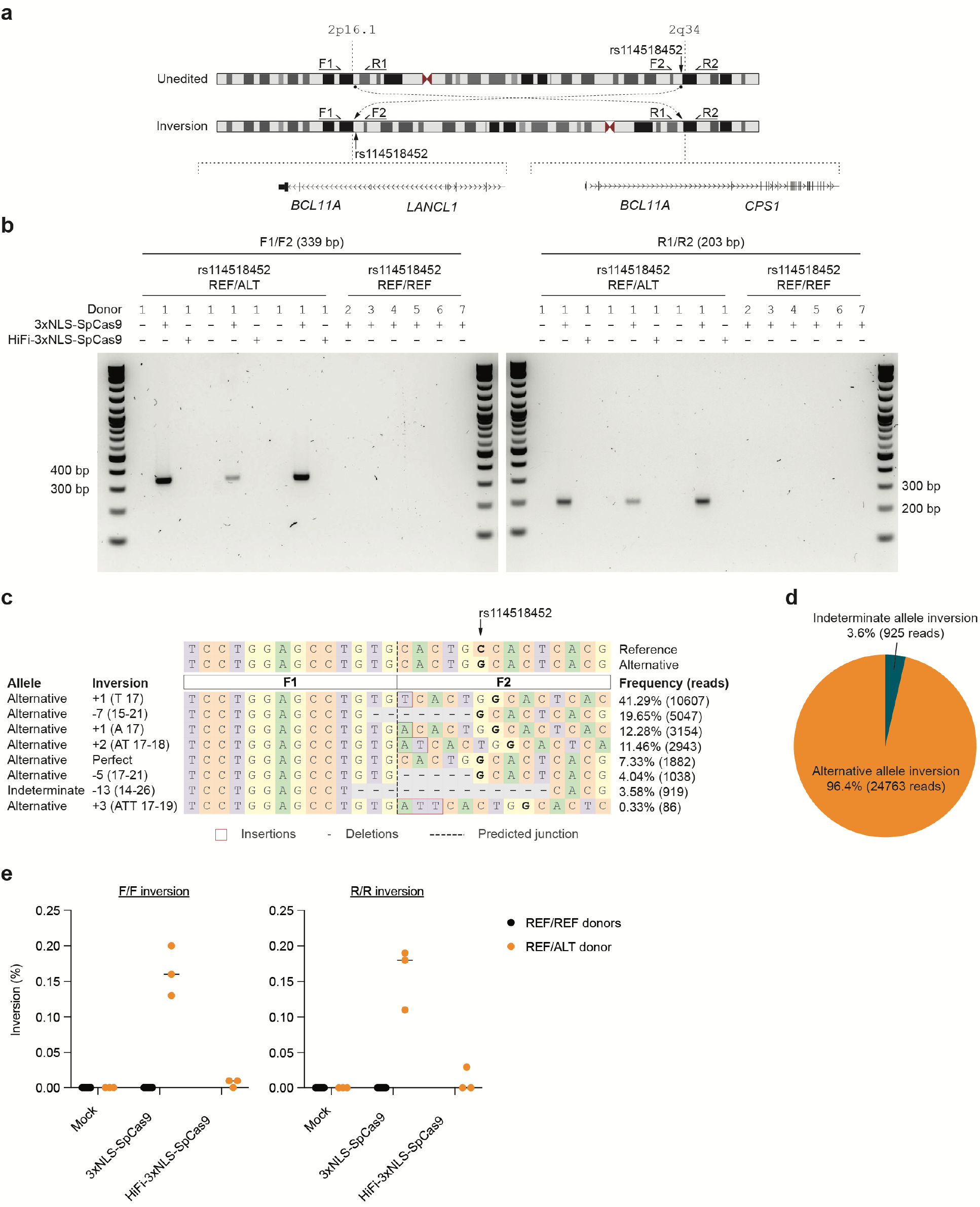
Allele-specific pericentric inversion following *BCL11A* enhancer editing due to off-target cleavage. **a)** Concurrent cleavage of the on-target and off-target-rs114518452 sites could lead to pericentric inversion of chr2 as depicted. PCR primers F1, R1, F2, and R2 were designed to detect potential inversions. **b)** Human CD34+ HSPCs from a donor heterozygous for rs114518452-G/C (Donor 1) were subject to 3xNLS-SpCas9:sg1617 RNP electroporation, HiFi-3xNLS-SpCas9:sg1617 RNP electroporation, or no electroporation with 3 biological replicates. Human CD34+ HSPCs from 6 donors homozygous for rs114518452-G/G (Donors 2-7, REF/REF) were subject to 3xNLS-SpCas9:sg1617 RNP electroporation with 1 biological replicate per donor. Gel electrophoresis for inversion PCR was performed with F1/F2 and R1/R2 primer pairs on left and right respectively with expected sizes of precise inversion PCR products indicated. **c)** Reads from amplicon sequencing of the F1/F2 product (expected to include the rs114518452 position) from 3xNLS-SpCas9:sg1617 RNP treatment were aligned to reference and alternative inversion templates. The rs114518452 position is shown in bold. **d)** Reads classified based on allele (indeterminate if the rs114518452 position deleted). **e)** Inversion frequency by ddPCR from same samples as in (**b**). F/F indicates forward and R/R reverse inversion junctions as depicted in (**a**).

Various high-fidelity Cas9 variants may improve the specificity of gene editing, although at the possible cost of reduced efficiency^26^. Gene editing following the same electroporation protocol using a HiFi variant 3xNLS-SpCas9 (R691A)^27^ in heterozygous cells revealed 82.3 ± 1.6% on-target indels with only 0.1 ± 0.1% indels at the rs114518452-C off-target site, i.e. a ∼48-fold reduction compared to SpCas9 (Fig. 3c). Inversions were not detected following HiFi-3xNLS-SpCas9 editing (Fig. 4b,e).

To examine the pervasiveness of alternative allele off-target potential, we evaluated an additional 13 gRNAs in clinical development or otherwise widely used for SpCas9-based nuclease or base editing^28–37^ and 6 gRNAs for non-SpCas9-based editing such as for SaCas9 and Cas12a^33, 38–41^ (Supplementary Table 4**, Supplementary Files 2-3, Supplementary Note 8**). CRISPRme analysis including the 1000G and HGDP genetic variant datasets showed 18% (95% confidence interval 13-23%) of the total nominated off-targets were due to alternative allele-specific off-targets. Most alternative allele-specific off-targets were associated with rare variants (MAF <1%), although candidate off-targets associated with common variants were identified for each gRNA (Fig. 5a). None of these alternative allele-specific off-target sites were described in the original manuscripts reporting the editing strategies and off-target analyses.

**Figure 5.**
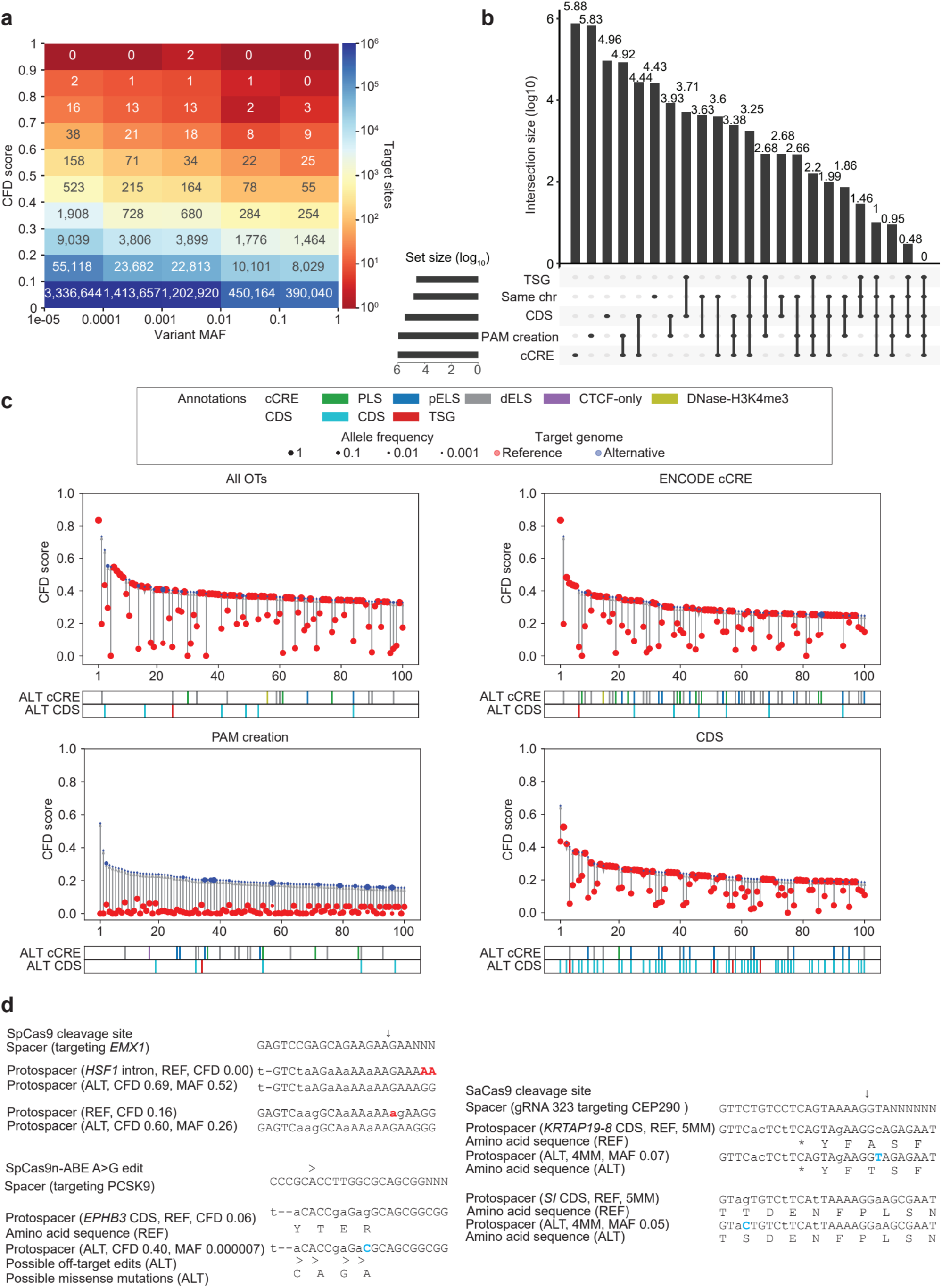
CRISPRme illustrates prevalent off-target potential due to genetic variation. **a)** Heatmap showing the distribution of alternative allele nominated off-targets for SpCas9 guides by CFD score and MAF. **b)** UpSet plot showing overlapping annotation categories for candidate off-targets (TSG, tumor suppressor gene; candidate off-targets on the same chromosome as the on-target; CDS regions; cCRE from ENCODE and PAM creation events). **c)** Top 100 predicted off-target sites ranked by CFD score for the gRNA targeting *PCSK9* with no filter, found in cCREs, corresponding to PAM creation events, and in CDS regions) **d)** *Top left:* Candidate off-target sites with increased predicted cleavage potential introduced by common (MAF 52% and 26%) indel variants for a SpCas9 gRNA targeting *EMX1. Right:* Candidate off-target cleavage sites within coding sequences with increased homology to a lead gRNA for SaCas9 targeting of *CEP290* to treat congenital blindness in current clinical trials due to common SNPs. *Bottom:* Potential missense mutations in the *EPHB3* tumor suppressor resulting from candidate off-target A-to-G base editing by a preclinical lead gRNA targeting *PCSK9* to reduce LDL cholesterol levels. Deletions shown in red, SNPs shown in blue.

CRISPRme produces visualizations to specifically highlight alternative allele-specific candidate off-target sites overlapping candidate cis-regulatory elements and protein coding sequences (including putative tumor suppressor genes^42^) and/or which involve PAM creation events (Fig. 5b-c, Supplementary Figure 4). For example, within the top 20 candidate off-targets nominated by CRISPRme for a SpCas9 gRNA targeting *EMX1*^35^, two sites involve genetic variants with high MAF (52% and 26%) and are associated with substantial increases in CFD score from REF to ALT (+0.69 and +0.44). The first is an intronic PAM creation variant, while the second introduces two PAM-proximal matches to the gRNA (Fig. 5d). Notably, both of these candidate off-targets involve indel variants, underscoring the utility of CRISPRme to account for variants beyond SNPs.

In addition to visualizing candidate off-target sites by predictive score rank (such as CFD or CRISTA) for SpCas9 derived editors, CRISPRme can also visualize candidate off-targets by number of mismatches and bulges, which may be especially useful for Cas proteins with distinct PAMs for which predictive scores are not readily available. For example, SaCas9 is a clinically relevant nuclease whose small size favors packaging to AAV. For a SaCas9-associated gRNA targeting *CEP290*^40^ currently being evaluated in clinical trials to treat a form of congenital blindness (NCT03872479), CRISPRme nominated two candidate off-targets associated with common SNPs (MAF 7% and 5%) that reduced mismatches from 5 (REF) to 4 (ALT) which are predicted to produce cleavages within coding sequences (Fig. 5d).

CRISPRme can nominate variant off-targets for base editors and evaluate their base editing susceptibility within a user-defined editing window. For a gRNA targeting *PCSK9*^37^ that has been used with SpCas9-nickase adenine base editor in vivo in preclinical studies to reduce LDL cholesterol levels, 4 of the top 5 candidate off-target sites involve alternative alleles, including one with CFD_ref_ 0.2 and CFD_alt_ 0.75 found in an ENCODE candidate enhancer element. CRISPRme nominated a candidate off-target associated with a rare variant (MAF 0.0007%) that increased the CFD score from 0.06 (REF) to 0.40 (ALT) which would be predicted to produce missense mutations in *EPHB3*, a putative tumor suppressor gene (Fig. 5d).

The underlying computational challenge that CRISPRme addresses extends beyond CRISPR-based applications to other technologies based on nucleic acid sequence recognition. For example, CRISPRme can nominate off-targets for RNA-targeting strategies, whether RNA-guided gene editors or even oligonucleotide sequences used as RNA interference (RNAi) or antisense oligo (ASO) therapies (Supplementary Figure 5). We performed a variant-aware search (without PAM restriction) for the FDA-approved antisense oligonucleotide Nusinersen^43, 44^, which targets *SMN2* pre-mRNA to treat spinal muscular atrophy. Using CRISPRme, we identified a potential off-target site within a coding region wherein a common SNP (MAF 2%) reduces the number of mismatches from 3 (REF) to 2 (ALT). Similarly, analysis of the FDA-approved RNAi therapy Inclisiran^45^, which targets *PCSK9* mRNA to treat hypercholesterolemia, revealed that its antisense strand has a candidate off-target in the 3’ UTR of the ribosomal gene *RPP14* for which a common insertion variant (MAF 36%) reduces the number of mismatches and bulges from 7 (REF) to 4 (ALT).

These results demonstrate how personal genetic variation may influence the off-target potential of sequence-based therapies like genome editing. Increased availability of haplotype-resolved genomes of diverse ancestry would enhance ability to nominate variant-associated off-target sites present in human populations. The practical implications of allele-specific off-target editing need to be considered on a case-by-case basis (also see **Supplementary Note 7**). In the case of *BCL11A* enhancer editing, up to ∼10% of SCD patients with African ancestry would be expected to carry at least one rs114518452-C allele, leading to ∼10% cleavage at an off-target site that was not identified in prior studies of this gRNA using currently available tools (Supplementary Table 2). Our results highlight that allele-specific off-target editing potential is not equally distributed across all ancestral groups, but especially concentrated in those of African ancestry. Therefore, gene editing efforts that include subjects of African ancestry (like those targeting sickle cell disease) might pay particular attention to this issue. At the same time, our analysis shows that allele-specific off-targets can be private to an individual, so all humans could be susceptible to such effects. Of note, as is true for off-target genetic changes in general, the mere possibility of somatic genetic alteration does not imply functional consequence. Now that we have developed CRISPRme to enable scalable, variant-aware off-target nomination, a challenge is that depending on the prevalence of a variant, it may be difficult to obtain primary cells of relevant genotype to perform biological validation in a relevant therapeutic context. However, one limitation of current tools including CRISPRme is the fact that potential off-targets cannot be enumerated based on structural variants or other complex genetic events such as combinations of indels, SNP. Future extensions of CRISPRme based on new data structures such as graph genomes^46, 47^ could enable these complex searches and also improve their efficiency.

We recommend several steps to minimize risk of unintended allele-specific off-target effects during therapeutic genome editing, consistent with regulatory guidance to consider effects of genetic variation^48^. First, prioritize use of genome editing methods, such as high-fidelity editors and pulse delivery, that maximize specificity. Second, nominate off-targets in a variant-aware manner, with particular attention toward genetic variants found in relevant patient populations. Third, employ off-target detection assays that are variant-aware to empirically evaluate the likelihood of off-target editing (see **Supplementary Note 7**). Fourth, perform a risk assessment of variant off-target editing given predicted genomic annotations, mechanisms of DNA repair, delivery to target cells and disease context. Fifth, if excess allele-specific genome editing risks are identified, consider including genotype among the subject inclusion/exclusion criteria. Finally, for therapeutic genome editing indications in which it is feasible (such as hematopoietic cell targeting), prospectively monitor somatic modifications in patient samples to gather information about the frequency and consequence of such events.

CRISPRme offers a simple-to-use tool to comprehensively evaluate off-target potential across diverse populations and within individuals. CRISPRme is available at http://crisprme.di.univr.it and may also be deployed locally to preserve privacy (**Supplementary Note 9**).

## Data and software availability

CRISPRme source code is available at https://github.com/pinellolab/crisprme and the webapp is online at http://crisprme.di.univr.it. The version of CRISPRme (1.8.8) used to generate the results presented in this manuscript has been deposited on Zenodo: https://doi.org/10.5281/zenodo.5047489. Sequencing data is deposited in the NCBI Sequence Read Archive database under accession number PRJNA733110.

## Acknowledgement

L.P. received support from the U.S. NIH (R35 HG010717 and RM1 HG009490). D.E.B. was supported by the National Heart, Lung, and Blood Institute (OT2HL154984, P01HL053749), Burroughs Wellcome Fund and the St. Jude Children’s Research Hospital Collaborative Research Consortium. R.G received support from European Union’s ERA-NET JPCOFUND2 (JPND2019-466-037). We thank Stuart H. Orkin, Guillaume Lettre, J. Keith Joung, Vikram Pattanayak, Karl Petri, Anne H. Shen, Elia Dirupo and Francesco Masillo for helpful input.

## Author contributions

S.C., L.Y.L., M.T., N.B., R.G., and L.P. created the software, J.Z., M.A.N., S.A.M., M.F.C., V.K., S.Q.T., M.A., S.A.W., D.E.B. designed and conducted experiments, S.C., J.Z., L.Y.L., R.G., D.E.B., and L.P. performed data analysis, S.C., R.G., D.E.B., and L.P. conceived the work, S.C., J.Z., L.Y.L, R.G., D.E.B., and L.P. wrote the paper with input from all authors.

## Competing financial interests statement

L.P. has financial interests in Edilytics, Inc., Excelsior Genomics, and SeQure Dx, Inc. L.P.’s interests were reviewed and are managed by Massachusetts General Hospital and Partners HealthCare in accordance with their conflict of interest policies.

## Materials & correspondence

Please direct requests to R.G., D.E.B., and L.P.

## Supplementary Information

**Supplementary Figure 1.**
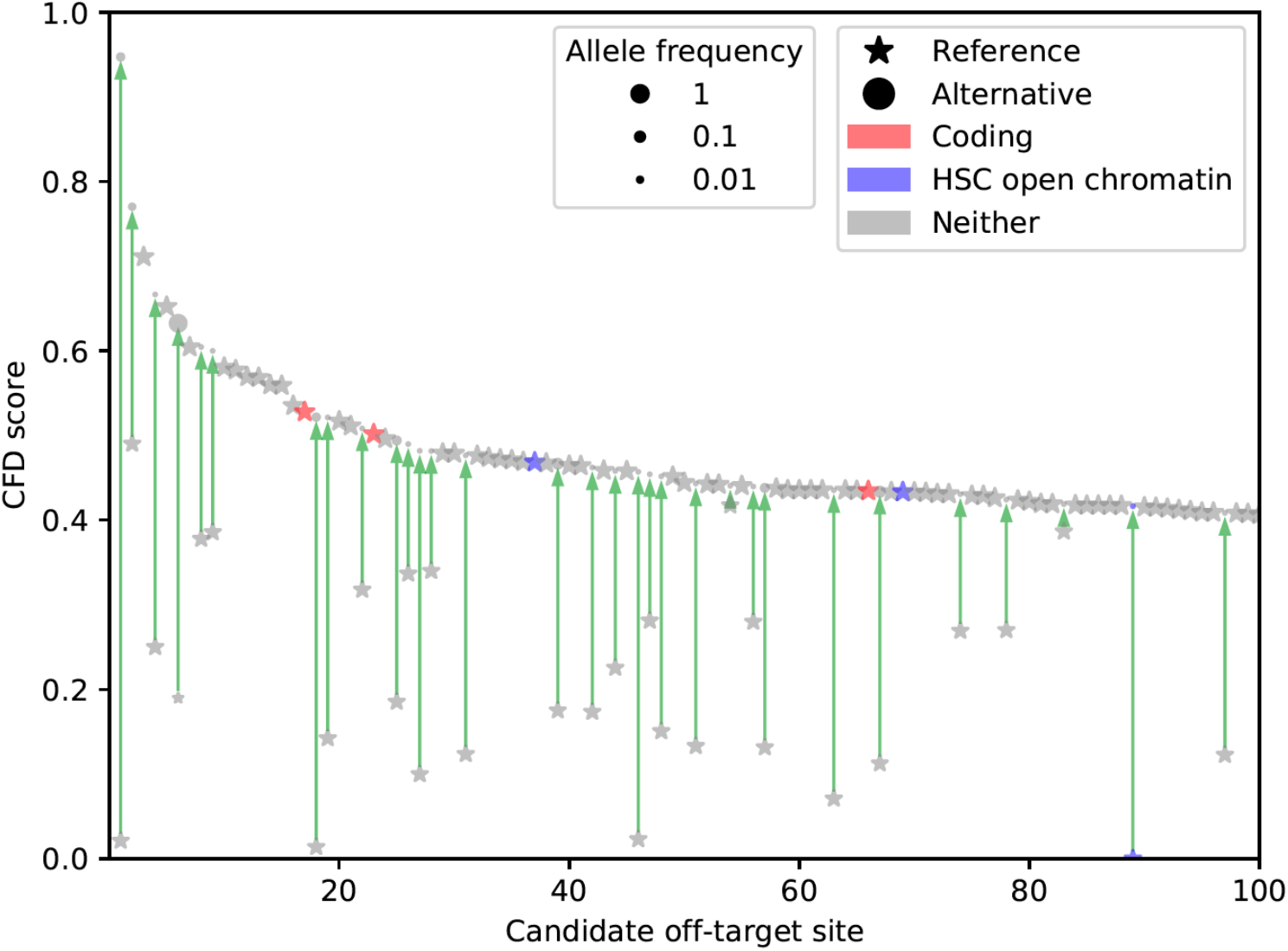
Top 100 predicted off-target sites for *BCL11A*-1617 spacer by CFD score. CRISPRme search as in Fig. 1. Candidate off-target sites within coding regions based on GENCODE annotations and ATAC-seq peaks in HSCs based on user-provided annotations (data from Corces *et al.* 2016) are highlighted.

**Supplementary Figure 2.**
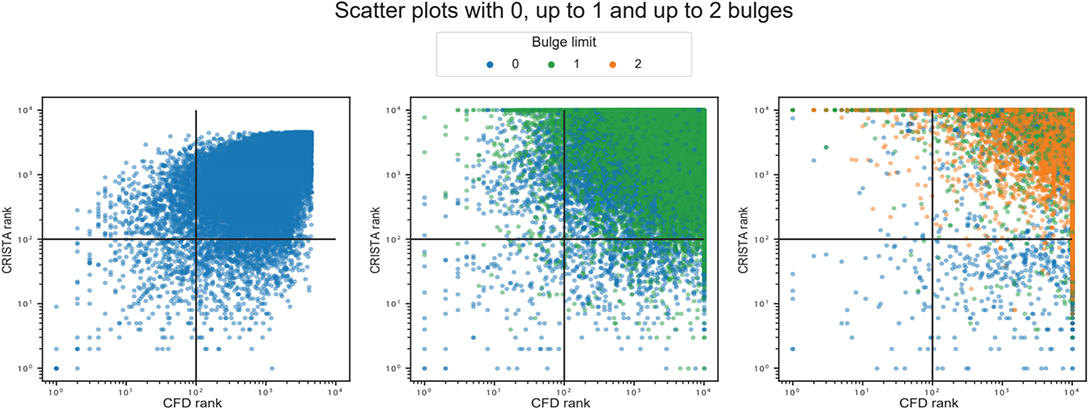
Plots with rank ordered correlation between CFD and CRISTA reported targets. Scatter plots show from left to right, the correlation of ranked targets, extracted by selecting top 10000 targets ordered by CFD and CRISTA score, respectively. The left plot shows the rank correlation of targets with 0 bulges (Pearson’s correlation: 0.57, Spearman’s correlation: 0.55), the center plot shows rank correlation of targets with 1 bulge (Pearson’s correlation: −0.16, Spearman’s correlation: −0.33) and the right plot shows the rank correlation of targets with 2 bulges (Pearson’s correlation: −0.55, Spearman’s correlation: - 0.80). All the correlations have p-values below 10^!”^. The colors represent the lowest count of bulges for each target, since the two scoring methods may prioritize different alignments and thus different number of mismatches and bulges of the same genomic target.

**Supplementary Figure 3.**
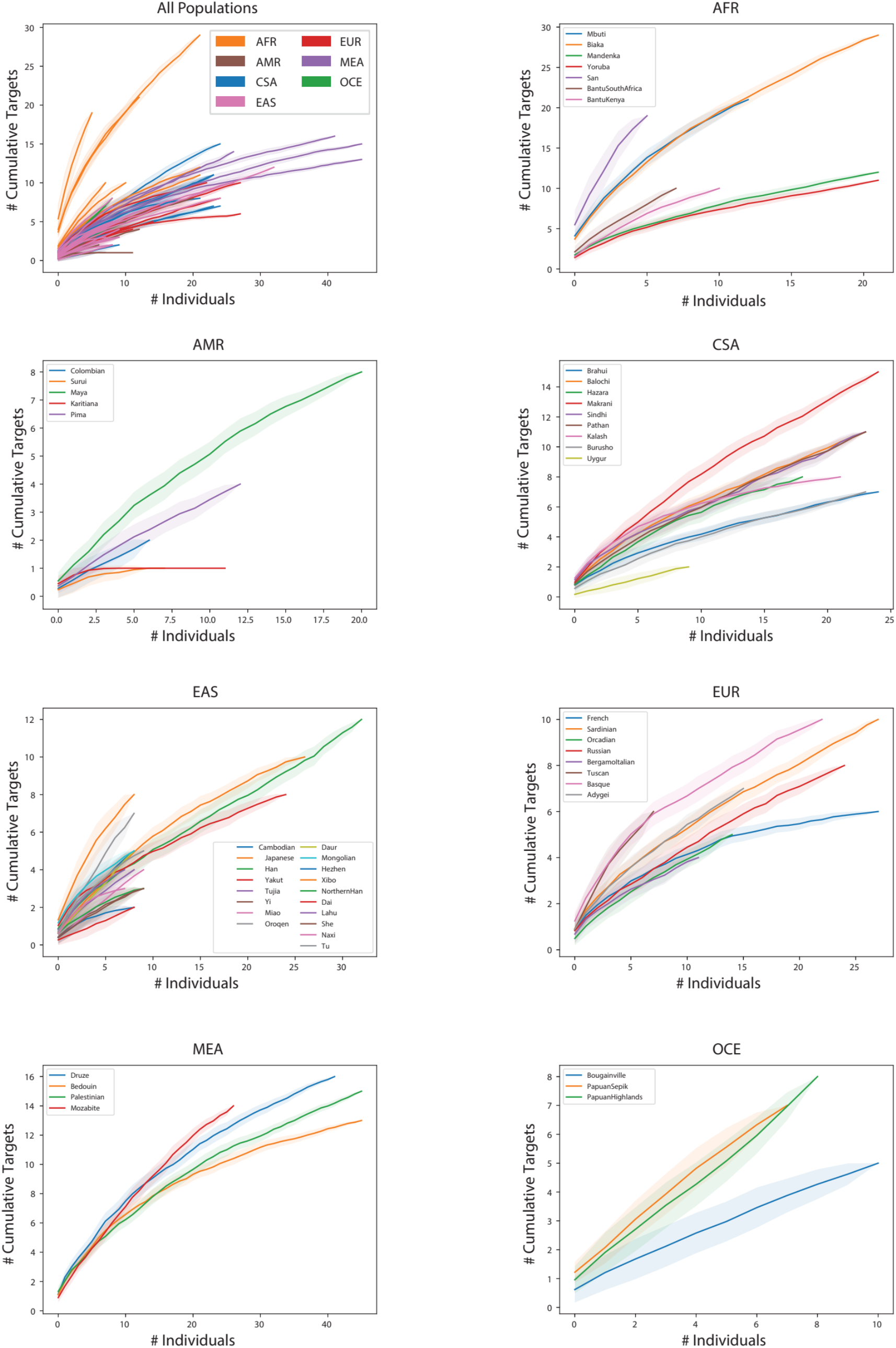
HGDP super-population distribution plots. HGDP variant off-targets with CFD≥0.2 and increase in CFD of ≥0.1. Individual samples from each of the seven super-populations were shuffled 100 times to calculate the mean and 95% confidence interval. First panel shows distribution within all 54 discrete populations, colored by super-population. Additional seven panels show distribution of discrete populations within each listed super-population.

**Supplementary Figure 4.**
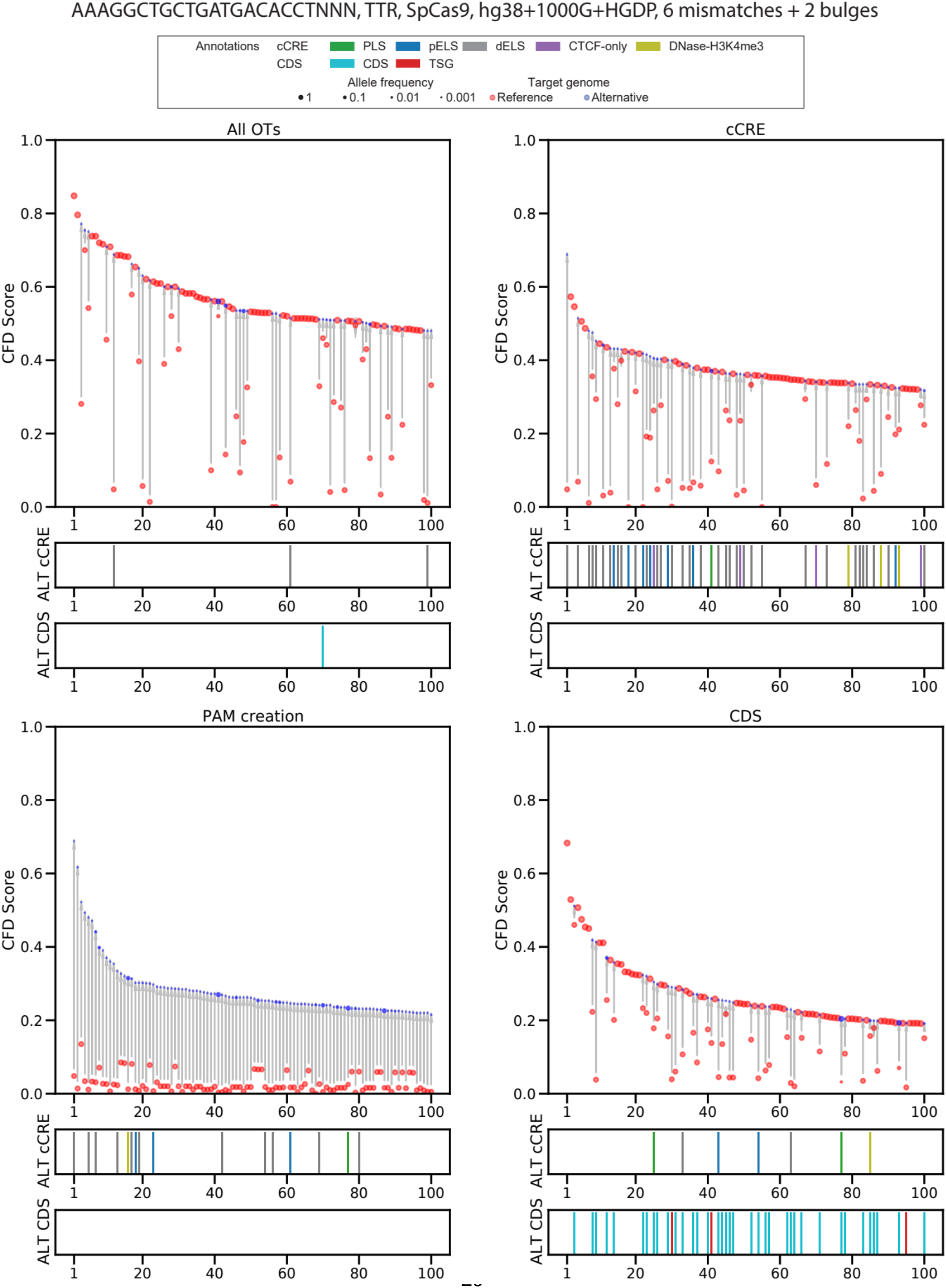
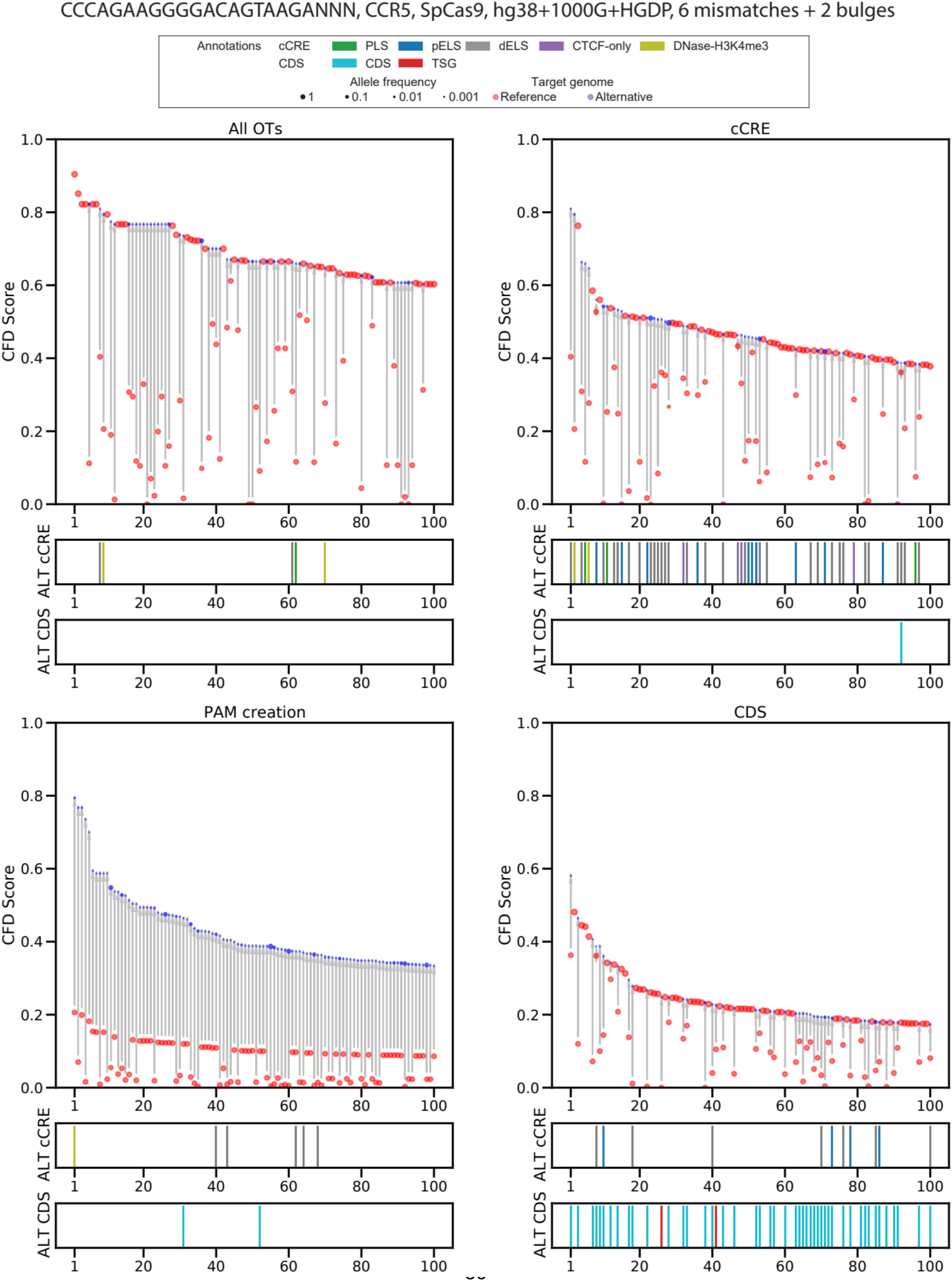
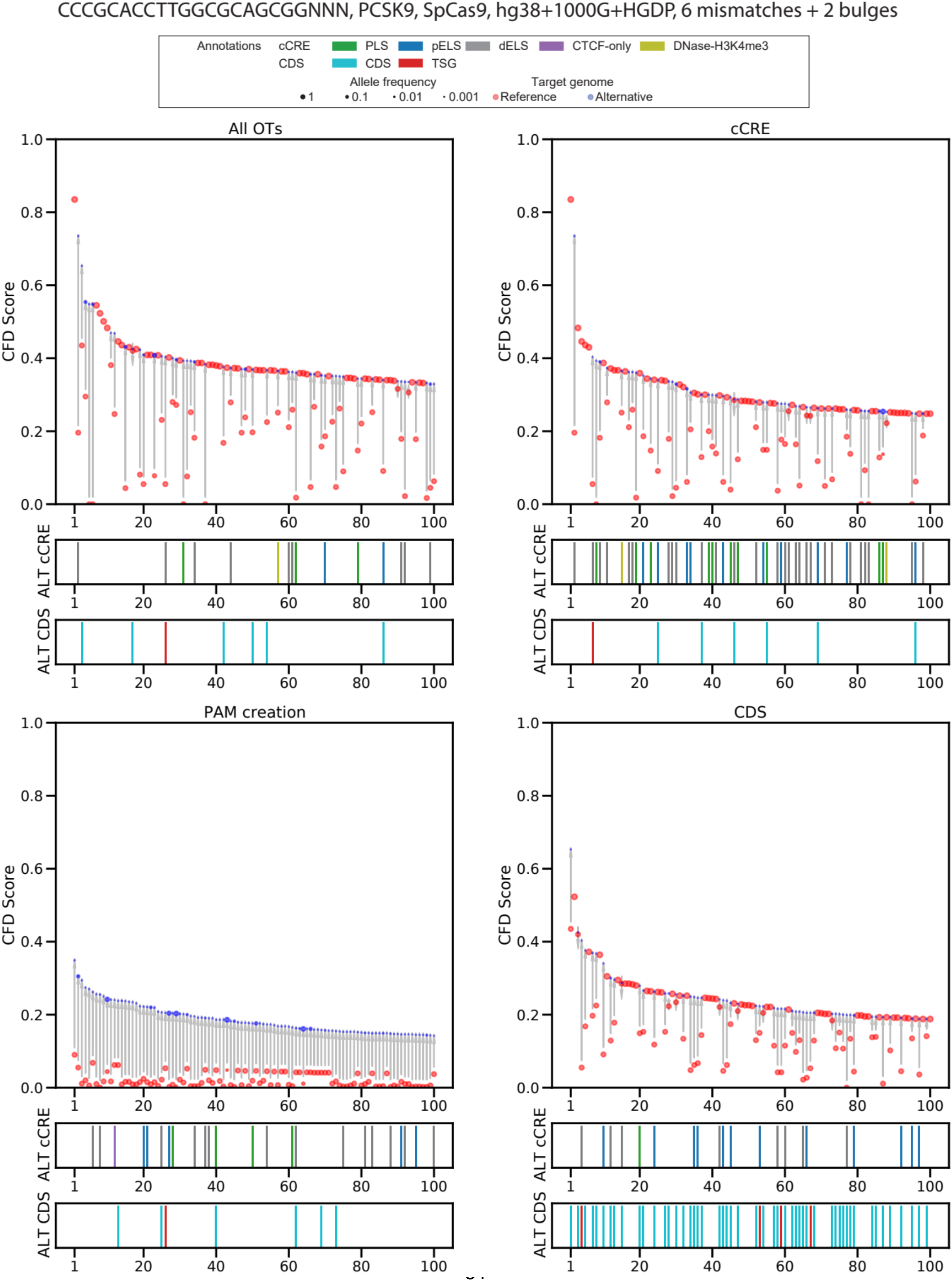
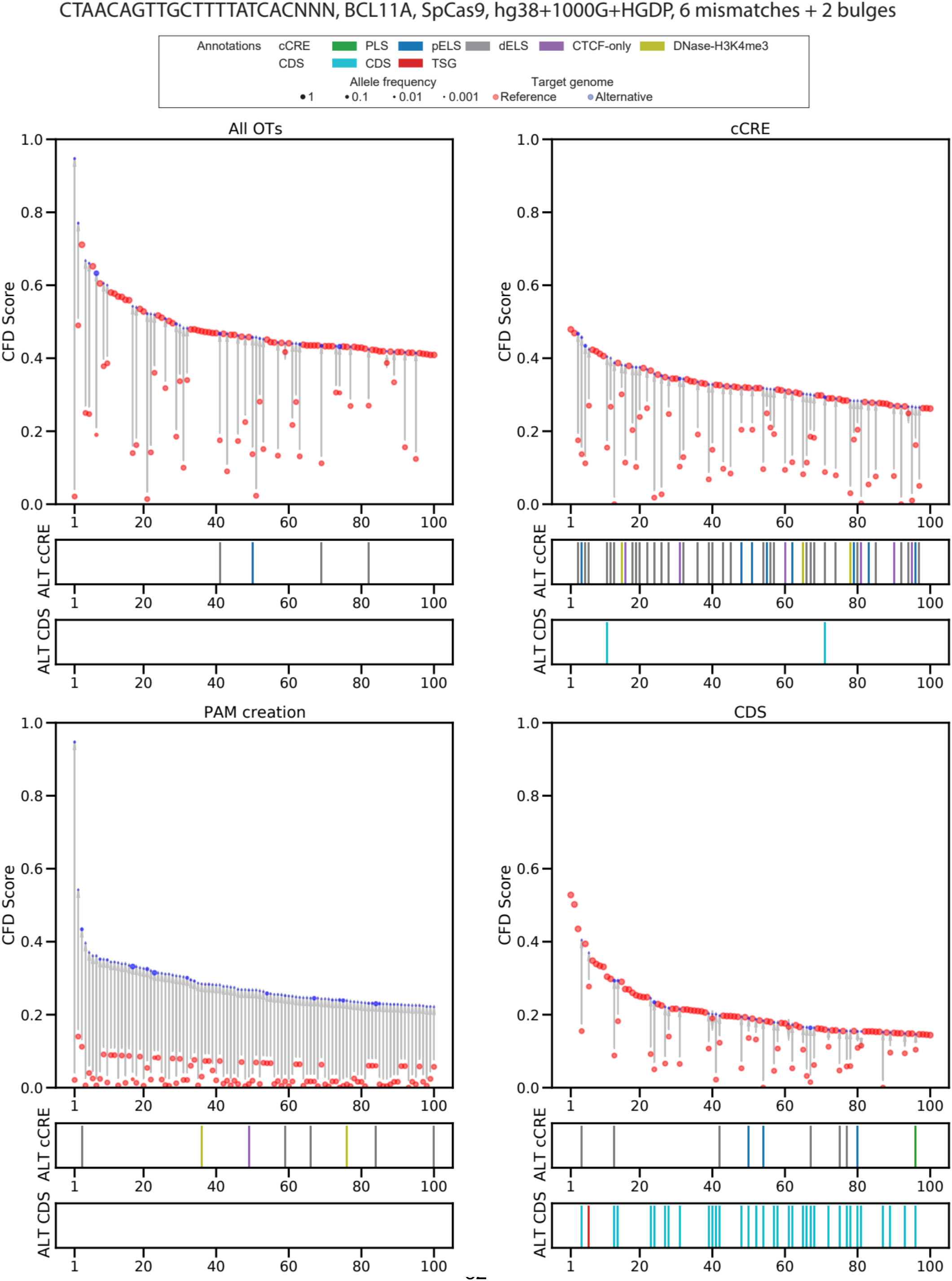
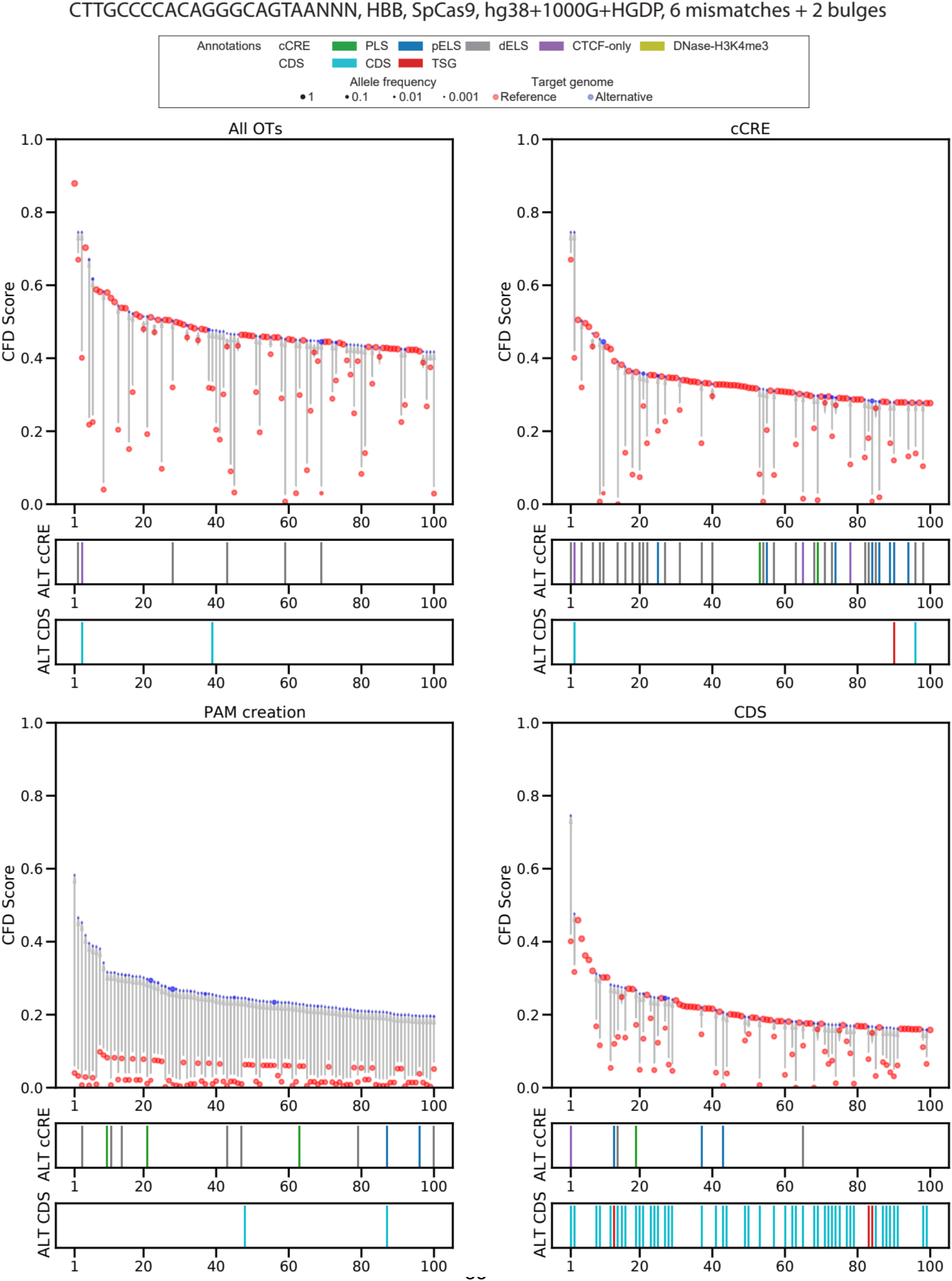
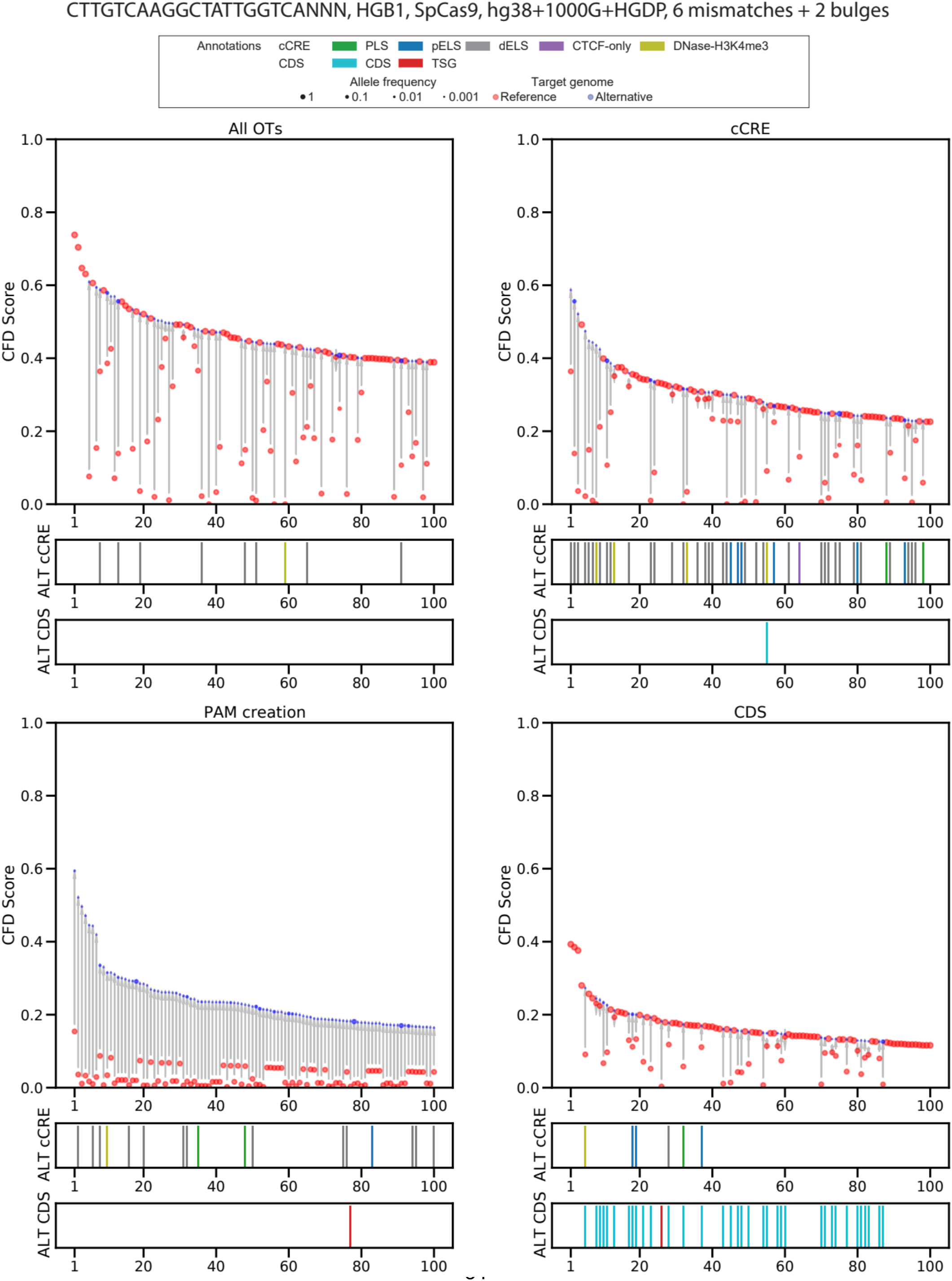
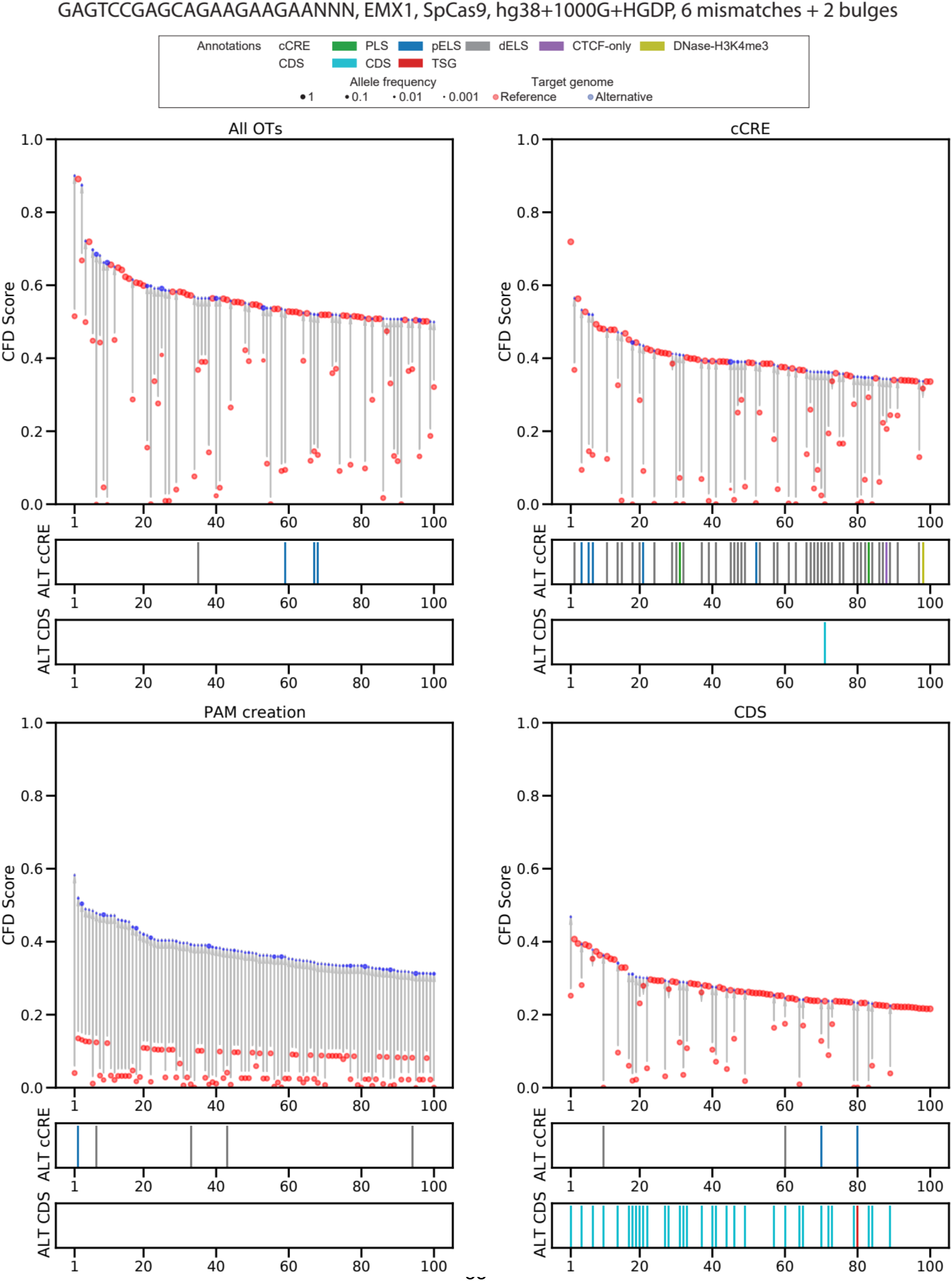
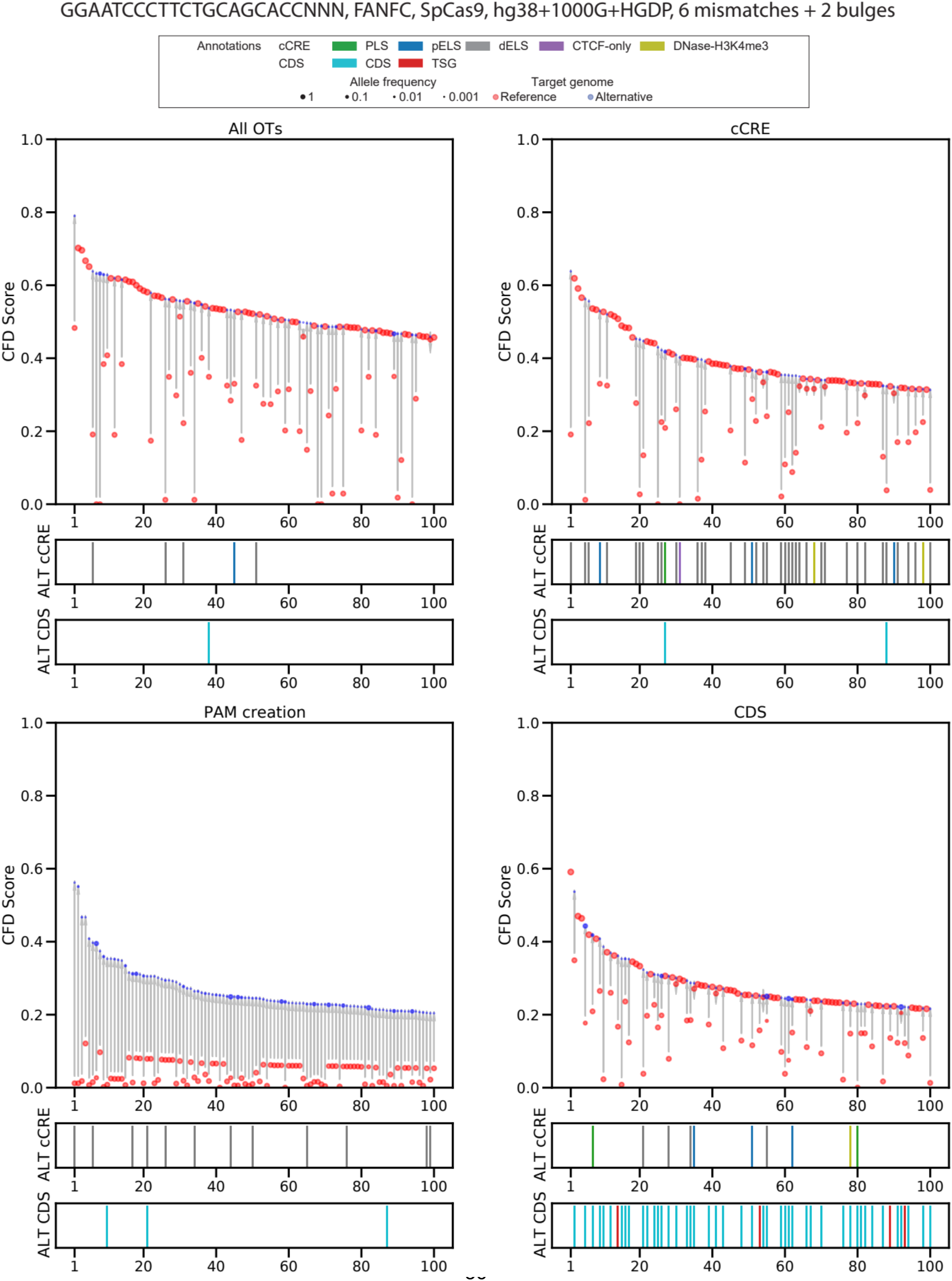
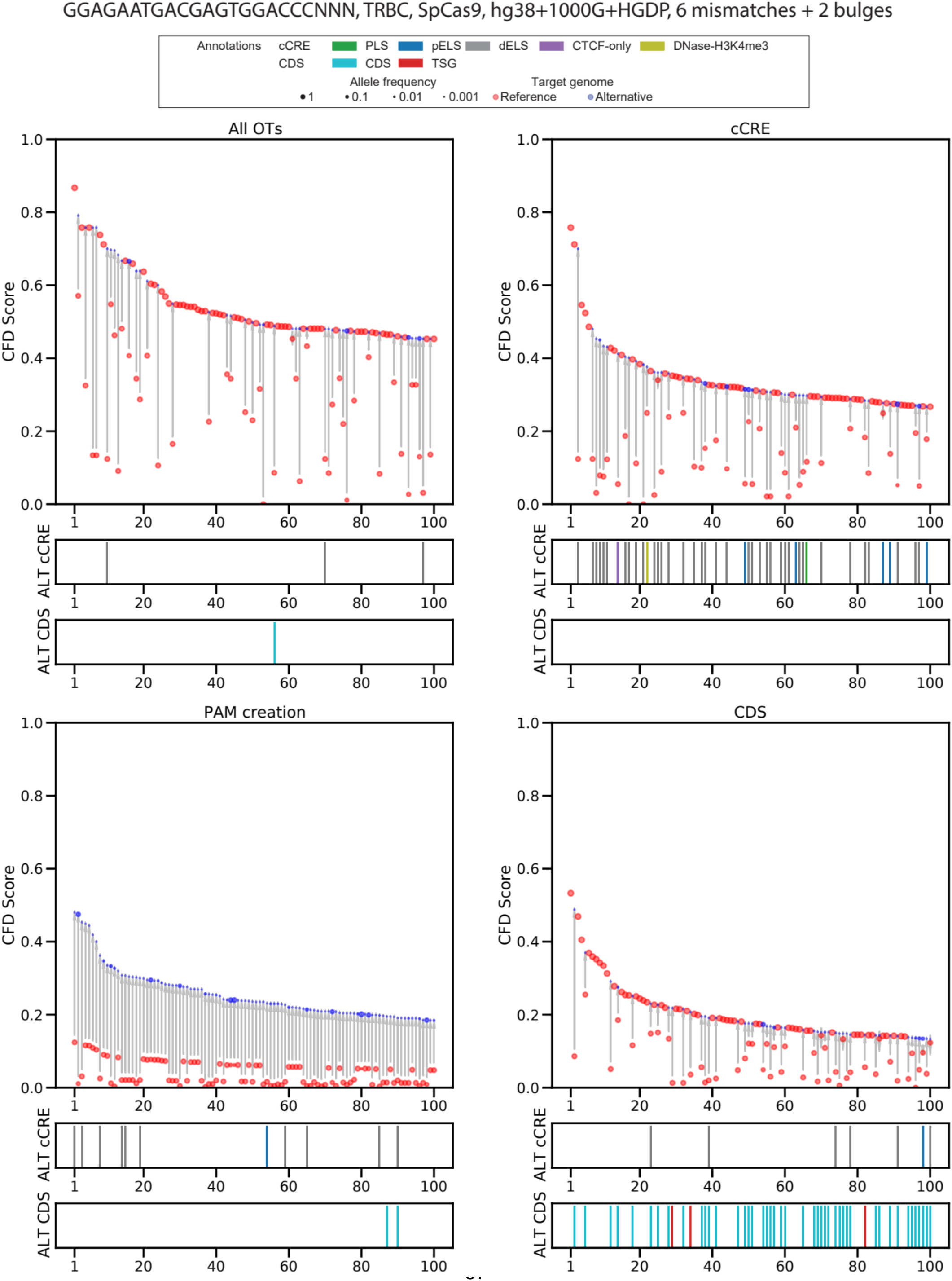
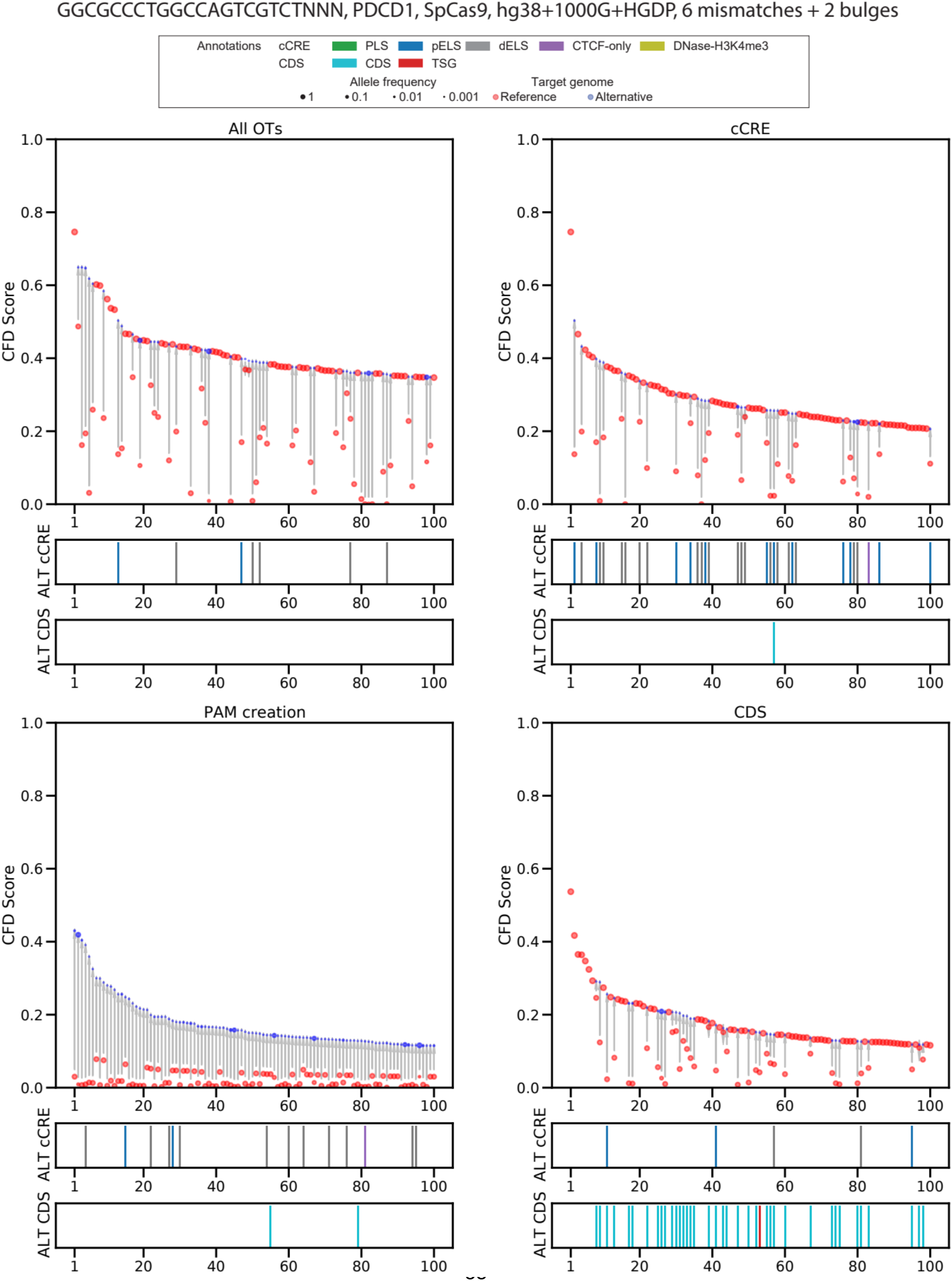
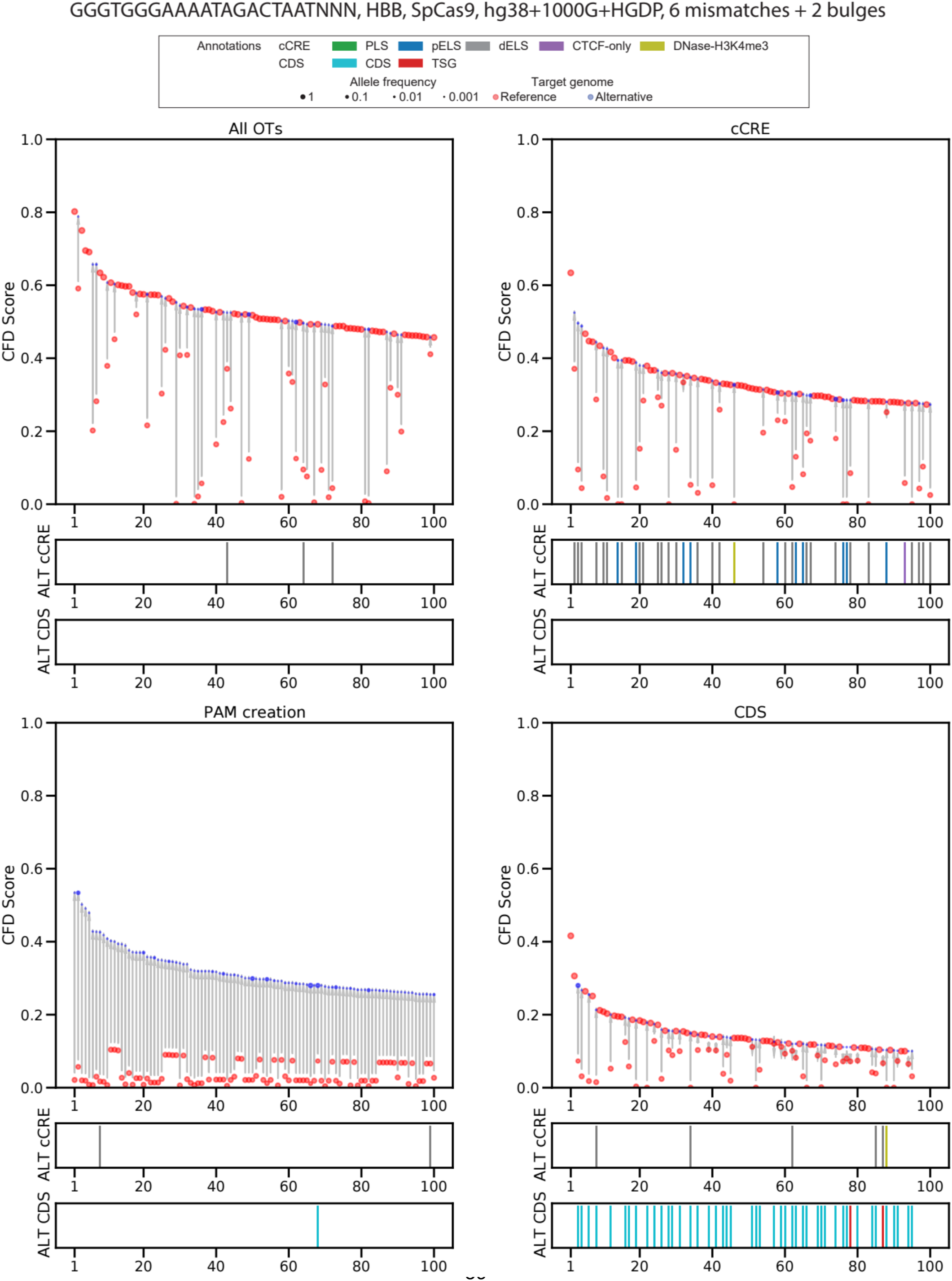
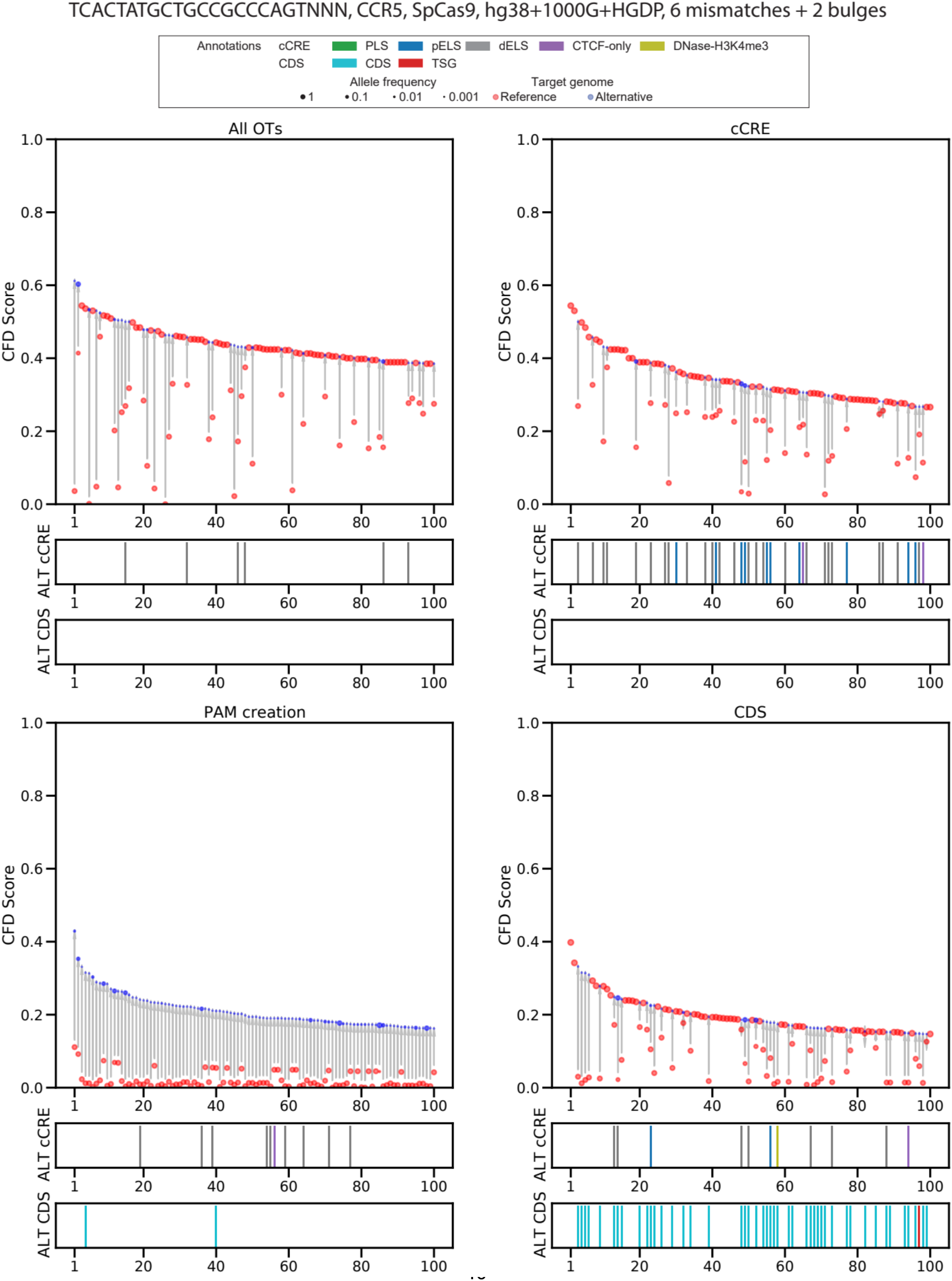
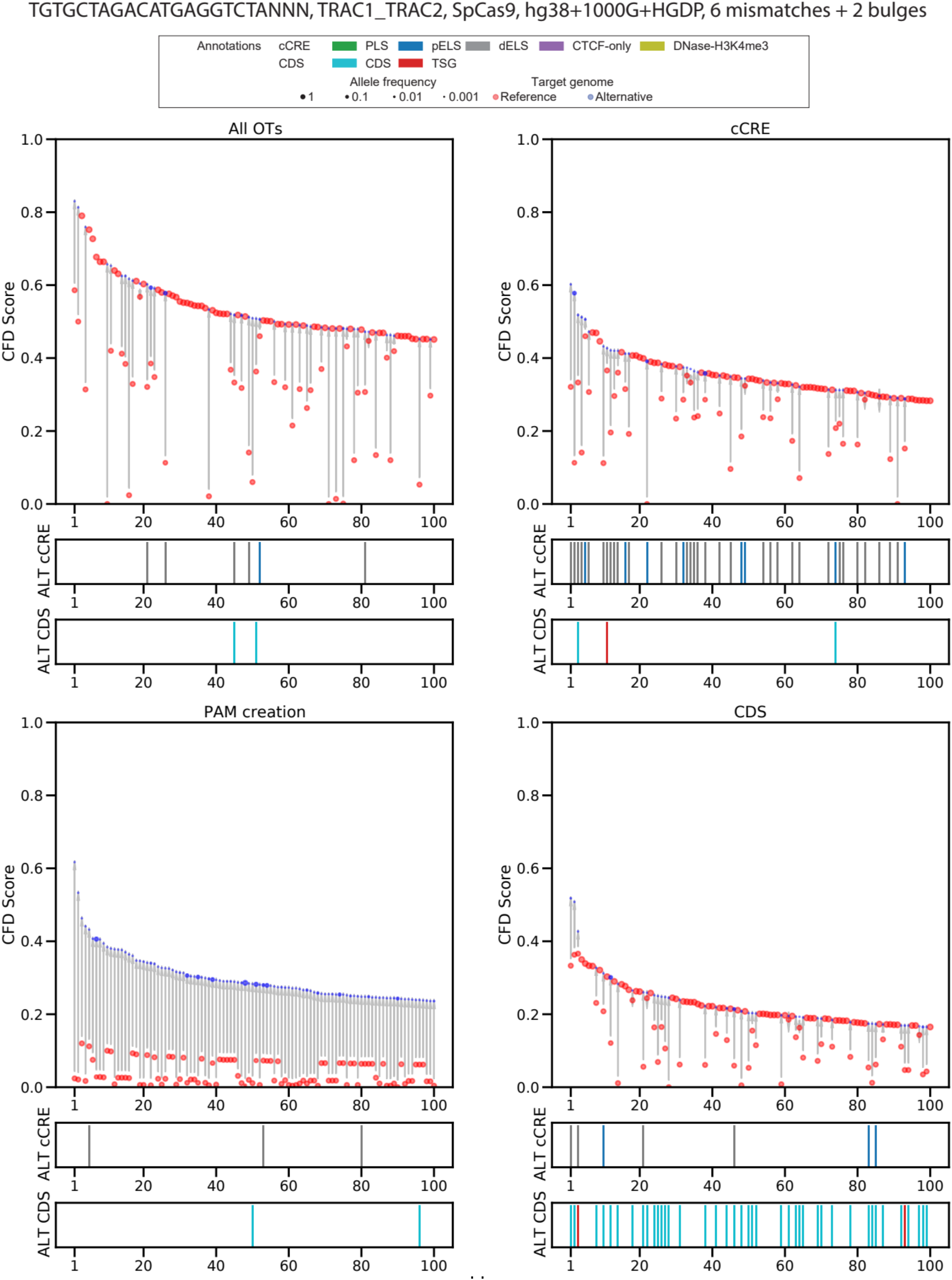
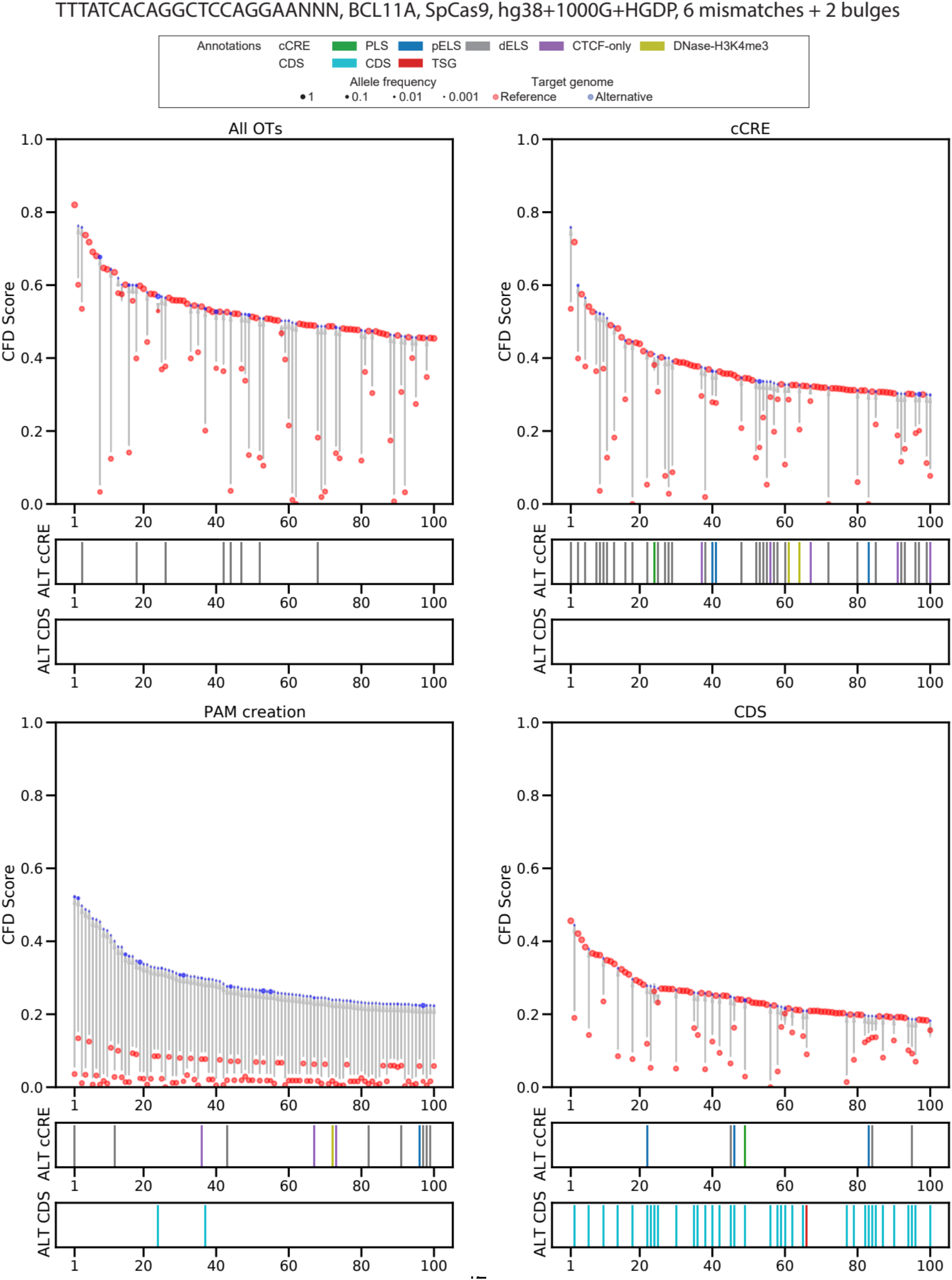
Set of plots representing reference and variant targets for 14 selected sgRNAs (including sg1617) and the variation in terms of CFD score induced by variant introduction. Top 100 predicted off-target sites ranked by CFD score, indicating the CFD score of the reference and alternative allele if applicable, with allele frequency indicated by circle size. Title lists spacer+PAM sequence, target gene, editor, genome (hg38 with 1000G and HGDP variants for all searches shown), mismatch and bulge threshold (6 and 2 for all searches shown). All OTs plot reports the top 100 scoring targets for the sgRNA without any filter. cCRE plot reports the top 100 scoring targets annotated as cCRE elements using ENCODE annotation data. PAM creation reports the top 100 scoring targets with a PAM creation event induced by variant introduction in the reference genome. CDS plot reports the top 100 scoring targets annotated as CDS using GENCODE annotation data.

**Supplementary Figure 5.**
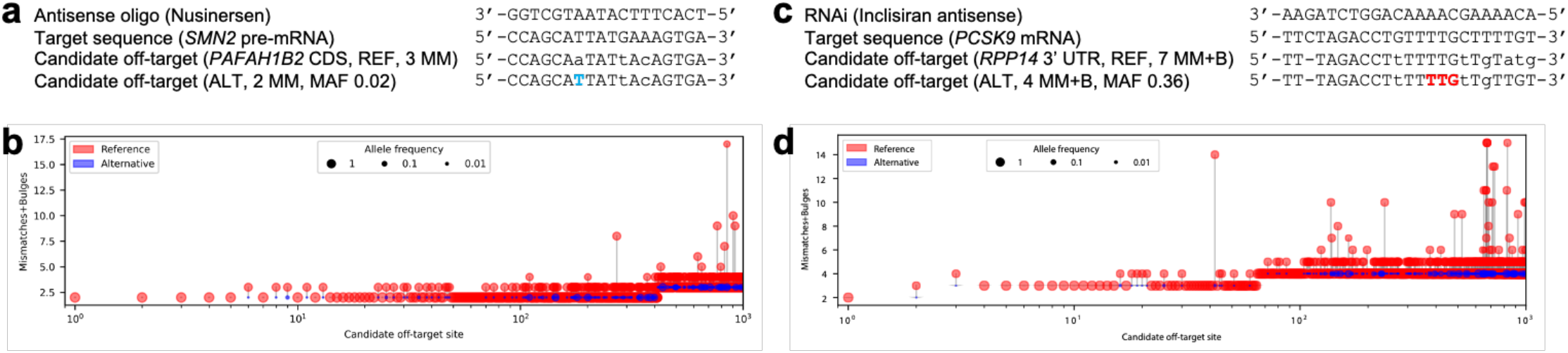
Candidate transcript off-targets introduced by common genetic variants for non-CRISPR sequence-based RNA-targeting therapeutic strategies. a) A common SNP (in blue) introduces a candidate CDS off-target site with 2 mismatches for the FDA-approved antisense oligo Nusinersen. b) Top 1000 candidate transcript off-targets ranked by mismatches and bulges for Nusinersen from a search performed with the 1000G and HGDP genetic variant datasets. c) A common insertion variant (in red) introduces a candidate 3’UTR off-target site with 4 mismatches + bulges for the FDA-approved RNAi therapy Inclisiran. d) Top 1000 candidate transcript off-targets ranked by mismatches and bulges for Inclisiran from a search performed with the 1000G and HGDP genetic variant datasets.

### Supplementary Note 1. CRISPRme web-based search and input requirements

CRISPRme is available as an online web app at http://crisprme.di.univr.it/ (tested for compatibility with Google Chrome and Mozilla Firefox), or offline as a local web app or a standalone command line package (see **Supplementary Notes 5 and 9**). The required inputs to perform an online search are: gRNA spacer(s), Cas protein, PAM sequence, genome build, and thresholds of mismatches and DNA/RNA bulges. Genetic variant datasets (1000G, HGDP and/or personal variants) and annotations can be included as optional inputs. can be included as optional inputs.

A CRISPRme search can be performed in three simple steps thanks to the user-friendly interface (Supplementary Figure 6). Several options are available to personalize a search.

**Supplementary Figure 6.**
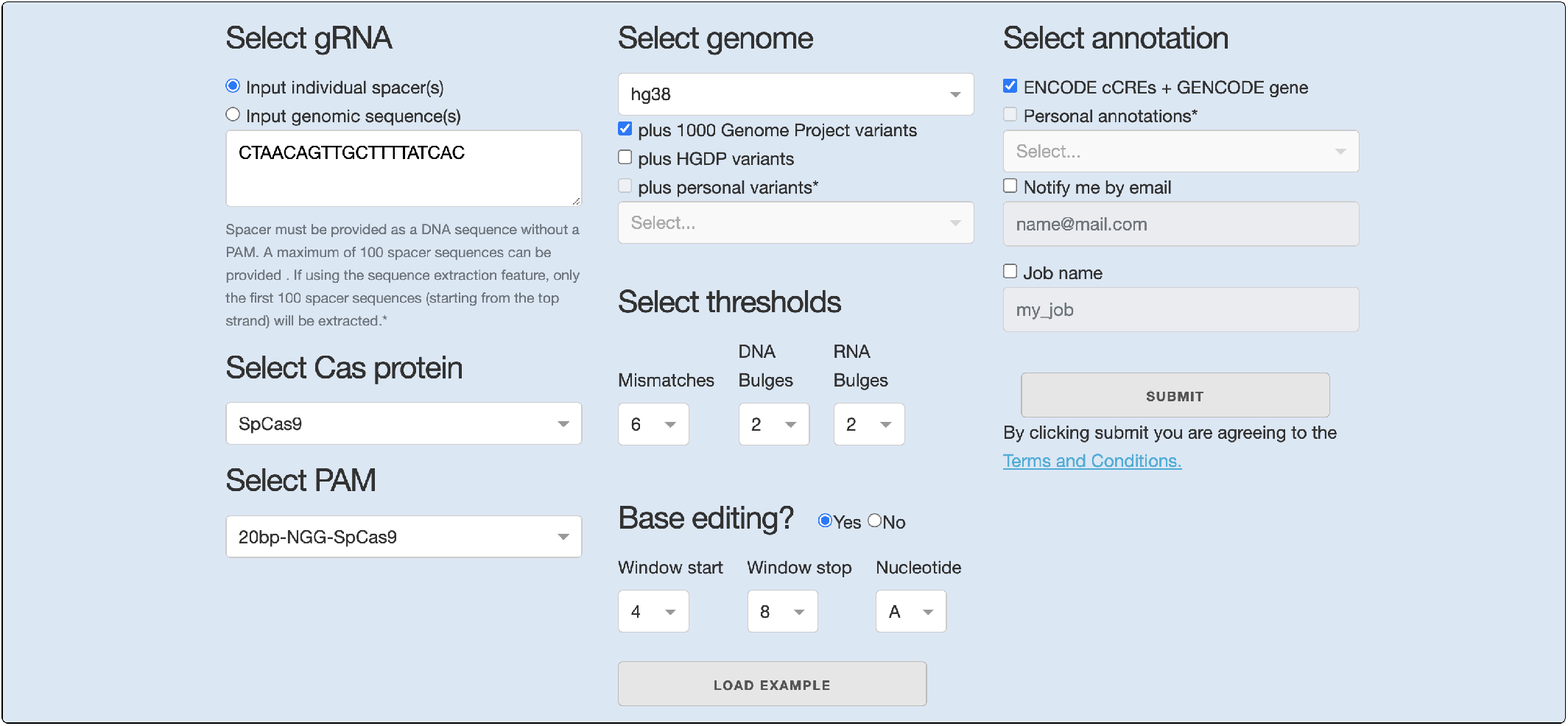
CRISPRme user interface.

#### STEP 1: Spacer, Cas protein and PAM selection

##### Spacer(s)

The guide RNA (gRNA) spacer sequence matches the genomic target protospacer sequence (typically 20 nucleotides) and directs Cas protein binding to the protospacer in the presence of a protospacer adjacent motif (PAM). The spacer sequence is represented as DNA (rather than RNA) in CRISPRme to allow easy comparison to the aligned protospacer sequence. CRISPRme accepts a set of gRNA spacer(s), one per line, each with the same length (max 100 sequences in the online version). The input spacer sequence should not include a PAM.

An example of a gRNA spacer: CTAACAGTTGCTTTTATCAC

##### Genomic sequence(s)

CRISPRme can alternatively take as input a set of genomic coordinates in BED format (chromosome start end) or DNA sequences in FASTA format (max 1000 characters in the online version). The BED coordinates will be treated as 0-based and CRISPRme (online version) will extract the first 100 possible spacer sequences within these coordinates starting with the positive strand. To use this type of input, the user must delimit each entry with a >header.

An example of BED coordinates:

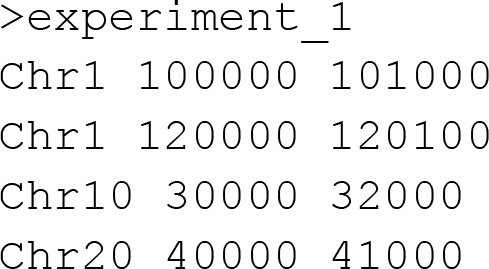

An example of a DNA sequence:

`>BCL11A AAGAGGTGAGACTGGCTTTTGGACACCAGCGCGCTCACGGTCAAGTGTGCAGCGGGAGGAAAGTAGTCATCCCCACAATA`

##### PAM sequence

The PAM is a short (∼2-6 nucleotide) DNA sequence adjacent to the protospacer necessary for the Cas protein to bind to a specific DNA target. CRISPRme (online version) supports a variety of PAMs and users must select one of them in order to perform the search. The software supports both 3’ (e.g. SpCas9) and 5’ (e.g. Cas12a) PAM sequences.

An example of a PAM: `NGG`

#### STEP 2: Genome selection and threshold configuration

##### Genome builds

The genome builds are based on FASTA files from UCSC, so any references available in FASTA format will be supported (such as transcriptomes, genomes from other organisms, and cancer genomes). The <u>hg38</u> genomic build, which includes mitochondrial DNA, is available by default with the option to incorporate variants from 1000G and/or HGDP in the search. The option to add personal variants is enabled only for the local offline and command line versions. For RNA-targeting strategies, the user can currently either input a personal transcriptome to search or use a (variant-enriched) genome, although the latter will miss off-targets found at splice junctions.

##### Search thresholds

CRISPRme allows users to specify the number of mismatches, DNA and RNA bulges tolerated in enumerating potential off-targets. The web-tool allows up to 6 mismatches and up to 2 DNA/RNA bulges (which can be consecutive (`NN--NN`) or interleaved (`NN-N-NN`)). For the local web and command line versions, these thresholds can be set freely and depend only on the available computational resources (see **Supplementary Note 9**).

##### Base editing thresholds (optional)

CRISPRme allows users to specify the window for base editing susceptibility if a base editor is selected as the Cas protein (see Supplementary Figure 6). The “Window start” and “Window stop” dropdowns are limited by the length of the input guide and determine where the “Nucleotide” should be searched for within the putative off-/on-target. The tool produces a final integrated file indicating the base editing susceptibility of candidate off-targets.

**Supplementary Figure 7.**
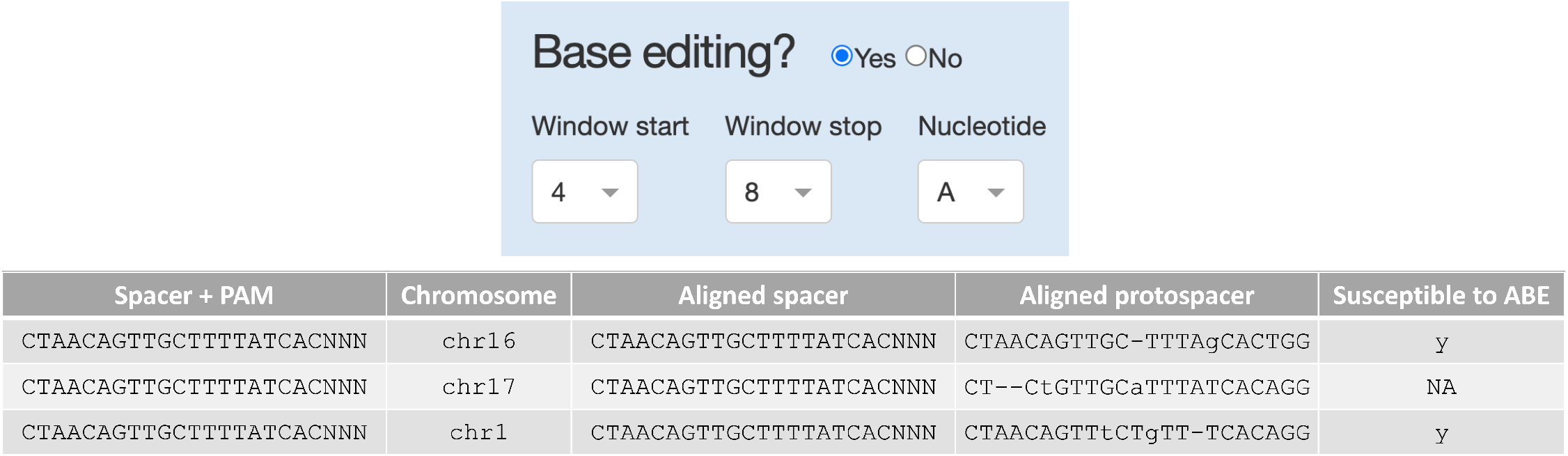
Base editing options to flag potential off-target sites susceptible to base editing. Example of two targets susceptible to A base editing and one target with no susceptibility. If a target contains the selected nucleotide in the chosen window, it is reported as susceptible to <NT>BE. If the nucleotide is not present in the window, it is reported as non-susceptible.

#### STEP 3: Annotation(s), email notification, and job name

##### Functional annotations (optional)

To assess the potential impact of off-target activity, CRISPRme provides a set of functional annotations for coding and non-coding regions. The annotations are based on files obtained from the Encyclopedia of DNA Elements (ENCODE) containing candidate cis regulatory elements^1^ and from GENCODE containing annotations for protein coding genes. In the offline versions of CRISPRme, users can add custom genome annotations, such as cell-type specific chromatin marks or off-target sites nominated by in vitro and/or cellular assays as simple BED files (see **Supplementary Note 5**).

##### Email notification (optional)

If an email address is provided, the user will receive a notification with a link to the results upon job completion.

##### Job name (optional)

If a job name is provided, it will be added as a prefix to the unique job ID to facilitate identification of a particular search e.g. my_job_G05B8KHU0H.

After selecting the desired inputs, clicking the Submit button starts the search. A new page will show the search progress, and upon completion, a “View Results” link will appear at the bottom of the status report page (Supplementary Figure 8).

**Supplementary Figure 8.**
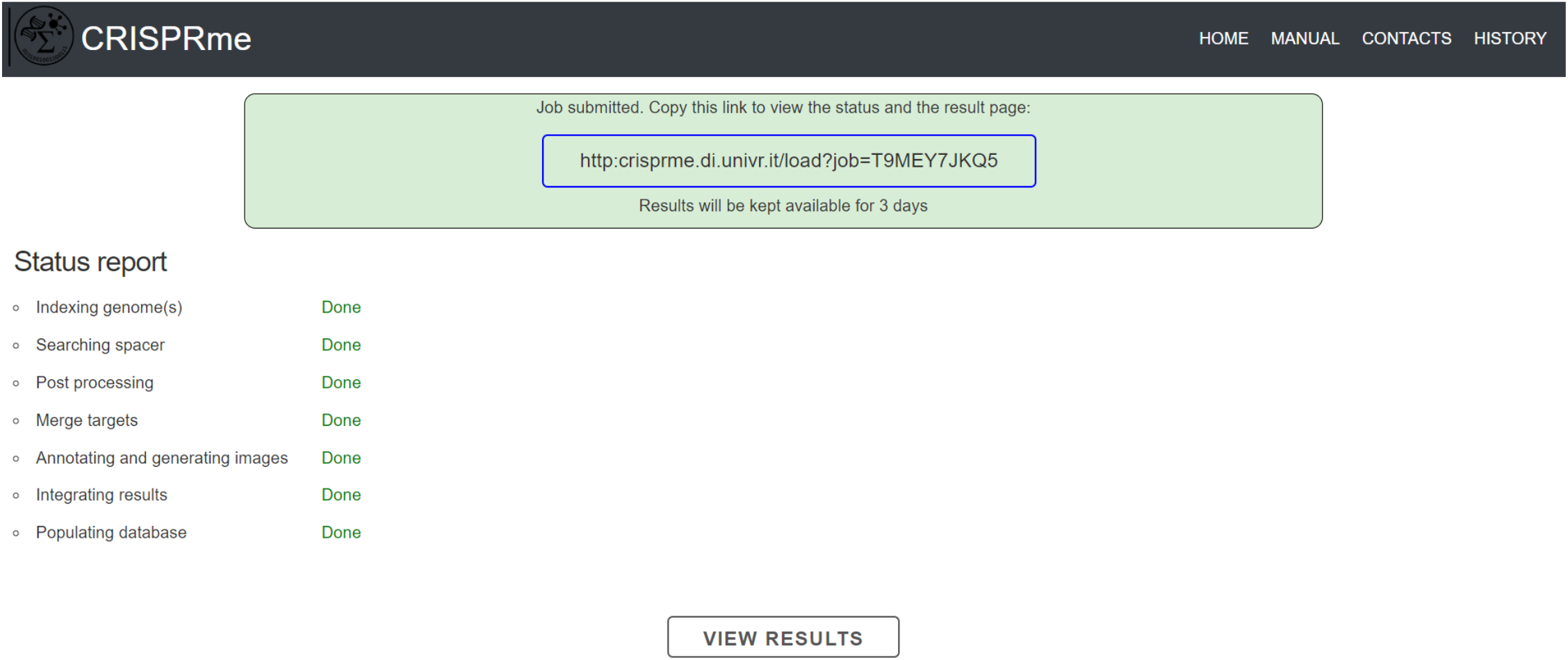
CRISPRme status report page.

The available results and graphical reports are described in detail in the next section.

### Supplementary Note 2. CRISPRme output and graphical reports

CRISPRme summarizes the results in a table highlighting for each gRNA its CFD specificity score and the count of on-targets and off-targets found in the reference and variant genomes grouped by number of mismatches and bulges (**Supplementary Figure**). Of note, the CFD specificity score was initially proposed for searches of up to three or four^2^ mismatches; as the number of mismatches increase, the specificity score decreases non-linearly. Importantly, these scores should be compared with caution between searches with different numbers of mismatches/bulges and/or different genetic variant datasets.

**Supplementary Figure 9.**
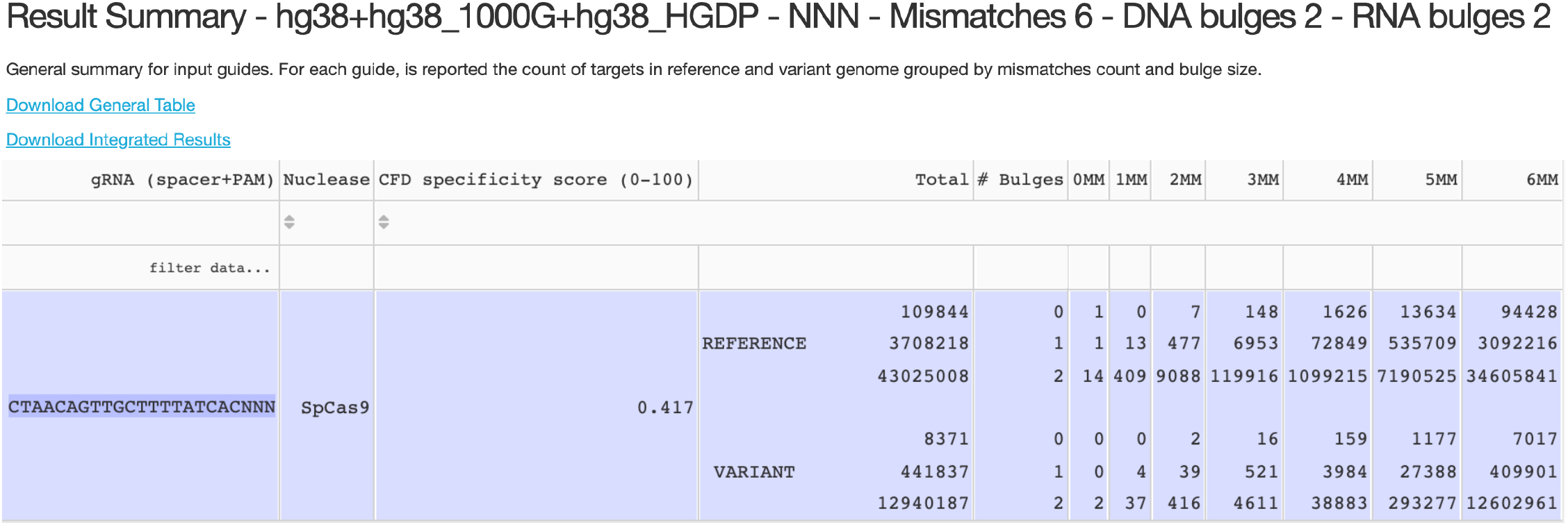
CRISPRme result summary. A table summarizing results based on the search with sg1617, NNN PAM, up to 6 mismatches and 2 DNA or RNA bulges on the human reference genome supplemented with the 1000G dataset with 5 super-populations as well as HGDP with 7 super-populations. The table reports the nuclease, the CFD specificity score and the number of targets in each category of mismatches and bulges. In the top left corner there is a “Download General Table” button allowing download of the table as a text file, as well as a “Download Integrated Results” button allowing download of the full results.

In addition, for each guide, six different interactive reports are generated and are available for download: *Custom Ranking, Summary by Mismatches/Bulges, Summary by Sample, Query Genomic Region, Graphical Reports* and *Personal Risk Cards* (described in **Supplementary Note 3**).

#### Custom Ranking

In this report, users can filter and rank potential off-targets based on number of mismatches and/or bulges, CFD/CRISTA score, Risk Score (increase in score due to genetic variants), or a combination of these options (Supplementary Figure 10).

**Supplementary Figure 10.**
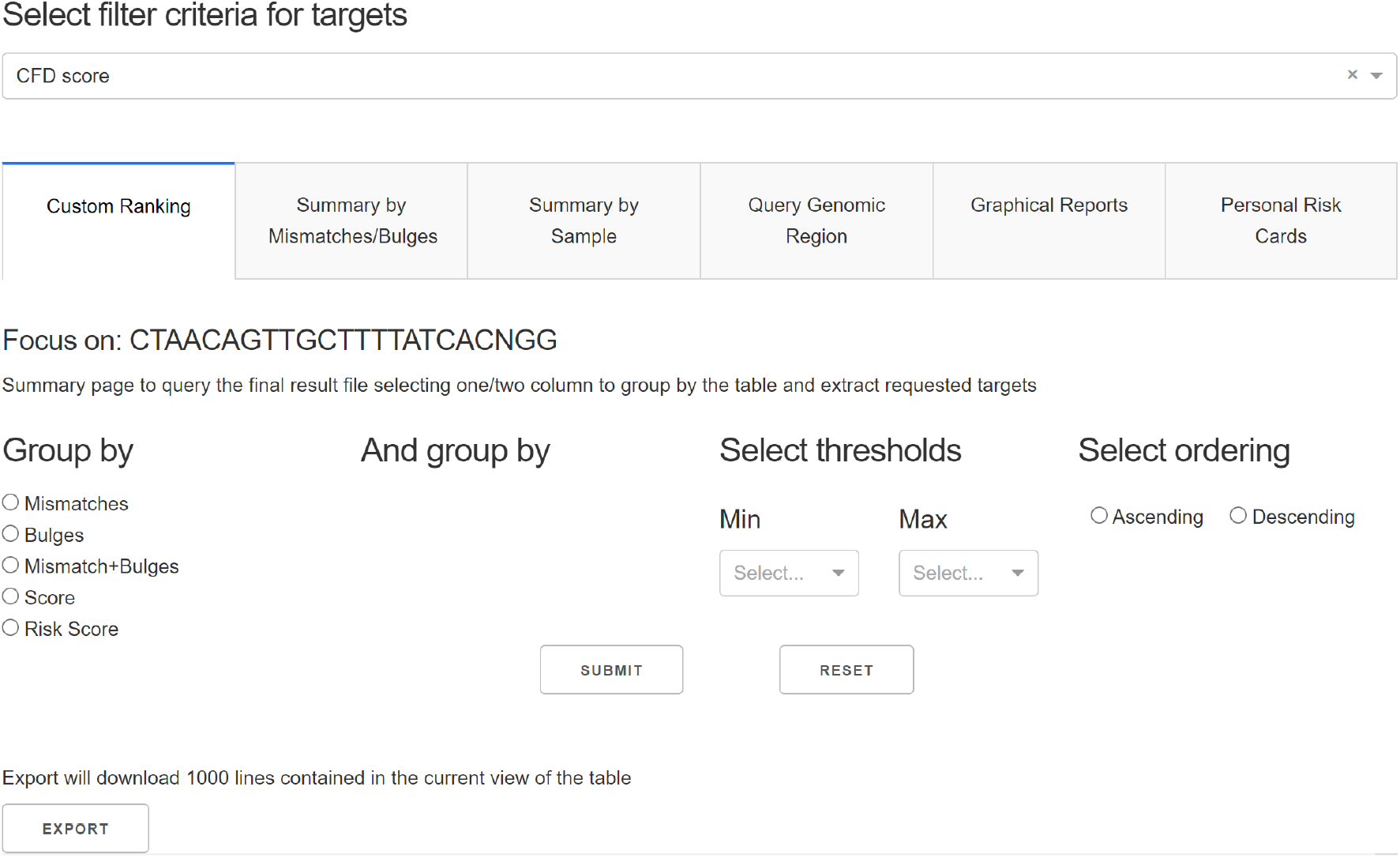
CRISPRme ranking and filtering of off-targets. Users can define filters, orders and group-by operations to easily retrieve results based on a custom logic suitable for their application. All the columns in the table are explained in Supplementary Table 1. Shown here are the first seven columns, containing sequence and positional information.

#### Summary by Mismatches/Bulges

This report shows a matrix separating off-targets into subgroups based on the type, mismatch count and bulge size. “X” targets contain only mismatches, “DNA” targets contain DNA bulges (and may contain mismatches), and “RNA” targets contain RNA bulges (and may contain mismatches) (Supplementary Figure 11).

**Supplementary Figure 11.**
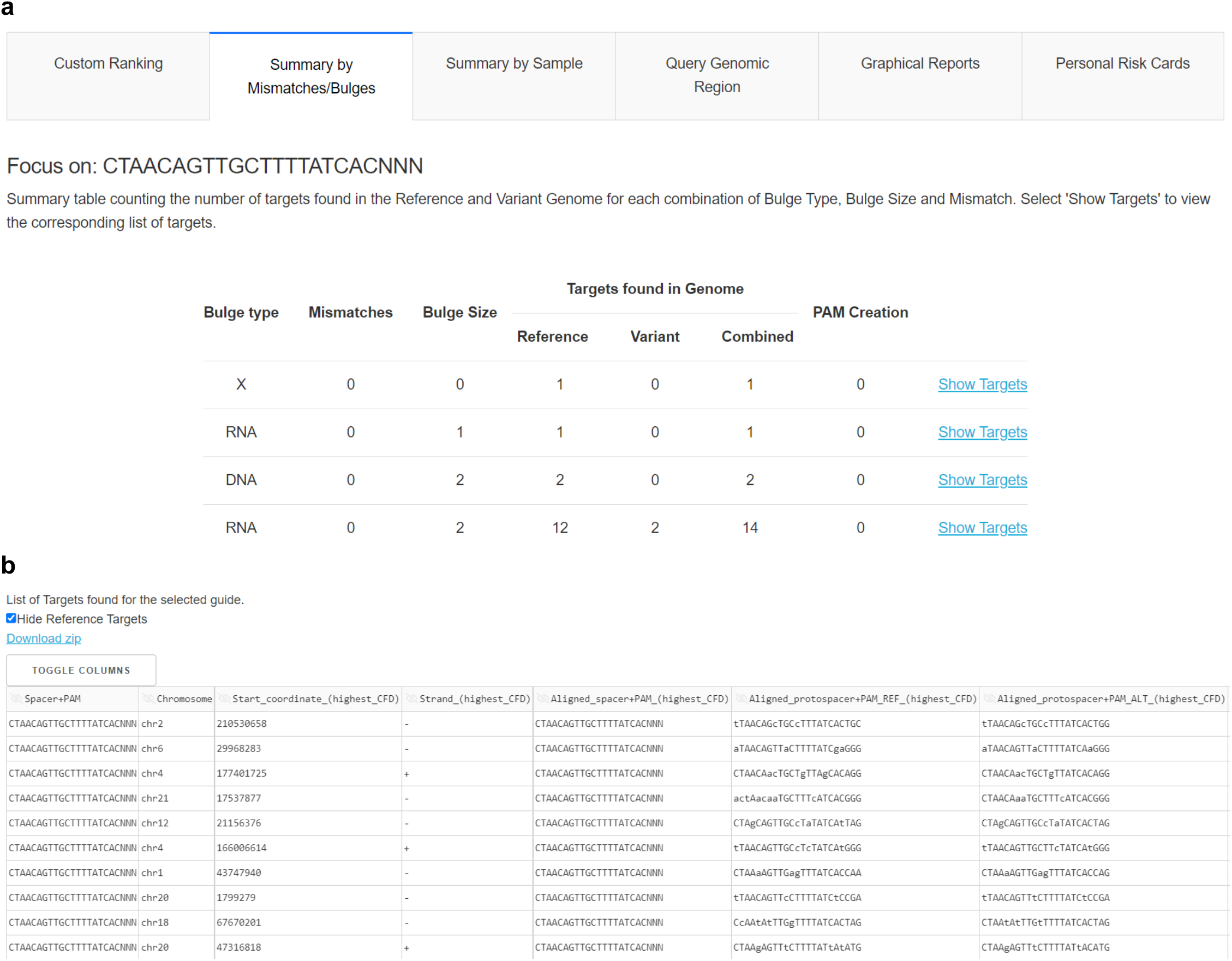
CRISPRme summary results by mismatches/bulges. a) Mismatches/Bulges summary table showing the first 4 of 33 rows for a search with up to 6 mismatches and 2 DNA or RNA bulges. The combined column indicates the sum of reference and variant off-targets. b) View of “Show Targets” with 3 mismatches and no bulges. The user can select which column to see using the “Toggle Columns” button on top of the table. All the columns in the table are explained in Supplementary Table 1. Shown here are the first seven columns, containing sequence and positional information.

#### Summary by Sample

This page shows all the samples present in the VCFs and allows users to extract and visualize targets related to each sample (Supplementary Figure 12).

**Supplementary Figure 12.**
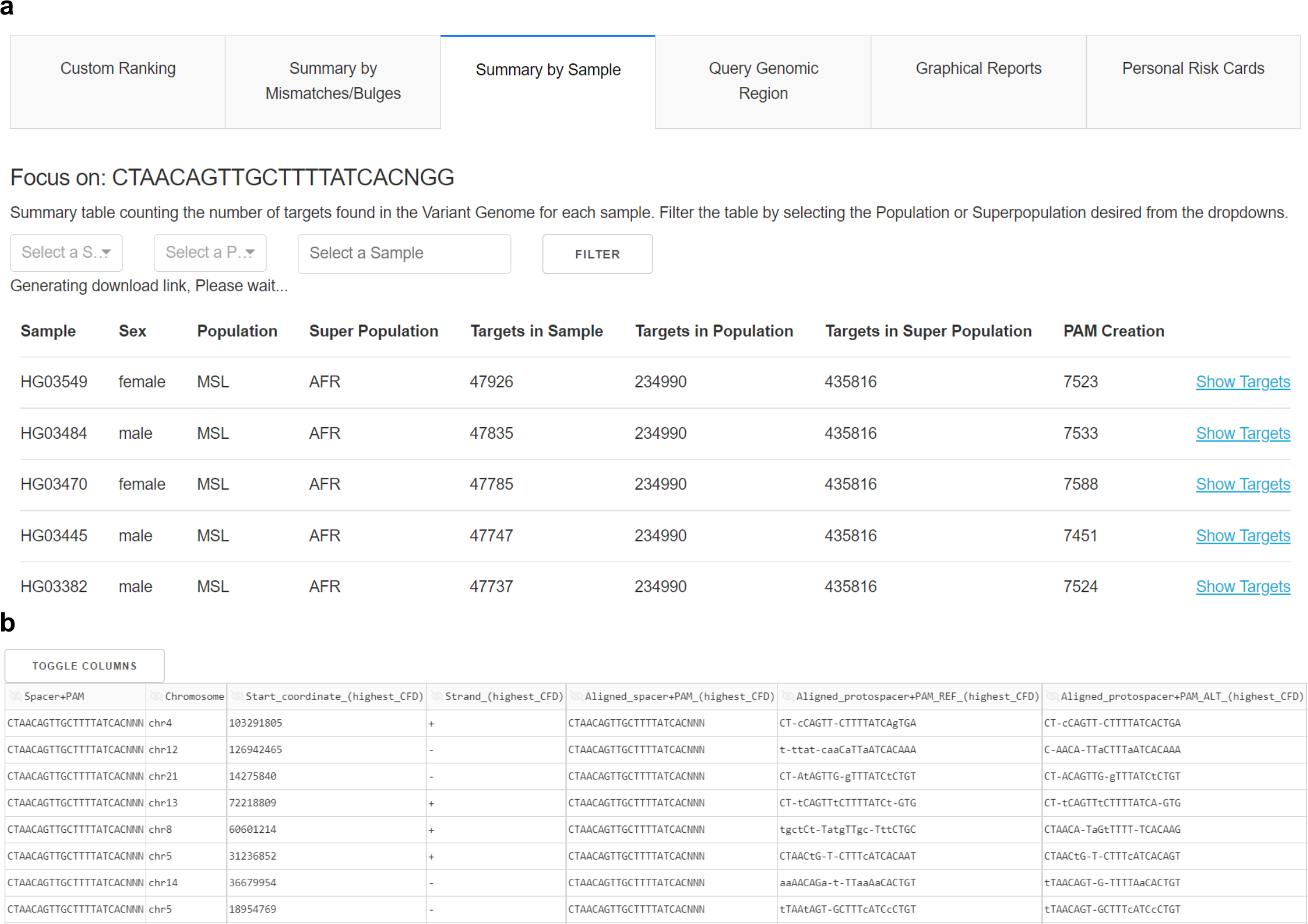
CRISPRme results by sample. **a)** Samples, alongside their sex, population and super-population information, are shown in a tabulated list with the count of variant off-targets for the sample, its population and super-population, as well as the number of PAM creation events for the sample. **b)** View of “Show Targets” for HGDP01211. The user can select which columns to see using the “Toggle Columns” button on top of the table. All the columns in the table are explained in Supplementary Table 1. Shown here are the first seven columns, containing sequence and positional information.

#### Query Genomic Region

This page allows the user to retrieve off-targets overlapping a specific genomic region, for example to quickly assess potential off-targets in a given regulatory element or coding region (Supplementary Figure 13).

**Supplementary Figure 13.**
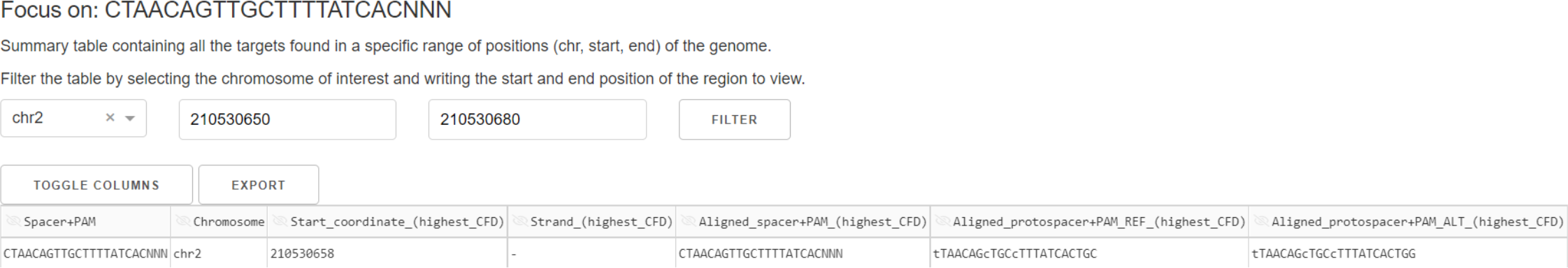
CRISPRme results by genomic region. A table showing the candidate off-target(s) within the region. All the columns in the table are explained in Supplementary Table 1. Shown here are the first seven columns, containing sequence and positional information

#### Graphical Reports

This page creates several graphical reports for each selected gRNA.

- A stem plot (Supplementary Figure 14a) shows how genetic variants affect predicted off-target potential. The arrow connecting the red (reference allele off-target) and blue (alternative allele off-target) dots shows the increase in predicted cleavage potential due to the variant.
- Bar plots depict how candidate off-targets are distributed across super-populations based on the number of mismatches and bulges (Supplementary Figure 14b).
- A radar chart based on annotations from GENCODE and ENCODE. A larger area in the chart represents a gRNA with more potential off-targets falling in annotated regions, possibly representing an undesirable outcome. A summary table provides the count and percentage of off-targets with a given annotation (Supplementary Figure 14b).
- A motif logo summarizing the frequency of mismatches and bulges (b) among the predicted off-targets for each base of the protospacer + PAM (Supplementary Figure 14b).

**Supplementary Figure 14.**
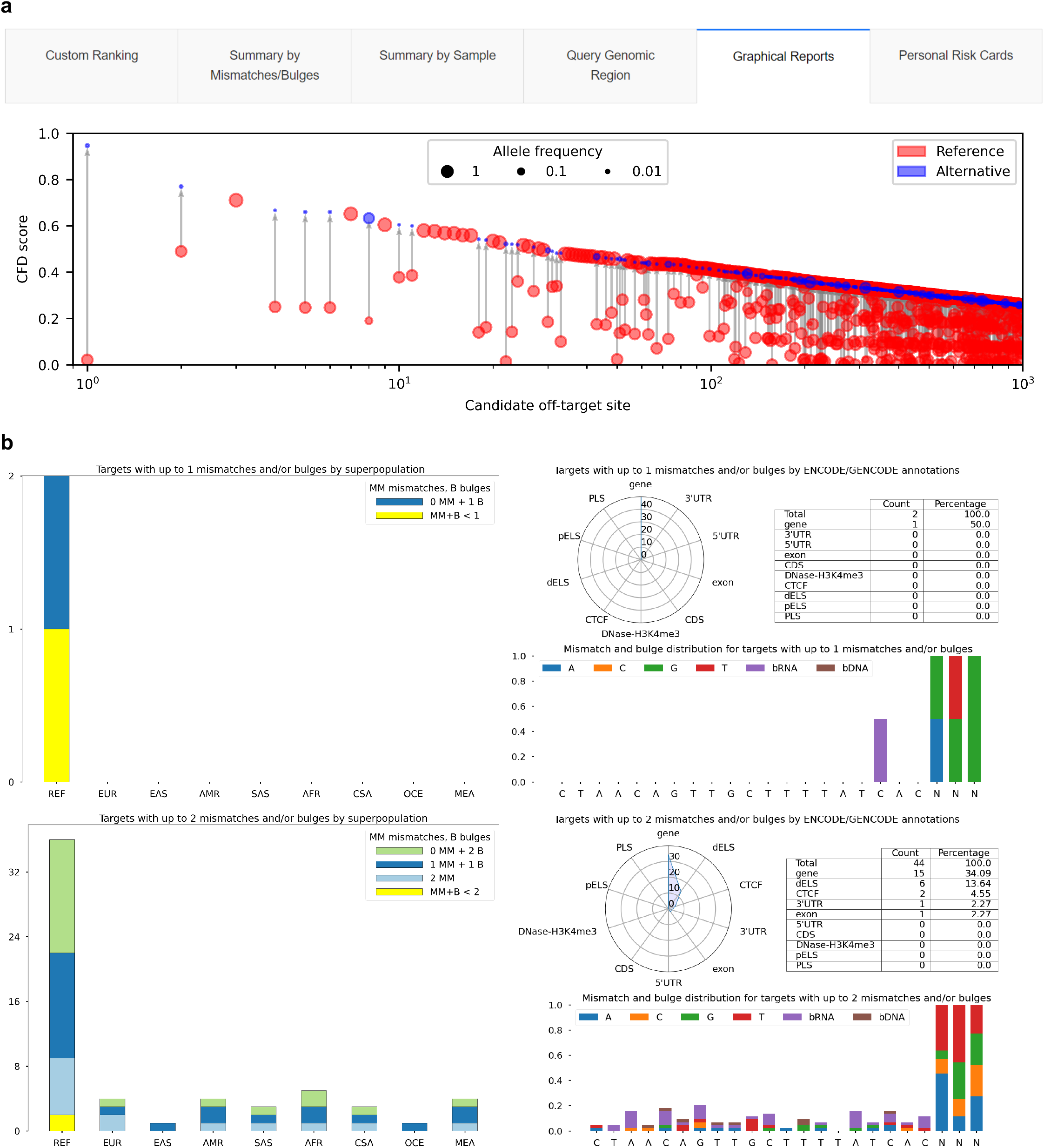
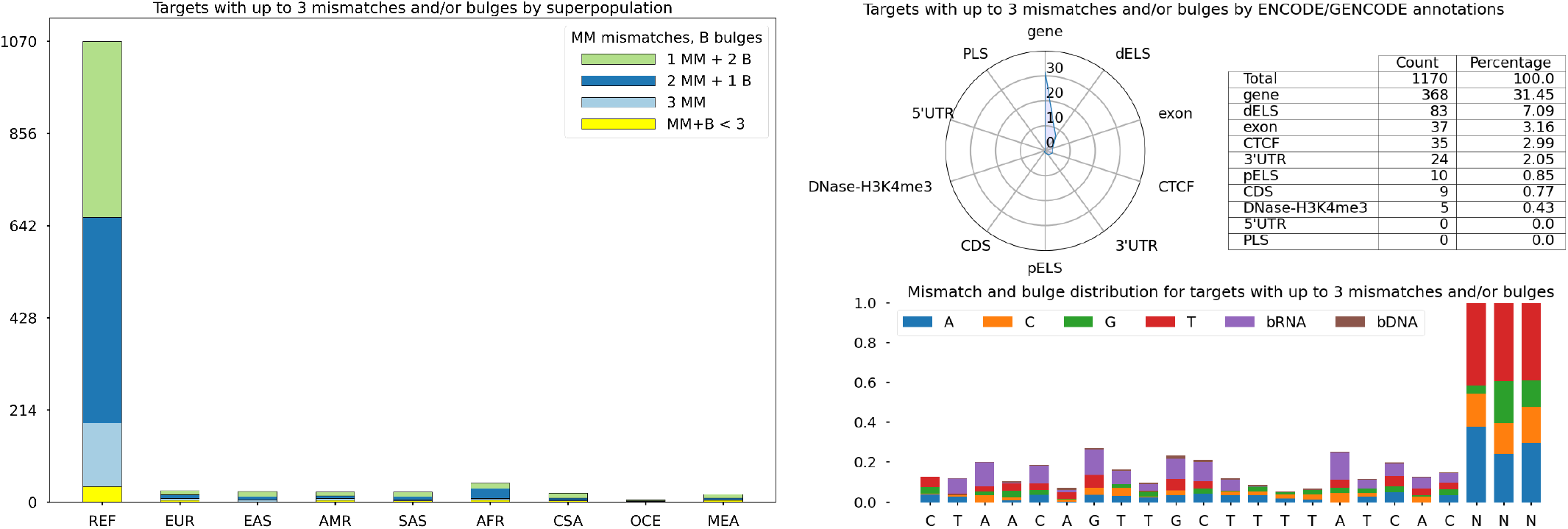
CRISPRme graphical reports. **a)** CRISPRme reference/alternative CFD comparison obtained using sg1617, NNN PAM and 6 mismatches plus 2 DNA/RNA bulges, tested on the hg38 reference genome plus 1000G and HGDP variants. A stem plot shows the distribution of CFD scores for candidate off-targets ranked in descending order by CFD score. For candidate off-targets for which a genetic variant increases the CFD score, the CFD scores for both the alternative (blue) and reference (red) allele at the same locus is shown. The area of the circle is proportional to allele frequency. **b)** On the left, stacked bar plots summarizing the number of candidate off-targets for each category of mismatch + bulge in super-populations present in the input variant data. On the right, radar chart showing the percentage of off-targets falling into a specific genomic annotation with respect to the overall count, table detailing the exact number of off-targets falling into each category (an off-target can fall under more than one category) and motif plot showing the distribution of mismatches and bulges with respect to the spacer+PAM.

### Supplementary Note 3. CRISPRme personal risk cards

CRISPRme provides a dedicated page to generate reports called *Personal Risk Cards* that summarize potential off-target editing by a particular gRNA in a given individual due to genetic variants. This feature is particularly useful for retrieving and investigating private off-targets.

The report contains two dynamically generated plots depicting all the candidate variant off-targets for the sample including those non-unique to the individual and those that are unique to the individual (Supplementary Figure 15a). These plots highlight how the introduction of genetic variants can change the predicted off-target cleavage potential, thereby demonstrating the importance of variant-aware off-target assessment as in CRISPRme. The report also contains two tables (Supplementary Figure 15b), consisting of a summary (top) and information on each extracted candidate off-target (bottom) with the following columns:

Top:

- `Personal`, count of all the candidate variant off-targets for the selected sample (including both variants unique and non-unique to the individual).
- `PAM creation`, count of all the instances where a genetic variant in the sample introduces a new PAM and the PAM used in the search in not found in the reference genome at the same locus.
- `Private`, count of all the candidate variant off-targets uniquely found in the selected sample.

Bottom: columns explained in Supplementary Table 1.

The personal card table file (as shown in Supplementary Figure 15b) is downloadable as a file (**Supplementary File 1**).

**Supplementary Figure 15.**
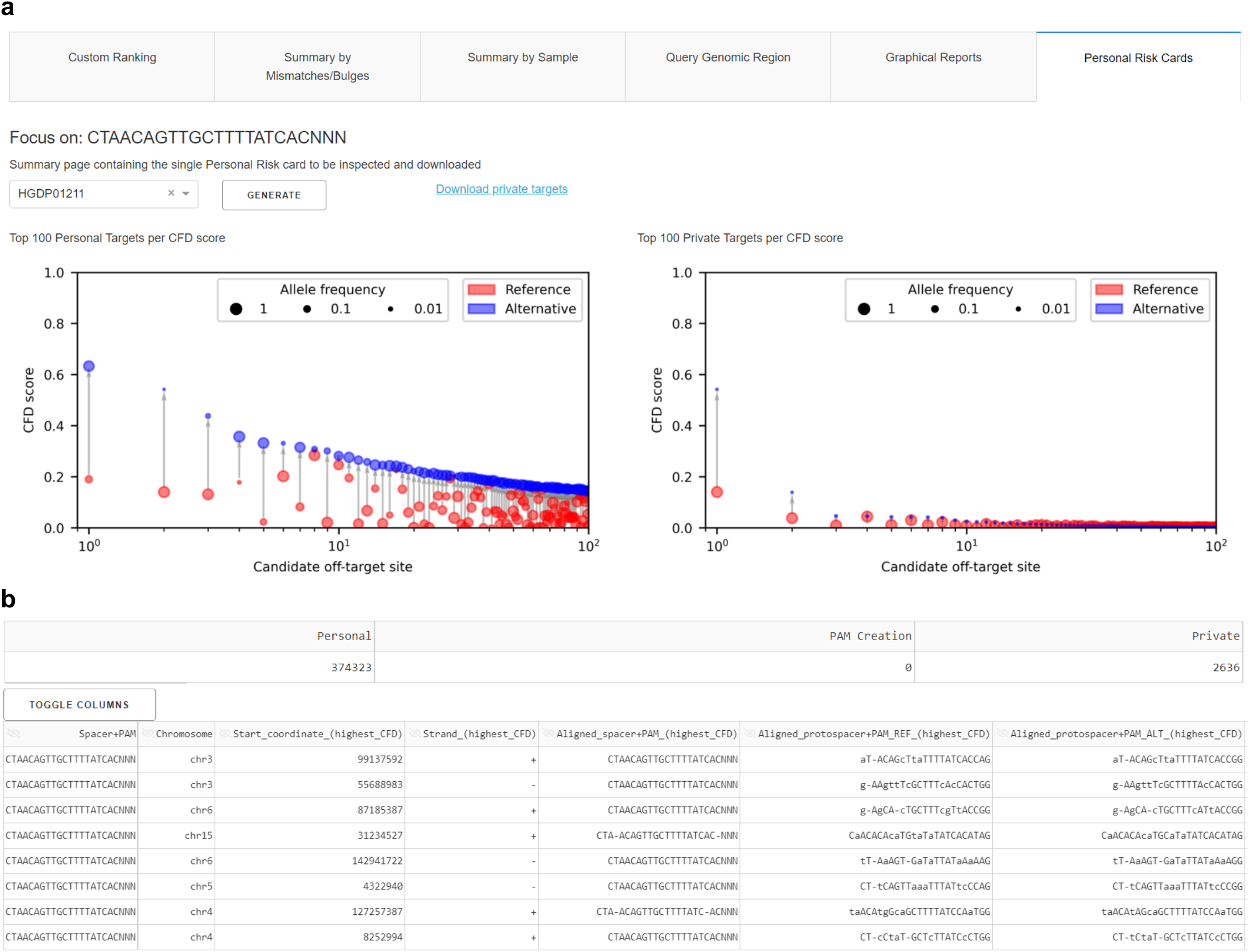
CRISPRme personal risk card. Example shown is for HGDP01211. **a)** *Left:* Plot of potential variant off-targets for the selected sample. *Right:* Plot of potential off-targets unique to the selected sample. **b)** The top table reports the number of personal variant off-targets, instances of PAM creation and private off-targets. The bottom table lists all the private targets for the selected sample. All the columns in the table are explained in Supplementary Table 1. Shown here are the first seven columns, containing sequence and positional information.

### Supplementary Note 4. Details of the CRISPRme implementation

#### CRISPRme software architectures

The CRISPRme web version and front end was developed in Dash, a Python framework to create responsive and interactive web applications (https://plotly.com/dash/). The back end (graphical report generation and data analysis) is based on Python and bash scripts. The search engine is developed in C++ to exploit its speed and stability and to fully leverage parallel computation and compiler optimizations.

#### CRISPRme genome enrichment with variants

CRISPRme performs the search of potential off-targets based on reference genomic sequences stored in FASTA files. A reference genome can be “enriched” with genetic variants (SNPs and INDELs) encoded in VCF files obtained for example from 1000G, HGDP and/or personal data. Enriched genomes are created by encoding SNPs using IUPAC notation, i.e. nucleotides corresponding to genetic variants can be represented via ambiguous DNA characters. For example, if at a given position the reference allele is G and the alternative allele is A, the tool encodes the two alternatives by using the ambiguous nucleotide R, which corresponds to the IUPAC code for G or A. Based on this procedure, the enriched genome contains all the SNP variants belonging to different samples, including multiallelic sites with three or more observed alleles. CRISPRme treats INDELs differently due to the nature of the variant itself. For each INDEL, it creates a fake chromosome containing the variant DNA sequence and 50 surrounding nucleotides (25 on each side). Finally, the association between samples and variants is based on a hash table to allow efficient querying of which samples contain a given SNP/INDEL. This procedure frees the user from the need to manually produce numerous genomes with variants, greatly simplifying this non-trivial operation and automatizing the association of targets with specific haplotypes to individual(s). This haplotype-aware search removes the possibility of reporting possible off-targets that are not present in any real genome. This operation takes into account the possibility of having more than one variant in any given off-target, generating many possible combinations that need to be analyzed. We created an ad-hoc algorithm that can process the set of candidate off-targets in polynomial time by grouping variants by set of individuals sharing them to avoid recomputing already tested combinations. Furthermore, the algorithm allows for either phased or unphased VCFs to be used as input. If phased VCFs are used, CRISPRme generates haplotypes for the positive and negative strands considering the position of the analyzed variant even when more than one variant is found in the same target. This operation saves a considerable amount of time during the computation and removes any artificial off-targets from the final report.

#### CRISPRme CFD scoring function

To score targets using Cutting Frequency Determination (CFD), CRISPRme uses a matrix generated by the authors who introduced the scoring system based on empirical data. This matrix is composed of all the possible pairs of mismatches between a RNA and DNA sequence with a length of 20 nucleotides. Each entry in the matrix file reports a RNA and DNA nucleotide pairing. For example, the entry “rA:dG,20’, F0.227” means that the RNA nucleotide A paired with the DNA nucleotide G in position 20 will have a score of 0.227. This value is multiplied with values for any other mismatches present to obtain the CFD score for the off-target sequence. If a sequence has only 1 mismatch, its final score will be the score of the mismatch, so in the previous example, the CFD score of an off-target with only one mismatch (20A>G) will be 0.227.

The matrix also contains scores for bulges, which are indicated as “-”. An example entry representing a bulge is “sS’r-:dA,2’, F0.692,” which indicates that a RNA bulge pairing with the DNA nucleotide A at position 2 has a score of 0.692. Bulges are not allowed in the first position of the RNA or DNA sequence. Computationally, targets with DNA bulges are reported with a longer sequence with respect to the original spacer. To avoid inconsistencies when calculating CFD score for off-targets with DNA bulges, we calculate the score based on only the last 20 nucleotides of the protospacer as intended by the original CFD scoring method.

The process also scores the PAM nucleotides. Since CFD score was derived from SpCas9 data, the matrix only contains scores for NGG and all the possible combinations of the last two positions of the PAM.

An example of an off-target with a DNA bulge:

Spacer: CTAACAG**T**TGCTT**-**TTATCACNNN

Protospacer: CTAACAG**c**TGCTT**C**TTATCACC**TC**

This off-target contains one mismatch (in lowercase) and one DNA bulge (aligned with gap in the spacer sequence). When we calculate its CFD score, we do not consider the first nucleotide of the protospacer because the protospacer is one nucleotide longer than the spacer. Then each mismatch and bulge is scored according to the matrix and the value of each pair is multiplied together to obtain the final CFD score.

In this example, the first mismatch is rT:dC in position 7 (we skip the first nucleotide since there is 1 DNA bulge), and the score for that pair is 0.588. Then, we encounter the bulge r-:dC in position 13, and the score of this pair is 0. We keep moving along the sequence until we reach the PAM. In this case, the nucleotide couple TC has a score of 0. Finally, we multiply each value saved during the process, so the final score is calculated as 0.588 * 0 * 0, yielding an off-target CFD score of 0 for this example.

Scoring off-targets with RNA bulges is simpler since the spacer sequences are not elongated due to the bulges. An example:

Spacer: CTAACAGTTGCTTTTAT**C**ACNNN

Protospacer: CTAACAGTTGCTTTTAT**-**ACGTG

This off-target only contains a RNA bulge (protospacer gap). We scan the sequence and encounter the bulge in position 18, so we use matrix entry “rC:d-,18” with score 0. The final CFD score for the off-target will be 0.

#### CRISPRme indexing, search, and analysis

To perform efficient search operations, CRISPRme creates an index of reference/enriched genomes. This index encodes all the possible candidate off-targets in a tree-based data structure and can be used to efficiently find reference or variant enriched sites. However, genomic regions containing Ns (e.g. poorly assembled or repetitive regions) will not be considered during the search operation. In addition, CRISPRme introduces individual-specific analysis, extracting from the IUPAC encoding of the enriched genome haplotype-specific off-targets and the corresponding samples. For each site, CRISPRme also reports off-target potential scores, and if there are multiple possible alignments at a given site (e.g. when there are RNA/DNA bulges), the alignment with the highest score is reported. CRISPRme currently adopts two well-known scoring functions, CFD and CRISTA scores, because they can be efficiently computed for thousands of sites, handle bulges and perform well in predicting off-targets as validated by deep sequencing^3^. CRISPRme in principle could be extended to support other predictive off-target scores.

In addition, a global score called CFD specificity score is provided for each gRNA and defined as follows:

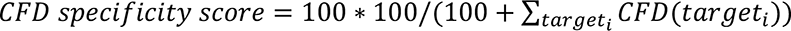

where *target*_i_ is one of the enumerated off-targets from the search. Its range is (0, 100], where a gRNA with no predicted off-targets given the search parameters has a CFD specificity score of 100. Of note, the CFD score is only applicable for SpCas9. CRISPRme does not calculate CFD scores when searching for editors other than SpCas9. In this case, a −1 value is reported for each off-target.

The CRISTA specificity score is calculated similarly as:

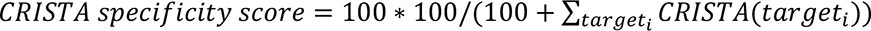

Importantly, thanks to the constructed hash table containing the mapping between samples and variants, after the initial search based on the IUPAC code, CRISPRme filters the results by reporting only targets matching haplotypes that exist in the populations and the corresponding individuals.

Given that multiple alignments may correspond to a given genomic region, CRISPRme outputs two lists of candidate off-target sites. The “integrated_results” file includes a single off-target site per genomic region, merging all possible off-targets within 3 bp (by default, adjustable in the command line version), and integrates annotation information if provided in the input. The off-targets included in the list are selected and sorted based on the highest CFD score by default, but users can select other criteria such as fewest mismatches and bulges by using the dynamic filters available on the website main results page (Supplementary Figure 10). When the CFD score is identical, the reference alignment is favored over alternative alignments. Meanwhile, the “all_results_with_alternative_alignments” file contains all the off-targets not included in the first file. This file preserves alternative alignments for off-targets as well as those containing other variants with lower CFD scores. **Supplementary File 1** provides examples of these files (top 1000 lines due to space constraints).

Candidate off-target files created by CRISPRme include the fields described in Supplementary Table 1.

**Supplementary Table 1.**
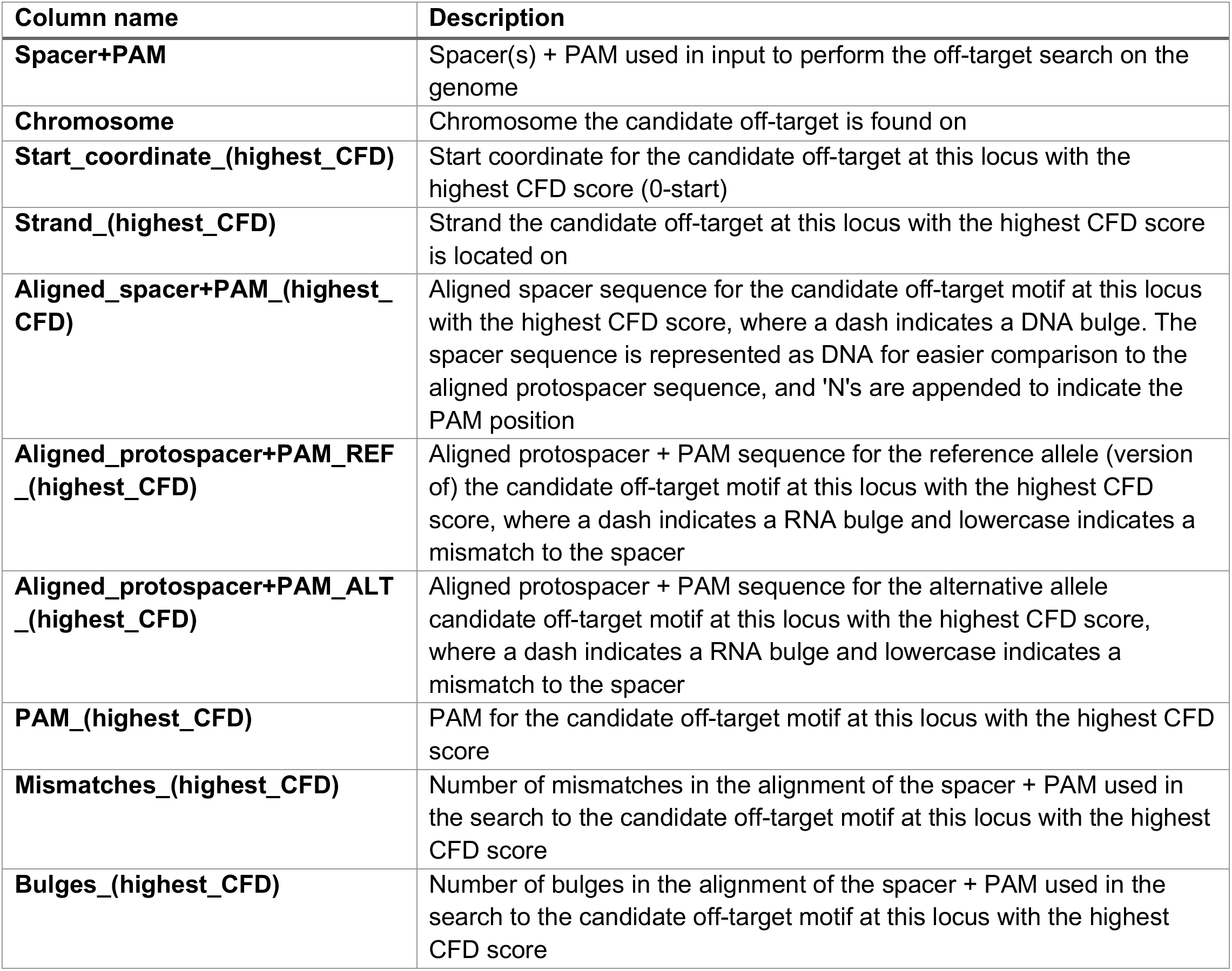
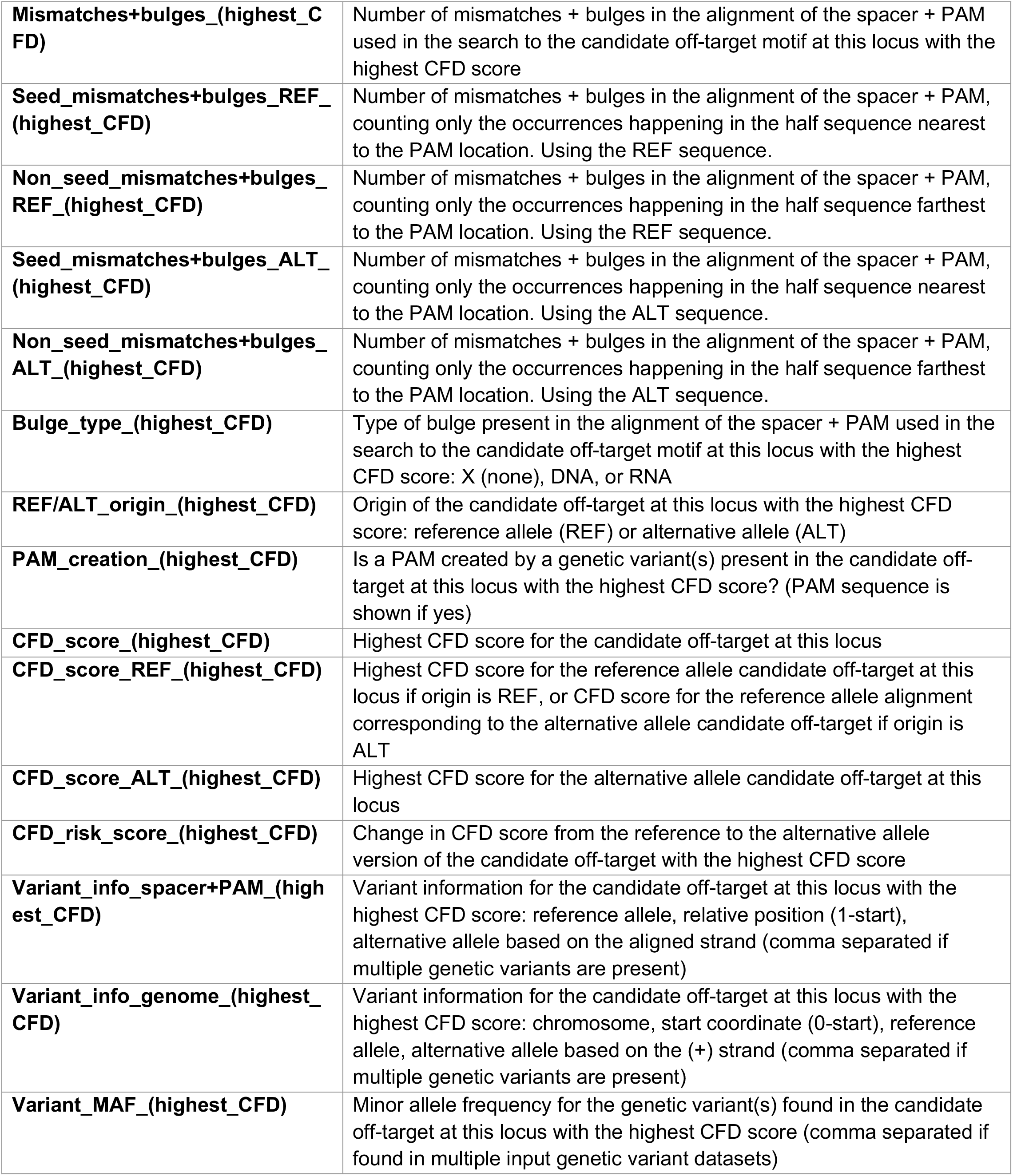
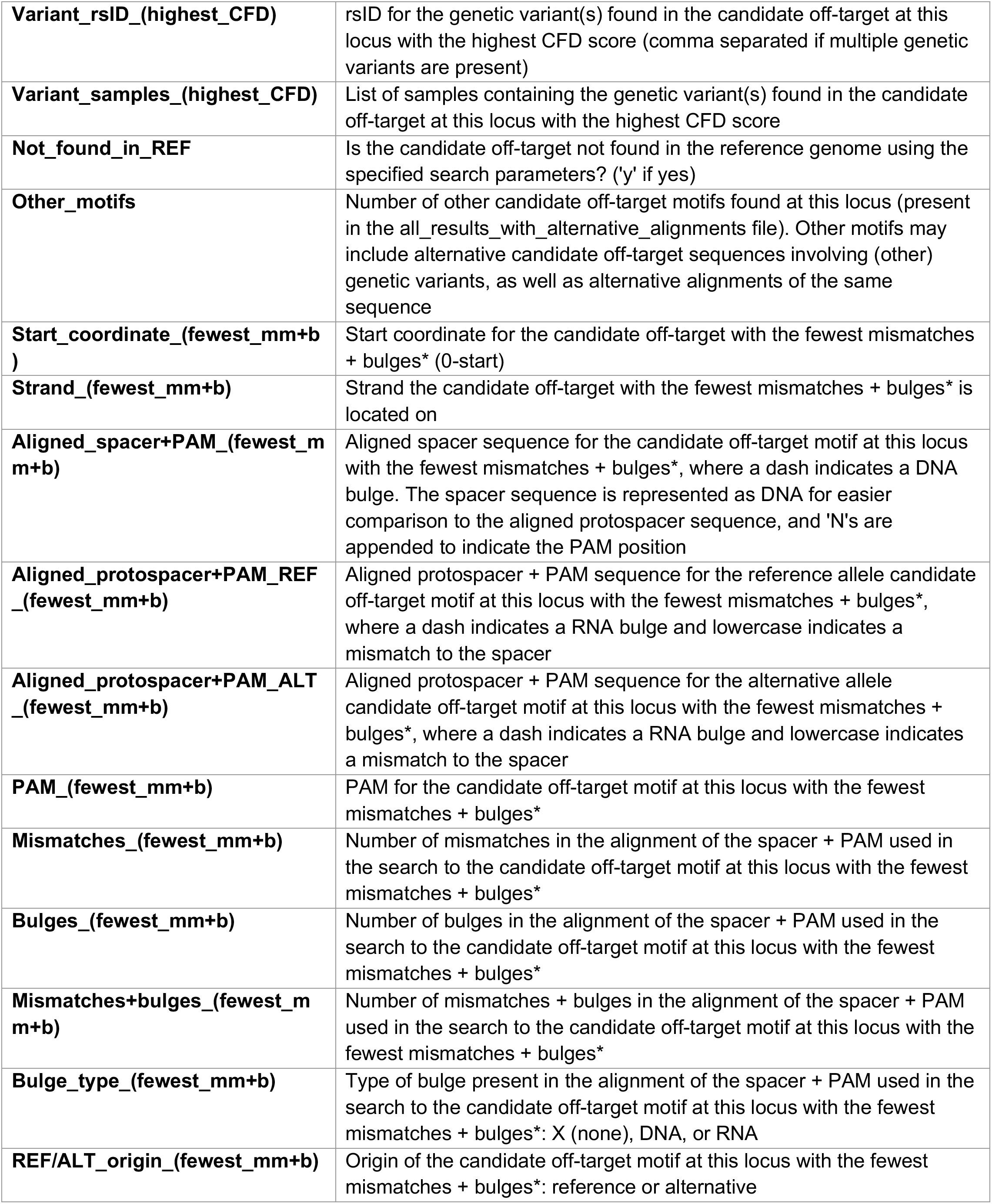
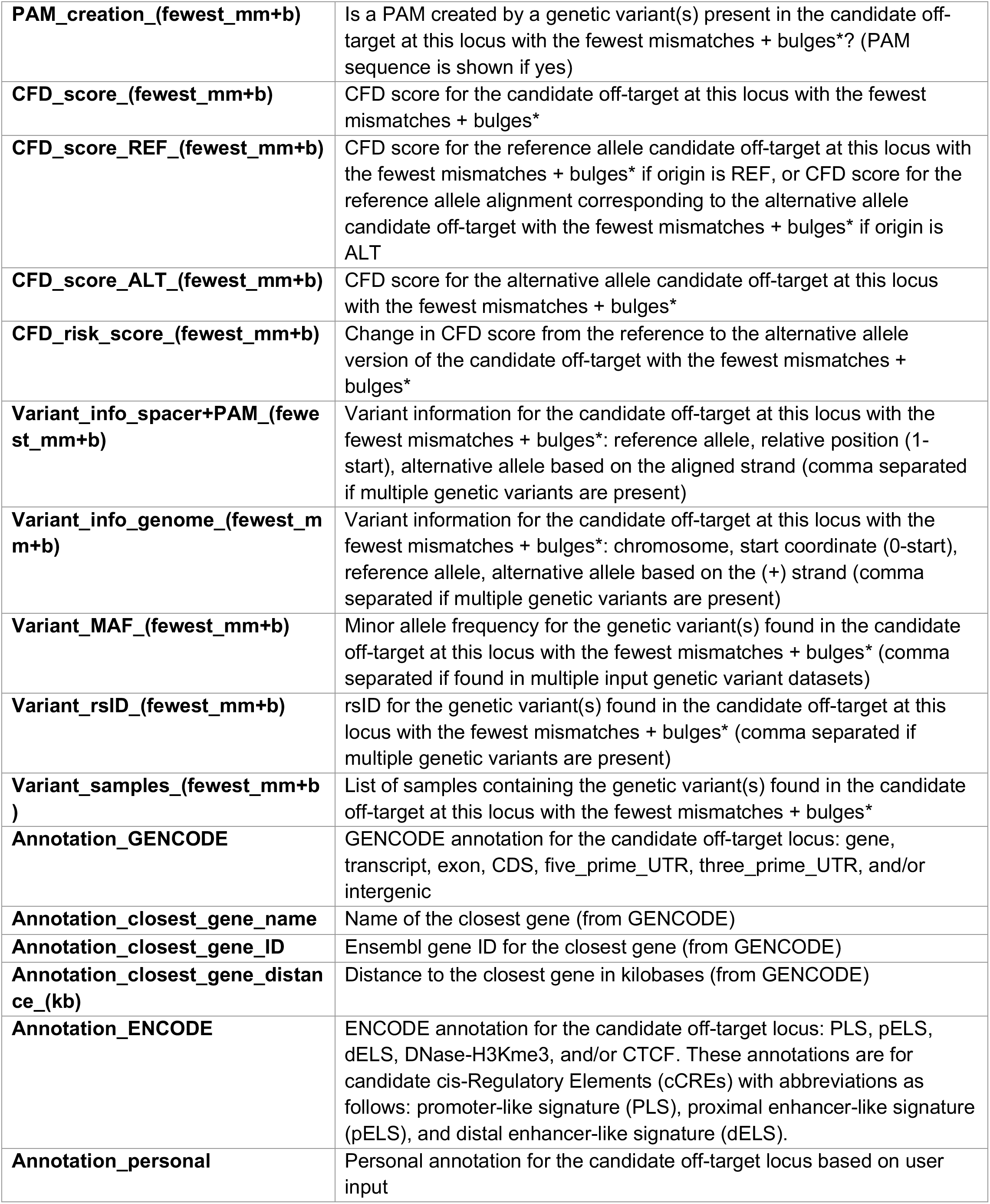
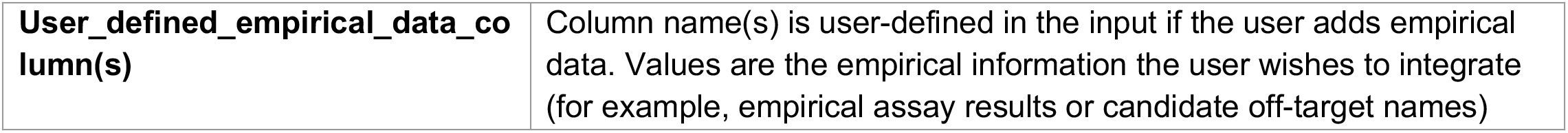
Table explaining columns found in the “integrated_results” file, “all_results_with_alternative_alignments” file and all user generated files from the webapp. CRISPRme reports all the possible alignments for any given genomic location (within 3 bp as default setting) meeting the user defined mismatches and bulges thresholds in the file “**all_results_with_alternative_alignments**”. If multiple alignments are possible at a given location for a candidate off-target (due to bulges or indel variants), CRISPRme will prioritize and report one best alignment each for highest CFD score, highest CRISTA score, and fewest mismatches plus bulges. The selected off-targets are reported in the integrated file “**integrated_results**” available for download from the website using the link “Download integrated results”(see Supplementary Figure 9). For non-SpCas9 editors (for which CFD and CRISTA scores do not apply), CFD and CRISTA scores are set to −1, so each best alignment is the one with the fewest mismatches plus bulges.

| Column name | Description |
| --- | --- |
| **Spacer+PAM** | Spacer(s) + PAM used in input to perform the off-target search on the genome |
| **Chromosome** | Chromosome the candidate off-target is found on |
| **Start_coordinate_(highest_CFD)** | Start coordinate for the candidate off-target at this locus with the highest CFD score (0-start) |
| **Strand_(highest_CFD)** | Strand the candidate off-target at this locus with the highest CFD score is located on |
| **Aligned_spacer+PAM_(highest_CFD)** | Aligned spacer sequence for the candidate off-target motif at this locus with the highest CFD score, where a dash indicates a DNA bulge. The spacer sequence is represented as DNA for easier comparison to the aligned protospacer sequence, and 'N's are appended to indicate the PAM position |
| **Aligned_protospacer+PAM_REF_(highest_CFD)** | Aligned protospacer + PAM sequence for the reference allele (version of) the candidate off-target motif at this locus with the highest CFD score, where a dash indicates a RNA bulge and lowercase indicates a mismatch to the spacer |
| **Aligned_protospacer+PAM_ALT_(highest_CFD)** | Aligned protospacer + PAM sequence for the alternative allele candidate off-target motif at this locus with the highest CFD score, where a dash indicates a RNA bulge and lowercase indicates a mismatch to the spacer |
| **PAM_(highest_CFD)** | PAM for the candidate off-target motif at this locus with the highest CFD score |
| **Mismatches_(highest_CFD)** | Number of mismatches in the alignment of the spacer + PAM used in the search to the candidate off-target motif at this locus with the highest CFD score |
| **Bulges_(highest_CFD)** | Number of bulges in the alignment of the spacer + PAM used in the search to the candidate off-target motif at this locus with the highest CFD score |
| **Mismatches+bulges_(highest_CFD)** | Number of mismatches + bulges in the alignment of the spacer + PAM used in the search to the candidate off-target motif at this locus with the highest CFD score |
| **Seed_mismatches+bulges_REF_(highest_CFD)** | Number of mismatches + bulges in the alignment of the spacer + PAM, counting only the occurrences happening in the half sequence nearest to the PAM location. Using the REF sequence. |
| **Non_seed_mismatches+bulges_REF_(highest_CFD)** | Number of mismatches + bulges in the alignment of the spacer + PAM, counting only the occurrences happening in the half sequence farthest to the PAM location. Using the REF sequence. |
| **Seed_mismatches+bulges_ALT_(highest_CFD)** | Number of mismatches + bulges in the alignment of the spacer + PAM, counting only the occurrences happening in the half sequence nearest to the PAM location. Using the ALT sequence. |
| **Non_seed_mismatches+bulges_ALT_(highest_CFD)** | Number of mismatches + bulges in the alignment of the spacer + PAM, counting only the occurrences happening in the half sequence farthest to the PAM location. Using the ALT sequence. |
| **Bulge_type_(highest_CFD)** | Type of bulge present in the alignment of the spacer + PAM used in the search to the candidate off-target motif at this locus with the highest CFD score: X (none), DNA, or RNA |
| **REF/ALT_origin_(highest_CFD)** | Origin of the candidate off-target at this locus with the highest CFD score: reference allele (REF) or alternative allele (ALT) |
| **PAM_creation_(highest_CFD)** | Is a PAM created by a genetic variant(s) present in the candidate off-target at this locus with the highest CFD score? (PAM sequence is shown if yes) |
| **CFD_score_(highest_CFD)** | Highest CFD score for the candidate off-target at this locus |
| **CFD_score_REF_(highest_CFD)** | Highest CFD score for the reference allele candidate off-target at this locus if origin is REF, or CFD score for the reference allele alignment corresponding to the alternative allele candidate off-target if origin is ALT |
| **CFD_score_ALT_(highest_CFD)** | Highest CFD score for the alternative allele candidate off-target at this locus |
| **CFD_risk_score_(highest_CFD)** | Change in CFD score from the reference to the alternative allele version of the candidate off-target with the highest CFD score |
| **Variant_info_spacer+PAM_(highest_CFD)** | Variant information for the candidate off-target at this locus with the highest CFD score: reference allele, relative position (1-start), alternative allele based on the aligned strand (comma separated if multiple genetic variants are present) |
| **Variant_info_genome_(highest_CFD)** | Variant information for the candidate off-target at this locus with the highest CFD score: chromosome, start coordinate (0-start), reference allele, alternative allele based on the (+) strand (comma separated if multiple genetic variants are present) |
| **Variant_MAF_(highest_CFD)** | Minor allele frequency for the genetic variant(s) found in the candidate off-target at this locus with the highest CFD score (comma separated if found in multiple input genetic variant datasets) |
| **Variant_rsID_(highest_CFD)** | rsID for the genetic variant(s) found in the candidate off-target at this locus with the highest CFD score (comma separated if multiple genetic variants are present) |
| **Variant_samples_(highest_CFD)** | List of samples containing the genetic variant(s) found in the candidate off-target at this locus with the highest CFD score |
| **Not_found_in_REF** | Is the candidate off-target not found in the reference genome using the specified search parameters? ('y' if yes) |
| **Other_motifs** | Number of other candidate off-target motifs found at this locus (present in the all_results_with_alternative_alignments file). Other motifs may include alternative candidate off-target sequences involving (other) genetic variants, as well as alternative alignments of the same sequence |
| **Start_coordinate_(fewest_mm+b )** | Start coordinate for the candidate off-target with the fewest mismatches + bulges* (0-start) |
| **Strand_(fewest_mm+b)** | Strand the candidate off-target with the fewest mismatches + bulges* is located on |
| **Aligned_spacer+PAM_(fewest_m m+b)** | Aligned spacer sequence for the candidate off-target motif at this locus with the fewest mismatches + bulges*, where a dash indicates a DNA bulge. The spacer sequence is represented as DNA for easier comparison to the aligned protospacer sequence, and 'N's are appended to indicate the PAM position |
| **Aligned_protospacer+PAM_REF_(fewest_mm+b)** | Aligned protospacer + PAM sequence for the reference allele candidate off-target motif at this locus with the fewest mismatches + bulges*, where a dash indicates a RNA bulge and lowercase indicates a mismatch to the spacer |
| **Aligned_protospacer+PAM_ALT_(fewest_mm+b)** | Aligned protospacer + PAM sequence for the alternative allele candidate off-target motif at this locus with the fewest mismatches + bulges*, where a dash indicates a RNA bulge and lowercase indicates a mismatch to the spacer |
| **PAM_(fewest_mm+b)** | PAM for the candidate off-target motif at this locus with the fewest mismatches + bulges* |
| **Mismatches_(fewest_mm+b)** | Number of mismatches in the alignment of the spacer + PAM used in the search to the candidate off-target motif at this locus with the fewest mismatches + bulges* |
| **Bulges_(fewest_mm+b)** | Number of bulges in the alignment of the spacer + PAM used in the search to the candidate off-target motif at this locus with the fewest mismatches + bulges* |
| **Mismatches+bulges_(fewest_m m+b)** | Number of mismatches + bulges in the alignment of the spacer + PAM used in the search to the candidate off-target motif at this locus with the fewest mismatches + bulges* |
| **Bulge_type_(fewest_mm+b)** | Type of bulge present in the alignment of the spacer + PAM used in the search to the candidate off-target motif at this locus with the fewest mismatches + bulges*: X (none), DNA, or RNA |
| **REF/ALT_origin_(fewest_mm+b)** | Origin of the candidate off-target motif at this locus with the fewest mismatches + bulges*: reference or alternative |
| **PAM_creation_(fewest_mm+b)** | Is a PAM created by a genetic variant(s) present in the candidate off-target at this locus with the fewest mismatches + bulges*? (PAM sequence is shown if yes) |
| **CFD_score_(fewest_mm+b)** | CFD score for the candidate off-target at this locus with the fewest mismatches + bulges* |
| **CFD_score_REF_(fewest_mm+b)** | CFD score for the reference allele candidate off-target at this locus with the fewest mismatches + bulges* if origin is REF, or CFD score for the reference allele alignment corresponding to the alternative allele candidate off-target with the fewest mismatches + bulges* if origin is ALT |
| **CFD_score_ALT_(fewest_mm+b)** | CFD score for the alternative allele candidate off-target at this locus with the fewest mismatches + bulges* |
| **CFD_risk_score_(fewest_mm+b)** | Change in CFD score from the reference to the alternative allele version of the candidate off-target with the fewest mismatches + bulges* |
| **Variant_info_spacer+PAM_(fewest_mm+b)** | Variant information for the candidate off-target at this locus with the fewest mismatches + bulges*: reference allele, relative position (1-start), alternative allele based on the aligned strand (comma separated if multiple genetic variants are present) |
| **Variant_info_genome_(fewest_mm+b)** | Variant information for the candidate off-target at this locus with the fewest mismatches + bulges*: chromosome, start coordinate (0-start), reference allele, alternative allele based on the (+) strand (comma separated if multiple genetic variants are present) |
| **Variant_MAF_(fewest_mm+b)** | Minor allele frequency for the genetic variant(s) found in the candidate off-target at this locus with the fewest mismatches + bulges* (comma separated if found in multiple input genetic variant datasets) |
| **Variant_rsID_(fewest_mm+b)** | rsID for the genetic variant(s) found in the candidate off-target at this locus with the fewest mismatches + bulges* (comma separated if multiple genetic variants are present) |
| **Variant_samples_(fewest_mm+b)** | List of samples containing the genetic variant(s) found in the candidate off-target at this locus with the fewest mismatches + bulges* |
| **Annotation_GENCODE** | GENCODE annotation for the candidate off-target locus: gene, transcript, exon, CDS, five_prime_UTR, three_prime_UTR, and/or intergenic |
| **Annotation_closest_gene_name** | Name of the closest gene (from GENCODE) |
| **Annotation_closest_gene_ID** | Ensembl gene ID for the closest gene (from GENCODE) |
| **Annotation_closest_gene_distance_(kb)** | Distance to the closest gene in kilobases (from GENCODE) |
| **Annotation_ENCODE** | ENCODE annotation for the candidate off-target locus: PLS, pELS, dELS, DNase-H3Kme3, and/or CTCF. These annotations are for candidate cis-Regulatory Elements (cCREs) with abbreviations as follows: promoter-like signature (PLS), proximal enhancer-like signature (pELS), and distal enhancer-like signature (dELS). |
| **Annotation_personal** | Personal annotation for the candidate off-target locus based on user input |
| **User_defined_empirical_data_column(s)** | Column name(s) is user-defined in the input if the user adds empirical data. Values are the empirical information the user wishes to integrate (for example, empirical assay results or candidate off-target names) |

### Supplementary Note 5. Search with custom personal genomes, VCFs, annotation files and PAMs

CRISPRme can be also deployed to any private server (**Supplementary Note 9**) and the offline version offers additional functionalities, including the option to input personal data (such as genetic variants, annotations, and/or empirical off-target results) as well as custom PAMs and genomes. There is no limit on the number of spacers, mismatches, and/or bulges used in the offline search. The required inputs are similar to the online version (see **Supplementary Note 1**) but the user can upload any personal data as long as they follow the format described below.

CRISPRme automatically creates the following folders to help organize the data that needs to be provided by the user. See Supplementary Figure 16 for an example.

• `Genomes:` it contains the genomes in FASTA format. Each genome must be saved into a separate folder. The name of the folder will be used to identify the genome itself and all the linked data such as VCFs and samplesIDs. In Supplementary Figure 16 the `Genomes` folder contains:
- `hg19:` FASTA files for human reference genome build 19.
- `hg38:` FASTA files for human reference genome build 38.
• `VCFs:` it contains the VCF datasets. Each dataset must consist of chromosome separated VCF files and be saved into a separate folder. The name of the folder is composed of the genome release name followed by the VCF dataset name. In Supplementary Figure 16 the `VCFs` folder contains:
- `hg38_HGDP`: VCF files from HGDP based on hg38 (ftp://ngs.sanger.ac.uk/production/hgdp/hgdp_wgs.20190516).
- `hg38_1000G`: VCF files from 1000G based on hg38 (ftp://ftp.1000genomes.ebi.ac.uk/vol1/ftp/data_collections/1000_genomes_project/release/20190312_biallel ic_SNV_and_INDEL/).
• `samplesIDs:` it contains the samplesID files, one for each VCF dataset. The name of the file is composed of the name of the corresponding VCF folder followed by the samplesID suffix. In Supplementary Figure 16 the `samplesIDs` folder contains:
- `hg38_HGDP.samplesID.txt`: tabulated file with header to identify samples for the HGDP dataset.
• `Annotations:` it contains the annotation BED files. In Supplementary Figure 16 the `Annotations` folder contains:
- `encode+gencode.hg38.bed`: a BED file with annotations from the GENCODE and ENCODE datasets.
• `PAMs:` it contains the PAM text files, one for each PAM. In Supplementary Figure 16 the `PAMs` folder contains:
- `20bp-NGG-SpCas9.txt`: a text file with a single line PAM sequence. The file name contains the position and length of the spacer (20 bp), followed by the PAM sequence (NGG) and the Cas protein (SpCas9).

**Supplementary Figure 16.**
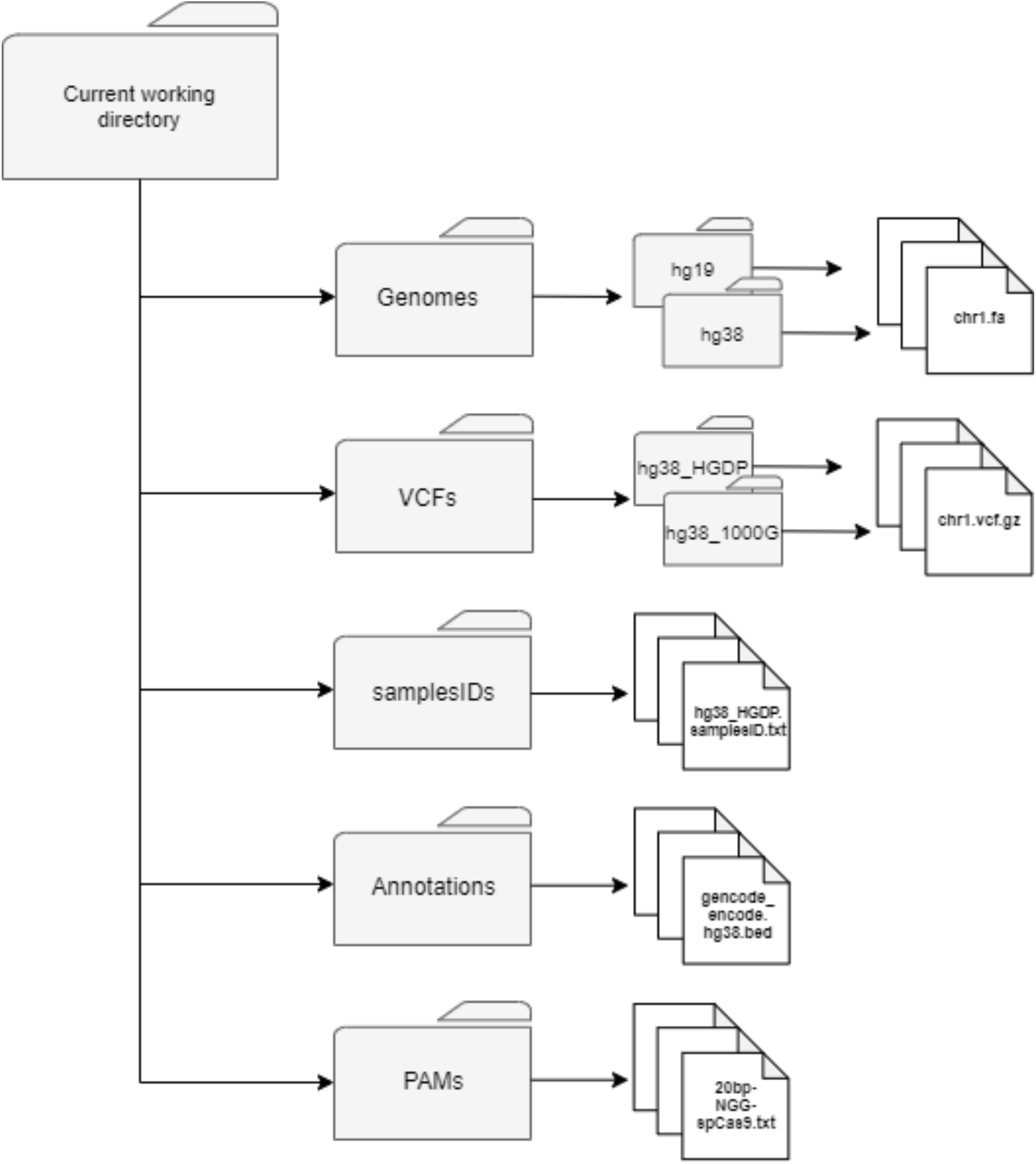
CRISPRme data storing structure. The directories created by CRISPRme running as an offline web-app or command line tool, used to upload personal data such as genomes, VCFs, annotations and PAMs.

The following sections detail the format of the required files in each folder.

#### Personal genome build

A personal genome can be added as a set of FASTA files (`.fa`), one for each chromosome (`chr1.fa, chr2.fa, chrN.fa`), all placed in a single folder. The supported format is based on the specifications for genome assemblies of the UCSC Genome Browser (e.g. hg38).

An example of a personal chromosome in FASTA format:

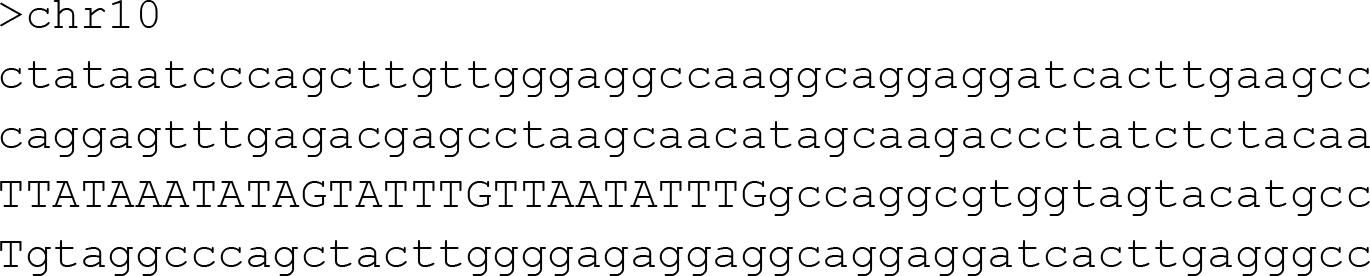

#### VCF files & phasing information

The Variant Call Format (VCF) file stores genetic variant information. CRISPRme accepts compressed VCFs in the 1000G or GATK v3/4 format (VCF v4.1 or newer) in chromosome separated files. VCF files containing variants from multiple chromosomes must be split by chromosome, which can be accomplished using BCFtools with the following command:

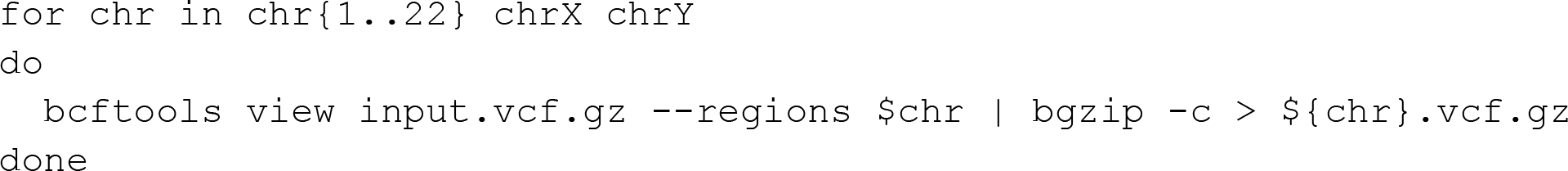

VCF files contain positional information (chromosome and position), reference and alternative nucleotide(s), and may contain sample genotype information (which, if present, can be either phased or unphased). A sample information file must be also provided for CRISPRme, i.e. a tabulated list containing all the samples present in the VCF files with their respective population (e.g. GBR), super-population (e.g. EUR) and sex (e.g. male) information. If the population, super-population, and/or sex information is not available, a placeholder such as ‘n’ can be used instead. VCF files from 1000G and HGDP are similar in format and report the same data. In 1000G VCFs, each sample column contains the phased genotype. In HGDP VCFs, each sample column contains the unphased genotype if available, along with supplementary data like the read depth and the genotype quality.

An example of VCF file header information from 1000G and HGDP:

• `#CHROM -` Chromosome
• `POS -` Position of the variant (1-start)
• `ID -` rsID or other identifier of the variant
• `REF -` Reference nucleotide
• `ALT -` Alternative nucleotide
• `QUAL -` Phred-scaled quality score for the assertion made in ALT
• `FILTER -` Testify if the ALT nucleotide passed quality filters. <u>Note that only variant calls that pass all</u> <u>quality filters (denoted with “PASS” in this field) are used for CRISPRme analysis.</u>
• `INFO -` a non-standard field containing details on the variants, including:
○ allele frequency
○ number of total samples
○ total alleles
○ any other possible information on the variants
• `FORMAT -` the format for the variant data reported in the subsequent columns
• `SAMPLE IDs -` a set of columns reporting the IDs of all the samples in the VCF

An example of the tab-delimited sample IDs text file needed:

| #SAMPLE_ID | POPULATION_ID | SUPERPOPULATION_ID | SEX |
| --- | --- | --- | --- |
| HG00096 | GBR | EUR | male |
| HG00097 | GBR | EUR | female |
| HG00098 | GBR | EUR | male |
| HG00099 | GBR | EUR | female |
| HG00100 | GBR | EUR | female |

• `#SAMPLE_ID -` sample identifier as reported in the VCF file header
• `POPULATION_ID -` population name as reported in the VCF file
• `SUPERPOPULATION_ID` – super-population name
• `SEX` – sex

In addition, CRISPRme supports gnomAD v3.1 VCFs based on an integrated parser. This process converts each gnomAD VCF into a CRISPRme-supported VCF. The parser takes as input a directory containing gnomAD v3.1 VCFs and a pre-generated samplesID file as shown in the following example. The pre-generated file simulates a set of samples, each one belonging to a gnomAD super-population, to be included in the gnomAD VCFs. The file is created by inspecting gnomAD v3.1 VCFs and extracting all the super-populations reported in the files (AFR, AMR, ASJ, EAS, FIN, NFE, MID, SAS and OTH). This file is provided with the test package (see **Supplementary Note 9**) and can be used and extended, if necessary, with any gnomAD v3.1 VCF file.

Example call of the VCF converter:

`crisprme.py gnomAD-converter --gnomAD_VCFdir VCFs/gnomAD_VCFdir/ --samplesID samplesIDs/hg38_gnomAD.samplesID.txt --thread 4`

Example of samples ID file to use with gnomAD v3.1 data:

| #SAMPLE_ID | POPULATION_ID | SUPERPOPULATION_ID | SEX |
| --- | --- | --- | --- |
| afr | AFR | AFR | n |
| ami | AMI | AMI | n |
| amr | AMR | AMR | n |
| asj | ASJ | ASJ | n |
| eas | EAS | EAS | n |
| fin | FIN | FIN | n |
| nfe | NFE | NFE | n |
| mid | MID | MID | n |
| sas | SAS | SAS | n |
| oth | OTH | OTH | n |

CRISPRme supports three types of VCF data:

1. <u>Individual-level, phased VCFs</u> (such as from 1000G) with genotypes for all samples, as well as information on which chromosome in a homologous pair each genotype call came from (delimited with ‘|’). When phase information is provided, CRISPRme is haplotype-aware and assesses off-target potential only for observed haplotypes.
2. <u>Individual-level, unphased VCFs</u> (such as from HGDP) with genotypes for all samples but lacking phase information (delimited with ‘/’). When individual-level information is provided but phase information is unavailable, CRISPRme assesses off-target potential for all possible haplotypes for each individual. However, this analysis may include false haplotypes in the case of nearby heterozygous variants present in the same individual.
3. <u>Population-level VCFs</u> (such as from gnomAD) with variant information for overall populations. CRISPRme can assess off-target potential when provided with many known population variants, but note that without individual-level and phase information, many unobserved haplotypes (with variants only observed in distinct individuals) may be included in the analysis.

Phase information prevents candidate off-targets arising from genetic variants that are not co-inherited from inflating the reported results. To demonstrate the value of phase information, Supplementary Figure 17 shows the results from Fig. 1 if CRISPRme were not haplotype-aware:

**Supplementary Figure 17.**
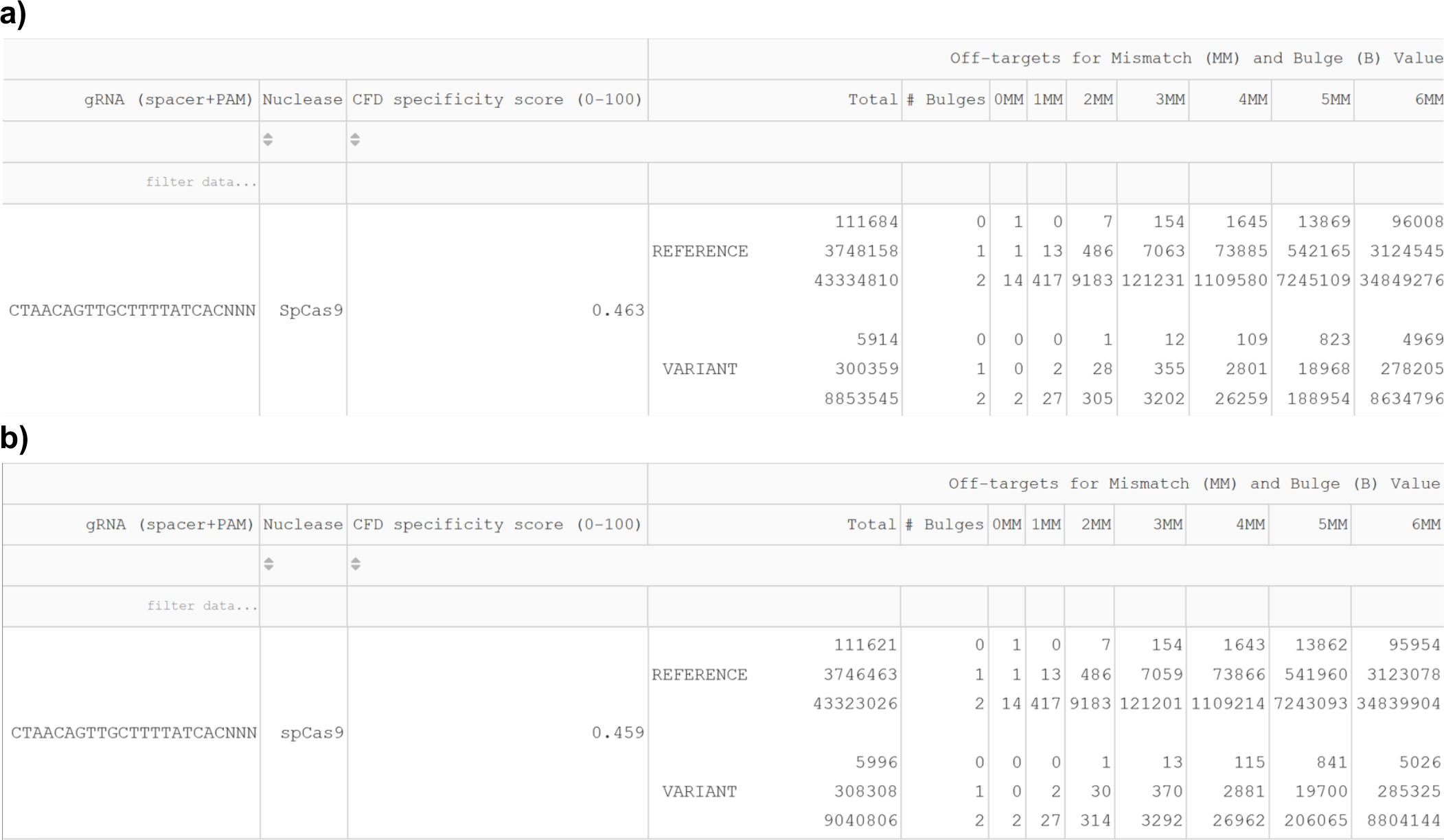
1000G VCFs results with vs. without phasing. **a)** Haplotype-aware CRISPRme results presented in Fig 1. **b)** CRISPRme results for the same search presented in Fig. 1 but without using phasing information. This results in an inflated number of candidate variant off-targets.

#### Custom annotation file, gene annotations and empirical off-target data

The offline version of CRISPRme can take in a custom annotation file in BED format and annotate the potential off-targets. This file can be used independently or in combination with the provided ENCODE and GENCODE annotation files. If an off-target is associated with multiple annotations, each one will be reported. The annotation file containing ENCODE+GENCODE data was created extracting data from ENCODE cCREs (https://api.wenglab.org/screen_v13/fdownloads/GRCh38-ccREs.bed) and the GENCODE v35 comprehensive gene annotation (“ALL” regions) file (http://ftp.ebi.ac.uk/pub/databases/gencode/Gencode_human/release_35/gencode.v35.chr_patch_hapl_scaff.b <u>asic.annotation.gff3.gz</u>).

An example annotation file, where the columns indicate the chromosome, the genomic start and end coordinates and the annotation description:

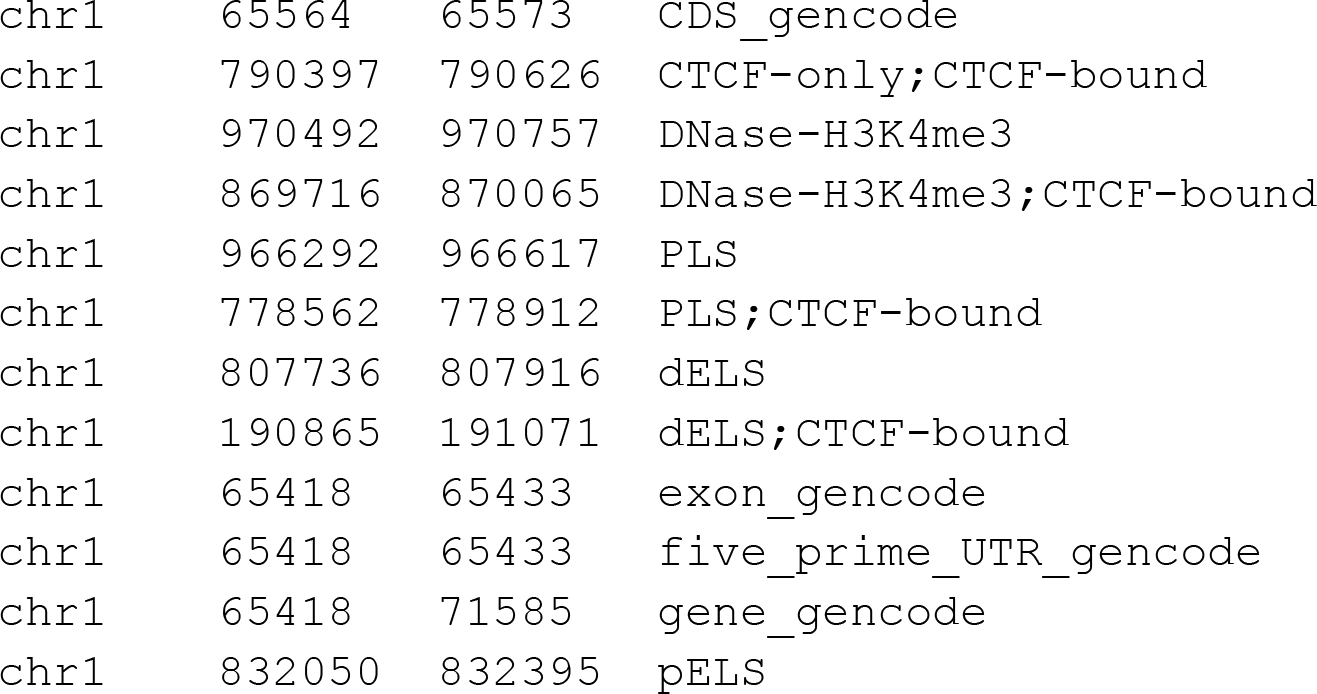
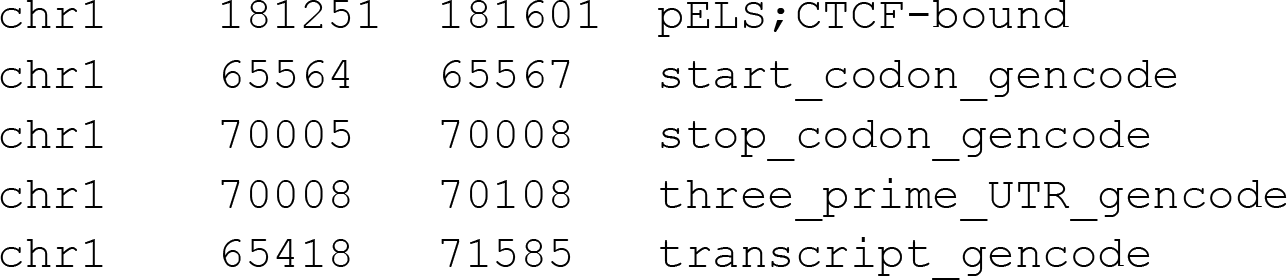

The coordinates in the custom annotation file are 0-start based to respect the BED format specifications (https://genome.ucsc.edu/FAQ/FAQformat.html#format1).

CRISPRme can also integrate information on the nearest gene to each candidate off-target. An example GENCODE file where the columns are chromosome, start position, end position, annotation information, strand, origin database, and annotation. Other columns are not shown for space reasons:

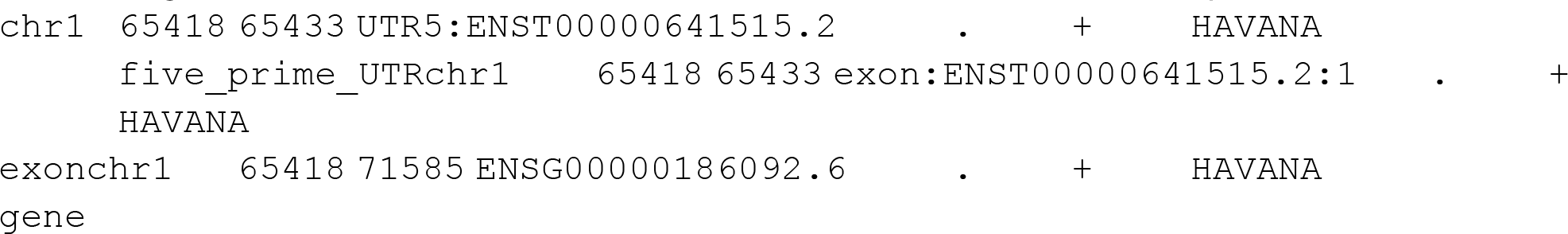

The command line version of CRISPRme also supports the integration of user-provided empirical off-target results, which can be useful for creating a master summary of candidate off-target sites nominated by in silico, in vitro and cellular methods. Below is an example of the input format, with the data representing several previously nominated off-target sites for gRNA #1617.

An example of empirical off-target data for integration:

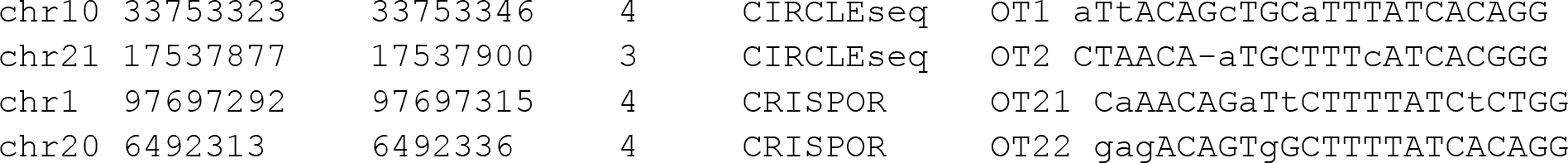

In order, the columns indicate chromosome, start position, end position, number of mismatches + bulges, user-defined column name for the CRISPRme output files, empirical information to integrate and off-target motif (including PAM). If the motif is not available, it can be substituted with any placeholder (‘n’ for example). The empirical information to integrate can be anything the user desires, such as other identifiers (as shown here in column 6) or numerical scores.

#### PAM sequence

In order to use a new PAM, the user must add a new file in the PAMs folder with the following naming convention:

*##bp_protospacer-PAM_seq-nuclease.txt* if the PAM is located at the 3’ end of the protospacer

*PAM_seq-##bp_protospacer-nuclease.txt* if the PAM is located at the 5’ end of the protospacer

The content of this file must consist of a series of Ns representing the protospacer and the actual PAM sequence immediately preceding or following it as appropriate. Then, after a whitespace, there must be an integer representing the length of the PAM sequence. E.g. if the PAM considered is NGG → 3. If the PAM sequence is located 5’ of the protospacer, then this value must be negative. E.g. if the PAM considered is TTTV → −4.

An example of a PAM file (NGG for SpCas9) with a protospacer length of 20 nt:

`20bp-NGG-spCas9.txt`

NNNNNNNNNNNNNNNNNNNNNGG 3

### Supplementary Note 6. Comparison of CRISPRme with available tools

Although numerous tools are available to enumerate CRISPR-Cas off-targets, to our knowledge only three previous studies^4–6^ have reported computational strategies to assess off-target potential in the presence of genetic variants. However, only crispRtool from Lessard et al and CALITAS from Fennel et al^6^ provide general command line software. Upon testing, CALITAS (v1.0) cannot use phasing information and doesn’t support directly the publicly available 1000 Genomes Project VCFs as input thereby limiting the utility of the tool for generalizable variant-aware analysis.

Therefore, we decided to focus our comparison of CRISPRme (v1.7.7) only with crisprRtool (v2.0.5). We considered available features and running times on the same hardware (AMD Ryzen Threadripper 3970X 32-Core Processor clocked at 2.2 GHz with 124 GB RAM) to provide a fair assessment. For our tests we used the 1617 sgRNA, NGG PAM, variants from 1000G and a variable number of mismatches and bulges.

Briefly, crisprRtool first adds variants (SNPs only) to the reference genome using IUPAC notation, and then searches the input gRNA(s) on the variant genome and reports a list of putative on-and off-targets with IUPAC nucleotides. The tool also offers the possibility to search each VCF file individually to resolve haplotypes (SNPs and INDELs) of the reported off-targets. However, for this step, the user needs to manually edit and execute a script for each VCF file. In addition, the search operation with crispRtool allows a maximum of 5 mismatches, does not account for bulges, and is not flexible in terms of PAM location relative to the protospacer (only 3’ is supported).

Using 5 mismatches and the settings described above, crispRtool took 9 hours to complete the non-haplotype-resolved search. The haplotype-resolved search only on chr1 using variants from 1000G (6 million SNPs and INDEL variants) took ∼37 hours. Conservatively extrapolating to all other chromosomes, the entire search would take more than 300 hours and will not be as complete as the search CRISPRme offers due to the lack of graphical reports and textual summaries encompassing results from all chromosomes.

On the other hand, by leveraging an efficient genome index and auxiliary data structures that are constructed only once during the installation (∼4 hours for NGG PAM, ∼12 hours for NNN PAM or can be downloaded directly in our complete test package), CRISPRme can complete a haplotype-aware search for a gRNA across the entire genome with 5 mismatches in ∼1 hour. The haplotype-resolved search on the entire genome with up to 6 mismatches and 2 DNA/RNA bulges only takes 2 hours (excluding the guide-independent indexing operation described above) and also includes a summary report.

### Supplementary Note 7. Assaying variant-aware off-target potential

**Supplementary Table 2.** Comparison of off-target assessment in studies involving sg1617.

| Study | Type | Search performed (method/tool if known) | Genome used | Sequencing validation results |
| --- | --- | --- | --- | --- |
| Wu et al. *Nat Med* 2019<sup>7</sup> | Pre-clinical | In vitro (CIRCLE-seq) & in silico with NGG PAM (CRISPOR) | Reference / donor DNA | No off-targets identified with $\geq 0.1\%$ indels (tested 24) |
| Frangoul et al. *NEJM* 2021 | Clinical trial | In cellulo (GUIDE-seq) & in silico search with up to 3 mismatches or 2 mismatches + 1 bulge and NNG, NGN, or NAA PAM | Reference / donor DNA | No off-targets identified with $\geq 0.2\%$ indels (tested 223) |
| This manuscript | | In silico search with up to 6 mismatches + 2 bulges and NNN PAM (CRISPRme) | Reference + 1000 Genomes Project variants | Allele-specific off-target identified with $\sim 10\%$ indels |

**Supplementary Table 3.**
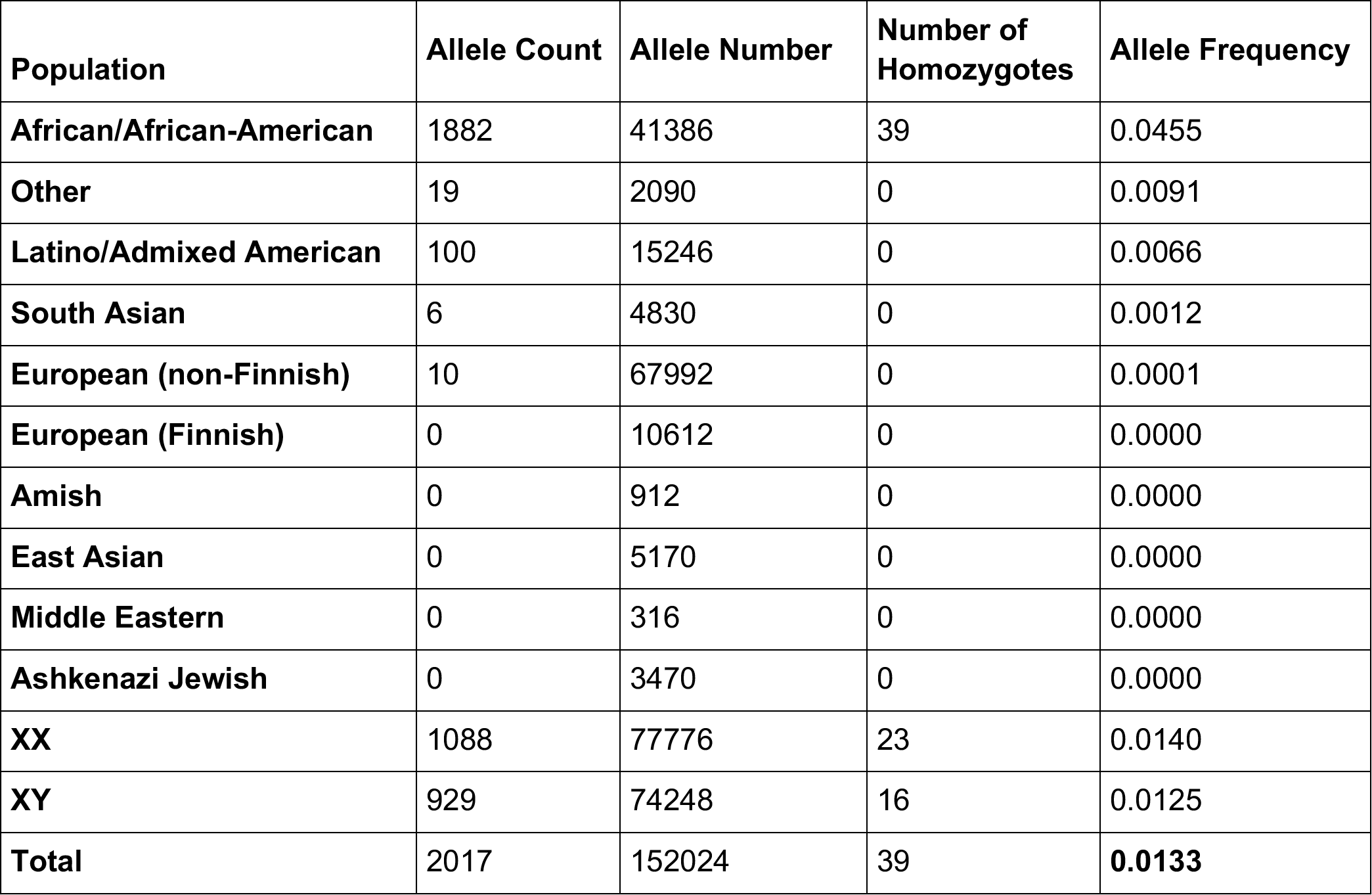
Complete population frequencies for rs114518452 from gnomAD v3.1.

| Population | Allele Count | Allele Number | Number of Homozygotes | Allele Frequency |
| --- | --- | --- | --- | --- |
| **African/African-American** | 1882 | 41386 | 39 | 0.0455 |
| **Other** | 19 | 2090 | 0 | 0.0091 |
| **Latino/Admixed American** | 100 | 15246 | 0 | 0.0066 |
| **South Asian** | 6 | 4830 | 0 | 0.0012 |
| **European (non-Finnish)** | 10 | 67992 | 0 | 0.0001 |
| **European (Finnish)** | 0 | 10612 | 0 | 0.0000 |
| **Amish** | 0 | 912 | 0 | 0.0000 |
| **East Asian** | 0 | 5170 | 0 | 0.0000 |
| **Middle Eastern** | 0 | 316 | 0 | 0.0000 |
| **Ashkenazi Jewish** | 0 | 3470 | 0 | 0.0000 |
| **XX** | 1088 | 77776 | 23 | 0.0140 |
| **XY** | 929 | 74248 | 16 | 0.0125 |
| **Total** | 2017 | 152024 | 39 | **0.0133** |

By design, genomic DNA based off-target detection methods (like cell-based GUIDE-seq or in vitro Digenome-seq and CIRCLE-seq^8–10^) test a small number of available donors. These assays are valuable tools to identify reference or very common variant-associated off-targets but are not scalable to evaluate a wide breadth of variant off-target sites which would require obtaining material from and testing innumerable donors of diverse genotypes. In contrast, massively parallel assays with libraries of synthetic oligonucleotide homologous targets including naturally occurring or synthetic variants can evaluate the potential of a given Cas:gRNA RNP to cleave a given target, testing many possible variant off-targets in parallel in a single assay^11, 12^. We would propose using these methods in conjunction with CRISPRme, first running CRISPRme to nominate all potential sites by homology and then to prioritize those for experimental testing based on both computational and experimental off-target prediction methods and existing annotations. Nonetheless, in vitro assays of oligonucleotide targets may imperfectly correlate with genetic modification in a relevant delivery and cellular context. When possible, we suggest editing cells bearing the variant genotype of interest and sequencing the endogenous variant site. Ex vivo edited patient cells could be tested by amplicon sequencing prior to infusion, although the functional importance of off-target edits may range from likely functional to likely neutral, so the mere presence of off-target editing in a cell product may not necessarily preclude its clinical use. For those edits that are likely neutral, ongoing monitoring of patient samples (somewhat analogous to vector integration site monitoring in gene therapy protocols) could help assess patient-specific risk, and also serve as valuable information for the broader field as to the frequency and in vivo dynamics of off-target edits. Our results highlight that variant off-target editing potential is not equally distributed across all ancestral groups but especially concentrated in those of African ancestry. Therefore gene editing efforts that include subjects of African ancestry (like those targeting sickle cell disease) might pay particular attention to this issue. Gene editing efforts that focus on a patient population should consider genetic variants enriched in that population in the off-target evaluation. However, our analysis also shows that variant off-targets may be private to a given individual, so all humans could potentially be susceptible to this kind of effect. Genetic variants may either increase or reduce off-target potential compared to a reference genome. In the case of variant-reduced off-target potential, off-target editing of the reference alleles might remain undesirable for a therapy targeting an admixed patient population. Overall, we suggest that a risk-assessment of variant off-target editing potential should be performed before clinical implementation, and ultimately to consider screening for variants associated with heightened risk in patient selection and monitoring of subjects for possible off-target effects after dosing.

### Supplementary Note 8. Candidate alternative allele-specific off-targets associated with therapeutic genome editing approaches

Besides the study using sg1617, several other clinical trials of ex vivo SpCas9 gene editing have been reported, such as one using a gRNA targeting *TRBC* for gene-edited CAR-T cells^13^. For this gRNA, 13 of the top 20 ranked off-target sites by CFD score are based on alternative alleles. One off-target site due to a rare variant with CFD_ref_ 0.05 and CFD_alt_ 0.5 is in coding sequences of *TAS2R10*, encoding a G-protein coupled receptor expressed in T-cells. A recent trial described in vivo SpCas9 gene editing with lipid nanoparticle delivery targeting the *TTR* gene in hepatocytes^14^. An off-target site for this gRNA has CFD_ref_ 0.01 and CFD_alt_ 0.65 that resides in an ENCODE candidate cis-regulatory element.

CRISPRme analysis is flexible to PAM sequences besides NGG. For base editors in particular, several reports have explored SpCas9 variants with non-NGG PAM restriction to optimally position the desired edit within the base editing window. For example, one report^15^ described converting the *HBB* sickle cell mutation to a benign hemoglobin variant Makassar using A-to-G base editing with a NGC PAM restricted SpCas9 variant and an IBE12 deaminase architecture with base editing window from protospacer position 2-16 centered around position 9. Several alternative allele nominated off-target sites overlap coding sequences, including one with 4 mismatches + bulges to reference genome but only 3 mismatches + bulges in the presence of an alternative allele (MAF 0.0007%) within *ARHGAP26* where base edits in the center of the editing window would be predicted to produce missense mutations. *AHRGAP26* has been described as a tumor suppressor gene^16^.

Another report described using the ABE8e-NRCH base editor to convert the sickle cell allele to the Makassar variant. CRISPRme analysis of alternative allele off-targets showed 1609 alternative allele off-targets resulting in 4 or fewer MM+B including 198 PAM creation off-targets. One alternative allele off-target (MAF 0.005%) overlapped the coding sequence for *PLA2G4B*, where edits in the base editing window would be expected to product splice acceptor site disruption loss-of-function alleles. PLA2G4B function has been linked to monocyte metabolic fitness^17^.

CRISPRme is also able to predict off-targets for Cas12a associated gRNAs with a 5’ TTTV PAM sequence. For example, for a Cas12a gRNA targeting the *HBG* promoter to treat SCD in current clinical trials (NCT04853576)^18^, CRISPRme identified an alternative allele specific off-target where the alternative allele creates the TTTV PAM and produces a protospacer with just 1 mismatch in the seed sequence that targets in an ENCODE candidate cis-regulatory element.

In summary, we show that CRISPRme enables comprehensive off-target nomination for a variety of guide RNAs in clinical development and for a number of Cas proteins (Supplementary Table 4, **Supplementary Files 1-3**). None of these alternative allele off-target sites appear to have been evaluated in the published descriptions of these gene editing approaches, thereby underscoring the utility of a tool like CRISPRme that can nominate candidate off-targets absent from the human reference genome but present in specific populations or individual patients for more comprehensive safety assessment.

**Supplementary Table 4.**
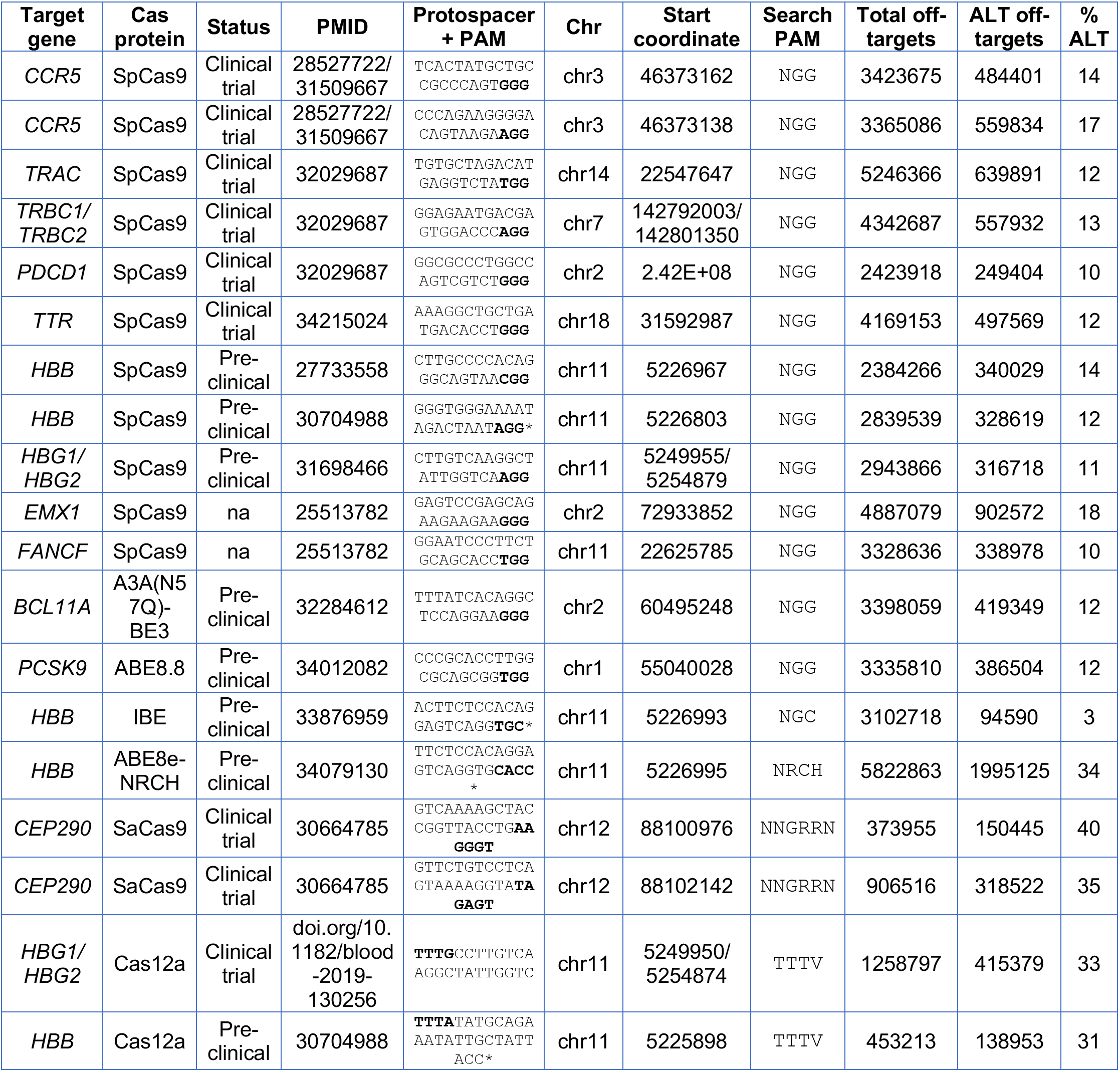
Additional gRNAs analyzed by CRISPRme representing a variety of target sequences, Cas proteins, and PAMs. These gRNAs are in clinical or preclinical development and/or widely tested in off-target studies. PAMs are indicated in bold. * indicates that the gRNA is specific to the sickle cell mutation (on-target site not found in hg38/REF). CRISPRme analysis was performed using the 1000G and HGDP datasets with up to 6 mismatches and 2 bulges.

### Supplementary Note 9. Details of CRISPRme installation and usage

#### Installation

CRISPRme can be used offline and is available as a comprehensive Conda package containing:

1. A command line version of the software and
2. 2. A web based tool with a graphical interface accessible from a local browser.

To install the package:

- Follow this link and install Conda: https://docs.conda.io/en/latest/miniconda.html
- After the installation is complete, add channels to the Conda local installation by typing the following commands into the terminal: `conda config --add channels defaults` `conda config --add channels bioconda` `conda config --add channels conda-forge` `conda install python=3.8 -y`
- Type into the terminal: `conda install crisprme -y`

The necessary inputs can be downloaded in one of two ways depending on user preference:

1. Download a small test package and desired inputs (genome, genetic variants, etc) directly from source: `gdown` https://drive.google.com/uc?id=11wn9pg6AWzDYZ7V_LjBIjGvgx95bnVJ1 `tar -xvf crisprme_test.tar.gz` `cd crisprme_test/` `bash crisprme_auto_test_conda.sh` The user can control which files are downloaded based on their desired use case by commenting out or uncommenting parts of `crisprme_auto_test_conda.sh.`
2. Download all the inputs presented in this manuscript (hg38, 1000G and HGDP variant datasets, annotations, and PAMs) along with precomputed indexes (to allow for faster processing): `crisprme_test_complete_package.sh` This will require ∼0.5 TB of disk space. After this command, CRISPRme will be ready to perform the searches presented in Fig. 1 and Fig. 2.

To allow use on any platform, CRISPRme is also available as a Docker image:

- Download the latest available CRISPRme image using the command `docker pull pinellolab/crisprme`
- After the pull is complete, a new Docker container can be created any time starting from the clean image

A complete installation guide is available at https://github.com/pinellolab/CRISPRme.

#### Usage

CRISPRme offline can be run as a web-app or a command line tool. The required input files are identical for the two versions (web-app and command line) and are described below and in **Supplementary Note 1**.

##### Web-app

1. Execute the command into the terminal: `crisprme.py web-interface`
2. Open a web browser and visit `127.0.0.1:8080.` The homepage of CRISPRme will open. If a remote server is used to host CRISPRme, input the IP address of the server in the web browser, e.g. `192.1.2.3:8080`, and the home page will open in the browser.
3. Now directly input data and select how to perform the search.

##### Command line

1. Write the following command into the terminal: `crisprme.py complete-search --help`
2. The above command will show all the mandatory and optional inputs for a CRISPRme complete-search. Please see **Supplementary Note 5** for the expected formats of these inputs.
  a. Mandatory input:

i. `--genome`, the reference genome folder
ii. `--guide`, the file that contains the guide(s) to use for the search
iii. `--pam`, the file containing the PAM to use for the search
iv. `--mm`, the maximum number of mismatches
v. `--bMax`, the maximum number of bulge
vi. `--output`, the output folder for the results
  b. Optional input:

i. `--vcf`, the file with the list of VCF folders to be used
ii. `--samplesID`, the file with the list of sample ID files (must have the same number of lines as the file passed to `--vcf`)
iii. `--annotation`, an ENCODE+GENCODE or custom file containing annotations for the reference genome
iv. `--gene_annotation`, a GENCODE or custom file to find the nearest gene for each candidate off-target
v. `--personal_annotation`, the file containing personal annotations for the reference genome
vi. `--bDNA`, the number of DNA bulges permitted in the search phase
vii. `--bRNA`, the number of RNA bulges permitted in the search phase
viii. `--merge`, the window to merge nearby off-targets (based on genomic position), using the off-target with the highest CFD score as the pivot [default 3]
ix. `--thread`, the number of threads to use in the search [default all available minus 2]
  c. Example call: crisprme.py complete-search --genome Genomes/hg38/ --vcf list_vcf.txt -- guide sg1617.txt --pam PAMs/20bp-NGG-SpCas9.txt --annotation Annotations/encode+gencode.hg38.bed --samplesID list_samplesID.txt -- gene_annotation Annotations/gencode.protein_coding.bed --bMax 2 --mm 6 -- bDNA 2 --bRNA 2 --merge 3 --output sg1617.6.2.2 --thread 4

Other CRISPRme functions:

1. Targets integration function, which can be used to generate an integrated file (see **Supplementary Note 4**) with user-defined empirical data. Command: `crisprme.py targets-integration` Input: a. `--targets,` bestMerge file (this file is generated automatically by the software) b. `--empirical_data`, empirical data file to integrate c. `--output`, output folder for the results
2. gnomAD converter function, which can be used to convert gnomADv3.1 VCFs to CRISPRme compatible VCFs (see **Supplementary Note 5**). Command: `crisprme.py gnomAD-converter` Input: a. `--gnomAD_VCFdir`, used to specify the directory containing gnomADv3.1 original VCFs b. `--samplesID`, used to specify the pre-generated samplesID file necessary to introduce samples into the gnomAD VCFs c. `--thread`, the number of threads to use in the process [default all available minus 2]
3. Generate personal card function, which can be used to generate a personal card for a specific sample (see **Supplementary Note 3**). Command: `crisprme.py generate-personal-card` Input: a. `--result_dir,` used to specify the directory containing results from a complete search b. `--guide_seq`, used to specify a guide + PAM sequence from the input set (e.g. CTAACAGTTGCTTTTATCACNNN) c. `--sample_id`, used to specify the ID of the sample to generate the personal card for (e.g. HGDP01211)
4. Web interface function, which can be used to start the web server necessary to host CRISPRme web-app locally. Command: `crisprme.py web-interface` No input is required for this function.

A more comprehensive explanation of CRISPRme functions can be found on GitHub at https://github.com/pinellolab/CRISPRme.

### Output files

CRISPRme outputs various results files:

- `Spacer+PAM_genome+VCFs_mismatches+bulges_integrated_results.tsv` contains comprehensive information on the candidate off-targets identified by CRISPRme, sorted by CFD score (see **Supplementary Note 4**).
- `Spacer+PAM_genome+VCFs_mismatches+bulges_all_results_with_alternative_alignment s.tsv` contains off-targets with alternative alignments and/or alleles that are found in the same genomic regions as those reported in the integrated_results file (see **Supplementary Note 4**)
- `jobID.summary_by_guide.GUIDE.txt` reports the number of candidate off-targets from reference, variant, combined (reference + variant), and corresponding to PAM creation events separated by bulge type and number of mismatches and bulges (see Supplementary Figure 11a).
- `jobID.summary_by_samples.GUIDE.txt` reports the number of personal candidate off-targets and PAM creation events for each sample (see Supplementary Figure 12a).
− `log.txt` contains the report of started and completed steps for the analysis process.
− `log_error.txt` contains the report of any errors encountered during the analysis.
− `log_verbose.txt` contains the verbose log of each step executed during the analysis.
− `crispritz_targets` directory: contains intermediate files generated by CRISPRitz.
- `refgenome_pam_guides.targets.txt` file containing all the reference targets generated from the search using CRISPRitz on the reference genome but without samples, scores, annotations and phasing information.
- `refgenome+vargenome_pam_guides.targets.txt` file containing all the variant targets generated from the search using CRISPRitz on the genome with variants encoded as IUPAC nucleotides but without samples, scores, annotations and phasing information.
- `indels_refgenome+vargenome_pam_guides.targets.txt` file containing all the indel targets generated from the search using CRISPRitz but without samples, scores, annotations and phasing information.
− `imgs` directory: contains all the images produced.
- `CRISPRme_top_1000_log_for_main_text_GUIDE.png` shows a visual summary of the top 1000 candidate off-targets by CFD score (see Fig. 1c **and** Supplementary Figure 14a).
- `populations_distribution_GUIDE_#total.png` shows the population distribution for each combination of mismatches + bulges (see Supplementary Figure 14b).
- `Summary_single_guide_GUIDE_#mm_#bul_TOTAL.png` shows the radar chart, table and motif logo of the examined guide for a specific combination of mismatches and bulges (see Supplementary Figure 14b). Other files necessary for computation and run-time queries are hidden from the user, but can be checked if desired and used for downstream analyses:
- `jobID.general_target_count.GUIDE.txt` contains a table showing the count of candidate off-targets grouped by number of mismatches and bulges (see Fig. 1b and **Supplementary Figure**).
- `jobID.acfd.txt` contains the CFD specificity score for each guide used in the search (see **Supplementary Note 4).**
- `jobID.PopulationDistribution.txt` contains the counts necessary to produce bar plots reporting how many off-targets belong to each superpopulation in each subcategory of mismatches + bulges.
- `jobID.bestMerge.txt` contains the candidate off-targets, with only the off-target having the highest CFD score within each merge window included to reduce the number of nearly redundant off-targets in the results. This intermediate file is used to generated the integrated_results file
- `jobID.bestMerge.txt.empirical_not_found.tsv` contains the reported empirical targets (if in input) that are were not associated to an in-silico predicted target.
- `Params.txt` contains the parameters of the search.
− `guides.txt` contains the list of guide(s) used in the search.
− `sampleID.txt` contains all the sampleIDs for each dataset used in the search.
− `guide_dict_GUIDE.json`, contains the dictionary to generate radar charts in real time (see Supplementary Figure 14b).
− `motif_dict_GUIDE.json`, contains the dictionary to generate motif plots in real time (see Supplementary Figure 14b).
− `jobID.db` is a SQL database created to perform real time queries on the integrated_results file (see **Supplementary File 1).**
− Files generated from user requests in the webapp containing candidate off-targets extracted from the integrated_results file using different criteria

### Supplementary Note 10. Experimental methods

#### Cell culture

Fresh G-CSF mobilized peripheral blood cells from healthy donor 1 were obtained from Miltenyi Biotec (Auburn, CA). CD34+ HSPCs were isolated using CliniMACS® CD34 reagent (Miltenyi, 130-017-501). Cryopreserved human CD34+ HSPCs from mobilized peripheral blood of deidentified healthy donors 2-7 were obtained from the Fred Hutchinson Cancer Research Center (Seattle, Washington). CD34+ HSPCs were cultured into Stem Cell Growth Medium (SCGM) (CellGenix, 20806-0500) supplemented with 100 ng ml ^-1^ human Stem Cell Growth Factor (SCF) (CellGenix, 1418-050), 100 ng ml ^-1^ human thrombopoietin (TPO) (CellGenix, 1417-050) and 100 ng ml ^-1^ recombinant human FMS-like Tyrosine Kinase 3 Ligand (Flt3-L) (CellGenix cat# 1415-050). HSPCs were electroporated with 3xNLS-SpCas9:sg1617 RNP or HiFi-3xNLS-SpCas9:sg1617 RNP 24 h after thawing. Twenty-four hours after electroporation, HSPCs were transferred into erythroid differentiation medium (EDM) consisting of IMDM (LIFE, 12440061) supplemented with 330 μg ml ^-1^ holo-human transferrin (Sigma, T0665-1G), 10 μg ml ^-1^ recombinant human insulin (Sigma, 19278-5ML), 2 IU ml^-1^ heparin (Sigma, H3149), 5% human solvent detergent pooled plasma AB (Rhode Island Blood Center), and 3 IU ml ^-1^ erythropoietin (Pharmacy). Five days after electroporation, cells were harvested for gDNA extraction.

#### Protein purification

3xNLS-SpCas9 was purified as previously described. HiFi-3xNLS-SpCas9 plasmids were transformed into BL21 (DE3) competent cells (MilliporeSigma, 702353) and grown in Terrific Broth (TB) media at 37°C until OD600 2.4-2.8. Cells were induced with 0.5 mM isopropyl ß-d-1-thiogalactopyranoside (IPTG) per liter for 20 hours at 20°C. Pellets were lysed in 25 mM Tris, pH 7.6, 500 mM NaCl, 5% glycerol, passed through homogenizer twice and centrifuged at 20,000 rpm for 1 hour at 4°C. Proteins were purified by Nickel-NTA resin and treated with TEV protease (1 mg lab made TEV per 40 mg of protein) and benzonase (100 units ml ^-1^, Novagen 70664-3) overnight at 4°C. Subsequently, the proteins were purified by size exclusion column (Amersham Biosciences HiLoad 26/60 Superdex 200 17-1071-01) and ion exchange with a 5 ml SP HP column (GE 17-1151-01) according to the manufacturer’s instructions. Proteins were dialyzed in 20 mM Hepes buffer pH 7.5 containing 400 mM KCl, 10% glycerol, and 1 mM TCEP buffer, and contaminants were removed by Toxin Sensor Chromogenic LAL Endotoxin Assay Kit (GenScript, L00350). Purified proteins were concentrated and filtered using Amicon ultra filter units – 30k NMWL (MilliporeSigma, UFC903008) and ultrafree-MC centrifugal filter (MilliporeSigma, UFC30GV0S). Protein fractions were further assessed on TGX stain free 4-20% SDS-PAGE (Biorad, 5678093) and quantified by BCA assay.

#### RNP electroporation

Electroporation was performed using Lonza 4D Nucleofector (V4XP-3032 for 20 μl as the manufacturer’s instructions). CD34+ HSPCs were thawed 24 h before electroporation. For 20 μl Nucleocuvette Strips, the RNP complex was prepared by mixing 3xNLS-SpCas9 protein^7^ (100 pmol) or HiFi-3xNLS-SpCas9 protein (100 pmol) and sgRNA (300 pmol, IDT) with glycerol (2% of final concentration, Sigma, G2025) and incubating for 15 min at room temperature immediately before electroporation. 50K HSPCs resuspended in 20 μl P3 solution were mixed with RNP and transferred to a cuvette for electroporation with program EO-100. The electroporated cells were resuspended with SCGM medium with cytokines and changed into EDM 24 h after electroporation.

#### Measurement of +58 BCL11A enhancer on-target and OT40 off-target indel and inversion

Editing frequencies were measured with cells cultured in EDM 5 days after electroporation. Briefly, genomic DNA was extracted using the Qiagen DNeasy Blood and Tissue kit (Qiagen, 69506). The *BCL11A* enhancer DHS +58 on-target site was amplified using forward primer AGAGAGCCTTCCGAAAGAGG (F1) and reverse primer GCCAGAAAAGAGATATGGCATC (R1). The off-target-rs114518452 site was amplified using forward primer TAAGATTCTTTTGGTTCTGGCT (F2) and reverse primer AGAGAGGCAGTATTTACGATGC (R2). The inversion junction was amplified using +58 forward primer (F1) and off-target-rs114518452 forward primer (F2), or +58 reverse primer (R1) and off-target-rs114518452 reverse primer (R2). KOD Hot Start DNA Polymerase (EMD-Millipore, 71086-31) was used for PCR and followed cycling conditions: 95 degrees for 3 min; 30 cycles of 95 degrees for 20 s, 60 degrees for 10 s, and 70 degrees for 10 s; 70 degrees for 5 min. 1 μl of locus specific PCR product was used for indexing PCR with KOD Hot Start DNA Polymerase and index primers following cycling conditions: 95 degrees for 3 min; 10 cycles of 95 degrees for 20 s, 60 degrees for 10 s, and 70 degrees for 10 s; 70 degrees for 5 min. Resulting PCR products were subjected to deep sequencing.

#### Amplicon deep sequencing and analysis

Amplicons were sequenced using paired-end 150 bp reads on an Illumina MiniSeq system with >18,000X coverage per sample for the off-target-rs114518452 site and >3,800X coverage per sample for the on-target site. Reads were trimmed for adapters and quality using Trimmomatic v0.36 in paired-end mode for the off-target-rs114518452 site and in single-end mode for the on-target site due to a nearby difficult-to-sequence homopolymer region. Editing outcomes were analyzed using CRISPResso v2.1.0 by aligning to the expected reference and/or alternative allele amplicons. A Needleman-Wunsch gap opening penalty of −30 (CRISPResso2 default: −20) was used to ensure more accurate alignment of reads to the reference vs. alternative allele amplicons for off-target-rs114518452 since they only differ by a single nucleotide. Only indels overlapping the expected SpCas9 cleavage site (3 bp upstream of the PAM) were counted as gene edits. The median observed indel frequency is reported for samples for which technical replicates were performed (n = 4), which includes all amplicon sequencing at the off-target-rs114518452 site for the donor heterozygous for rs114518452. Representative reads collapsed by allele identity and indel type are presented in the plots.

#### Inversion PCR

Nested PCR was performed to amplify the inversion junction. First step PCR was amplified using the outer primers on-target +58 forward, CACACGGCATGGCATACAAA, and off-target-rs114518452 forward, AATAGCCAAACTACTGAGCATTGTG; or the outer primers on-target +58 reverse, CACCCTGGAAAACAGCCTGA, and off-target-rs114518452 reverse, ACTAAGGCAATTGTTGTCCAAGC. KOD Hot Start DNA Polymerase was used for PCR and followed cycling conditions: 95 degrees for 3 min; 30 cycles of 95 degrees for 20 s, 60 degrees for 10 s, and 70 degrees for 10 s; 70 degrees for 5 min. 1 μl of PCR1 product was used for the second step PCR amplifying with inner primers on-target +58 forward (F1) and off-target-rs114518452 forward (F2), or on-target +58 reverse (R1) and off-target-rs114518452 reverse (R2) with cycling conditions: 95 degrees for 3 min; 10 cycles of 95 degrees for 20 s, 60 degrees for 10 s, and 70 degrees for 10 s; 70 degrees for 5 min. Resulting PCR products were loaded on a 2% agarose (VWR, 97062-250) gel. Images were captured by the BioRad ChemiDoc^TM^ MP Imaging System.

#### Droplet digital PCR

100 ng of gDNA was used for ddPCR with the ddPCR supermix (no dUTP, Bio-Rad, 1863024). See primer and probe sequences below. The premixed samples were placed into the Automated Droplet Generator (Bio-Rad, 1864101) that utilized Automated Droplet Generation Oil for Probes (Bio-Rad, 1864110) for droplet generation prior to PCR. The cycling conditions were: 1 cycle of 95°C for 10 min, 50 cycles of 94°C for 1 min sec (2°C/s ramp) and 56°C for 1 min (2°C/s ramp), 1 cycle of 98°C for 10 min, hold at 4°C. After thermal cycling, plate was placed in the QX200 Droplet Reader and plate layout set-up using QuantaSoft Software (Bio-Rad, 10031906).

**Supplementary Table 5.**
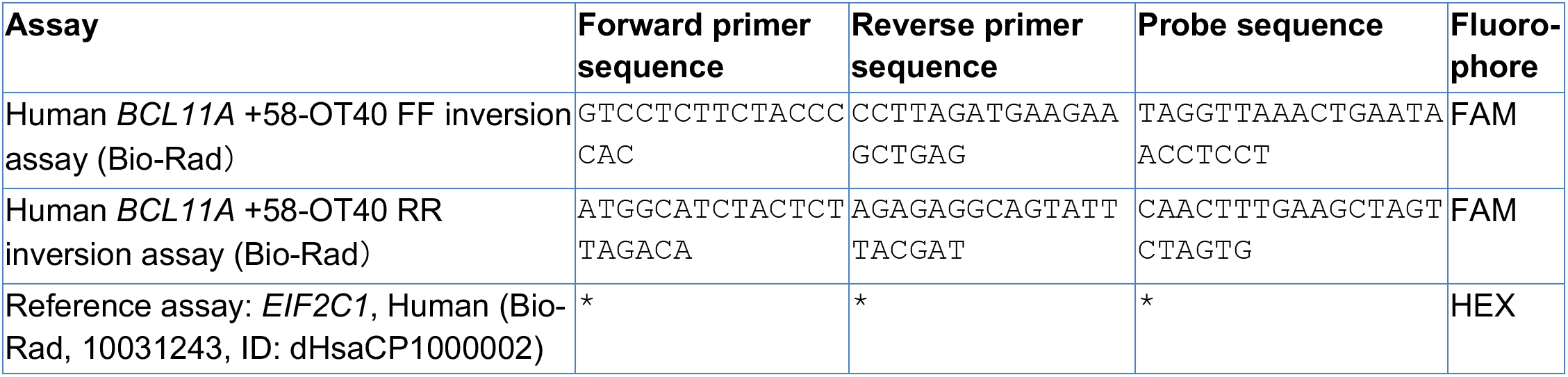
Assays for ddPCR. \**EIF2C1* reference assay is proprietary to Bio-Rad; the context sequence is hg19|chr1:36359339-36359461:+ GGGCGCGAGGTCTGGTTCGGCTTTCACCAGTCTGTGCGCCCTGCCATGTGGAAGATGATGCTCAACATTGATGGTGAGTGGGGAGAGCTATGGAGCCAGGGGCACCCCAAGTCCAGTGACCAC.

## Supplementary Files

Supplementary File 1. Top candidate off-targets in CRISPRme search results for sg1617 using hg38, 1000G, and HGDP data with up to 6 mismatches and 2 bulges (including the integrated_results, all_results_with_alternative_alignments, and private_targets files).

Supplementary File 2. Top candidate off-targets in CRISPRme search results for other example gRNAs with NGG PAMs.

Supplementary File 3. Top candidate off-targets in CRISPRme search results for example gRNAs with non-NGG PAMs.

Supplementary File 4. Top candidate off-targets in CRISPRme search results for example non-CRISPR based, RNA-targeting strategies (ASO and RNAi).

## Notes

http://crisprme.di.univr.it

### Summary of Updates

Improved software, new analyses.

